# Species-specific cutaneous protein signatures upon incision injury and correlation with distinct pain-related phenotypes in humans

**DOI:** 10.1101/2022.03.07.482980

**Authors:** Daniel Segelcke, Max van der Burgt, Christin Kappert, Daniela Schmidt-Garcia, Julia R. Sondermann, Stephan Bigalke, Bruno Pradier, David Gomez-Varela, Peter K. Zahn, Manuela Schmidt, Esther M. Pogatzki-Zahn

**Affiliations:** Department of Anaesthesiology, Intensive Care and Pain Medicine, University Hospital Muenster, Muenster, Germany; Max Planck Institute of Experimental Medicine, Goettingen, Germany; Clinic for Anaesthesiology, Intensive and Pain Medicine, Ruhr-University Bochum, BG-University Hospital Bergmannsheil gGmbH, Bochum, Germany; Division of Pharmacology and Toxicology, Department of Pharmaceutical Sciences, University of Vienna, Vienna, Austria

## Abstract

Pain after surgery is common, and its management remains a clinical challenge. Severe acute and prolonged post-surgical pain impairs immediate recovery and leads to long-term consequences like chronic pain, opioid dependency, and reduced quality of life. Althought rodent pain incision models exist, translation to patients is still hampered. To bridge this gap, we combined sensory phenotyping with quantitative proteomics and protein networks in humans and mice after skin incision representing an established model for surgical pain. Initially, we revealed, for the first time, similarities and differences of protein-protein interaction (PPI) networks across both species. Next, we comprehensively phenotyped humans for pain-related symptoms and observed phenotypes with incision-induced proteome changes. Remarkably, post-incision PPI-networks differed between volunteers with small incision-related hyperalgesic areas (“Low responder”) versus those with large areas (“High responder”). The latter exhibited a pronounced proteolytic environment associated with persistent inflammation, while an anti-inflammatory protein signature was observed in Low responders. Taken together, we provide unprecedented insights into peripheral processes relevant for developing hyperalgesia and pain after incision. This knowledge will immensely facilitate bidirectional translational pain studies and guide future research on the pathophysiology of pain after surgery and the discovery of novel targets for its treatment and prevention.

## Introduction

Worldwide more than 300 million people undergo surgery each year. Most of them experience acute pain, and treatment remains inadequate in a very high number of patients (1,2). Severe post-surgical pain not only leads to the suffering of patients after surgery; it hampers patients’ recovery by impairing mobilization and physiotherapy, increases complications, and delays discharge after surgery (3,4). Even more challenging are pain-related long-term consequences, including severe complications, long-term opioid intake, and chronic post-surgical pain (CPSP) (1–3,5–8). These consequences affect patients’ quality of life and have a tremendous economic burden for patients as well as the community explaining why chronic post-surgical pain will be for the first time included in the new ICD-11 classification (5). Due to insufficient knowledge about the multifaceted processes leading to acute and chronic pain after surgery, the current therapeutic options are sparse. For severe post-surgical pain, opioids and regional anesthesia techniques are most effective, but both are limited by side effects, risks, and long-term consequences (9). Pure systemic non-opioid analgesic regimens are therefore affordable, but most available drugs are of low efficacy and associated with significant adverse events and therefore not recommended for the treatment of post-surgical pain (10,11). To prevent chronic post-surgical pain, the efficacy of available drugs is limited, and their clinical relevance is rather uncertain (12). Together, there is an urgent need for novel, effective, and safe non-opioid analgesics to treat acute and prevent chronic pain after surgery.

For the last 20 years, specific rat and mouse models of incisional pain have been available and are increasingly used to uncover the mechanisms inherent in acute pain after surgery (13–17). These models are highly relevant as they provide tremendous insights into the unique pathophysiology of post-surgical pain (15). However, almost none of these studies have yet found their way into clinical practice – a common problem in pain research. Reasons here are manifold, culminating in limited translatability from rodents to human patients (18,19). Although species-specific differences exist, preclinical findings in rodents were only rarely validated in human patients, and if so, species-specific differences related to the nociceptive system were noticeable (19,20). One missing link in cross-species translation could be human pain models mimicking the tissue injury and aspects of pathological pain usually seen in patients (21). A human incision model has been developed for pain after surgery that shows symptoms similar to those of surgical patients, including acute pain and hyperalgesia (increased sensitivity) surrounding the injured tissue (22,23). Especially mechanical hyperalgesia has been shown to serve as a clinically relevant pain phenotype post-surgically; as indicated years ago, increased size of mechanical hyperalgesia surrounding the surgical wound was associated with chronic pain several months after abdominal surgery (24). Subsequent clinical studies supported this association (25–27). However, the pathophysiology underlying mechanical hyperalgesia after surgical incision has rarely been investigated in humans.

Another problem inherent to preclinical pain research is a narrow, biased approach; most studies investigating pain focus on single aspects, such as RNAs, receptors, and others, without pursuing a systems biology approach. As indicated in other disciplines such as oncology, understanding the system biology level is key to understanding complex disease-related processes (28–32). After incision injury, a multifaceted recovery process is initiated in the skin that proceeds in a highly orchestrated manner to protect the tissue from further harm and heal epithelial damage. Therefore, investigating single candidate proteins in this precisely balanced process at the level of protein-protein interaction (PPI) networks is not sufficient to understand the mechanistic underpinnings. In contrast to transcriptomics (i.e., studies on the mRNA level), quantitative proteomics enables an unbiased analysis of PPI networks and their dynamics during pathological conditions as mRNA levels only show limited correlation with protein abundance (33–37). Consequently, the analysis of PPI-networks, rather than single aspects, parallel studies in rodents and humans, and the combination of molecular data with pain phenotyping are mandatory for deciphering mechanisms relevant to hyperalgesia in patients.

We recently demonstrated the potential of quantitative proteomics by revealing a distinct PPI network in dorsal root ganglia of mice upon incision injury (38). To uncover relevant PPI for post-surgical pain in humans, we used here a parallel, unbiased, and quantitative proteomics approach combined with behavior and phenotype analysis in humans and mice. First, we quantified proteome changes in skin biopsies derived from humans and mice after incision and subsequently analyzed incision-modulated PPI networks. In this way, we revealed, for the first time, similarities and differences across both species after a comparable incision injury. Next, the determination of individual human volunteer psychophysics pre- and post-incision enabled detailed phenotyping of pain-related symptoms and a correlation of phenotypes with PPI-network dynamics. Remarkably, post-incision PPI-networks differed between volunteers with small incision-related hyperalgesic areas (termed “Low responder”) versus those with large areas (termed “High responder”). Taken together, we report here unprecedented insights into incision-induced proteome dynamics in skin biopsies: (1) a hitherto unknown and species-specific protein signature in humans and mice post-incision and (2) PPI-network changes of primary incisional wounds that correlate with distinct pain-related phenotypes. Thus, our study significantly increases our knowledge on species-specific alterations in the incision model of post-surgery pain with high translational impact for clinical research.

## Results

### Incision injury alters the protein signature in human and mouse skin with distinct species characteristics

We performed a complex integrated workflow to identify – in an unbiased manner - protein signatures in humans and mice in well-established human and mouse models of skin incision (39) (Fig. 1). In human volunteers, we induced a skin incision in one forearm followed by the assessment of incision-related symptoms, such as the size of the area of secondary hyperalgesic (time points: 1 and 24 h post-incision), acute incisional pain (during and after incision), and mechanical thresholds around the incision (time point: 1 and 24h post incision). Multilevel baseline (BL) characterization of each volunteer was performed to ensure that inclusion criteria were fulfilled (table S1) and that previous psychiatric diseases or limitations of sensory perception were absent. Extensive psychophysical testing, including a set of psychological questionnaires (table S2) and complete quantitative sensory testing (QST, fig. S1A, table S3 for raw data) on the volar forearm was used; no volunteer needed to be excluded from the study (for volunteer demographics, see Fig.1). At 24 h post-incision and after final psychophysical testing, we obtained skin biopsies from the ipsi- (incised; biopsy was taken from “within” the incision) and contralateral (control) forearm and submitted those to quantitative proteome analysis by data-independent acquisition mass spectrometry (DIA-MS). The time point of 24 h post-incision was chosen due to the well-known maximum peripheral and central sensitization at that time in this experimental incision model (22,39).

**Fig. 1.**
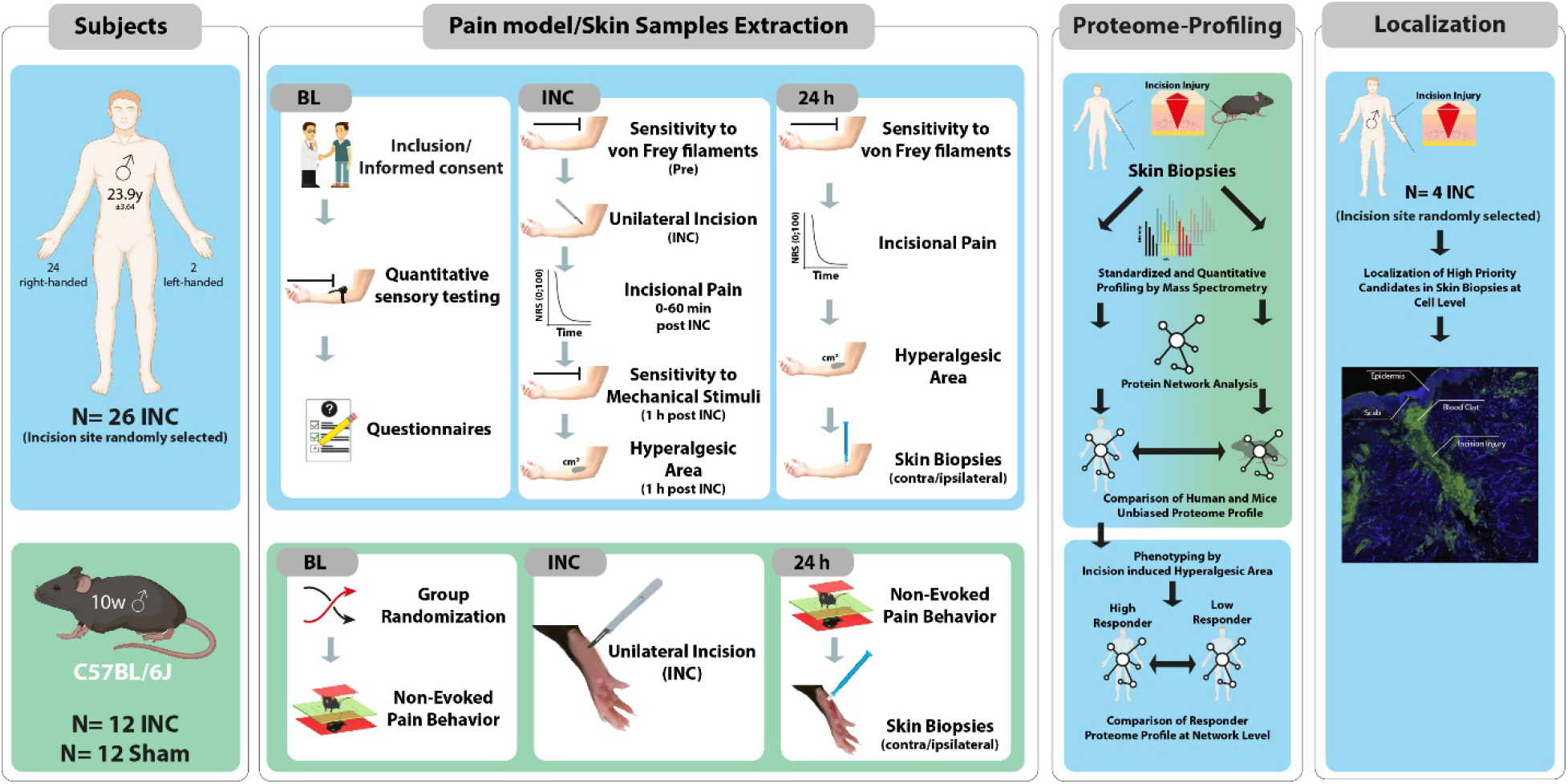
Study design. The study contains four major parts, (1) psycho-physics characterization of human volunteers (blue), behavior tests in mice (green), (2) experimental pain model induction and skin sampling, (3) proteome-profiling of incised skin from humans and mice, and (4) immunohistochemistry of high priority candidates in human incised skin. In total, 30 human male volunteers (26 right-handed) with a mean age of 23.9 years (y) were included. Mouse experiments were performed with 12 incision- (INC), and 12 sham-treated C57BL/6J male mice (10 weeks old). Experiments were performed at three different time points: one day prior incision (baseline, BL), at incision (INC) day, and 24 h post-incision. Skin biopsies (ipsi- and contralateral), including epidermis and fascia, were analyzed by quantitative mass spectrometry. Protein-protein interaction network analyses were performed across both species and pain phenotyping groups in humans. Localization of priority candidates was examined in incised skin samples 24h post-incision.

Upon incision on the ipsilateral volar forearm, incisional pain peaked right after, decreased rapidly within the first 10 minutes, and was abolished entirely 24 h post-incision (figure S1B). In particular, mechanical pain thresholds were decreased (MPT, figure S1C), and mechanical pain sensitivity (MPS, figure S1D) increased post-incision on the ipsilateral site near the incision, suggesting the development of primary mechanical hyperalgesia. In addition, a secondary mechanical hyperalgesic area (HA) developed around the incision in the unaffected skin in a time-dependent and individual manner (Fig 5A and figure S4A). These results are in line with previous reports using this experimental incision model in human volunteers (22,39).

Quantitative proteomics of 21 humans (exclusion of 5 volunteers, see methods and table S4) skin biopsies (B) from “within” the incision (B_inc_) and in the corresponding area of the contralateral non-incised volar forearm (B_con_) of each volunteer resulted in a highly reproducible list of 1569 quantified proteins in each sample (for a complete dataset, please see table S6). Unsupervised hierarchical clustering (Fig. 2A) and principal component analysis (PCA) (Fig. 2B) clearly showed the expected separation of proteome signatures of B_inc_ compared to B_con_. Upon statistical comparison (corrected for multiple testing using the Benjamini-Hochberg (BH) method to obtain a q-value (40–42); table S6), we observed pronounced overall proteome changes as visualized in the volcano plot (Fig. 2C). Specifically, 119 out of 1569 quantified proteins (i.e., ∼7.6% of all quantified proteins, in bold font in table S6) were significantly and robustly (log_2_ FC > |0.38|) regulated upon incision. Among these, 73 proteins were up-and 46 downregulated and predicted to be localized in diverse cellular components (GO_CC) by gene ontology (GO) analysis, e.g., secretory granules, fibrinogen complex, and phagocytic cup (for the whole list, please see figure S2A).

**Fig 2.**
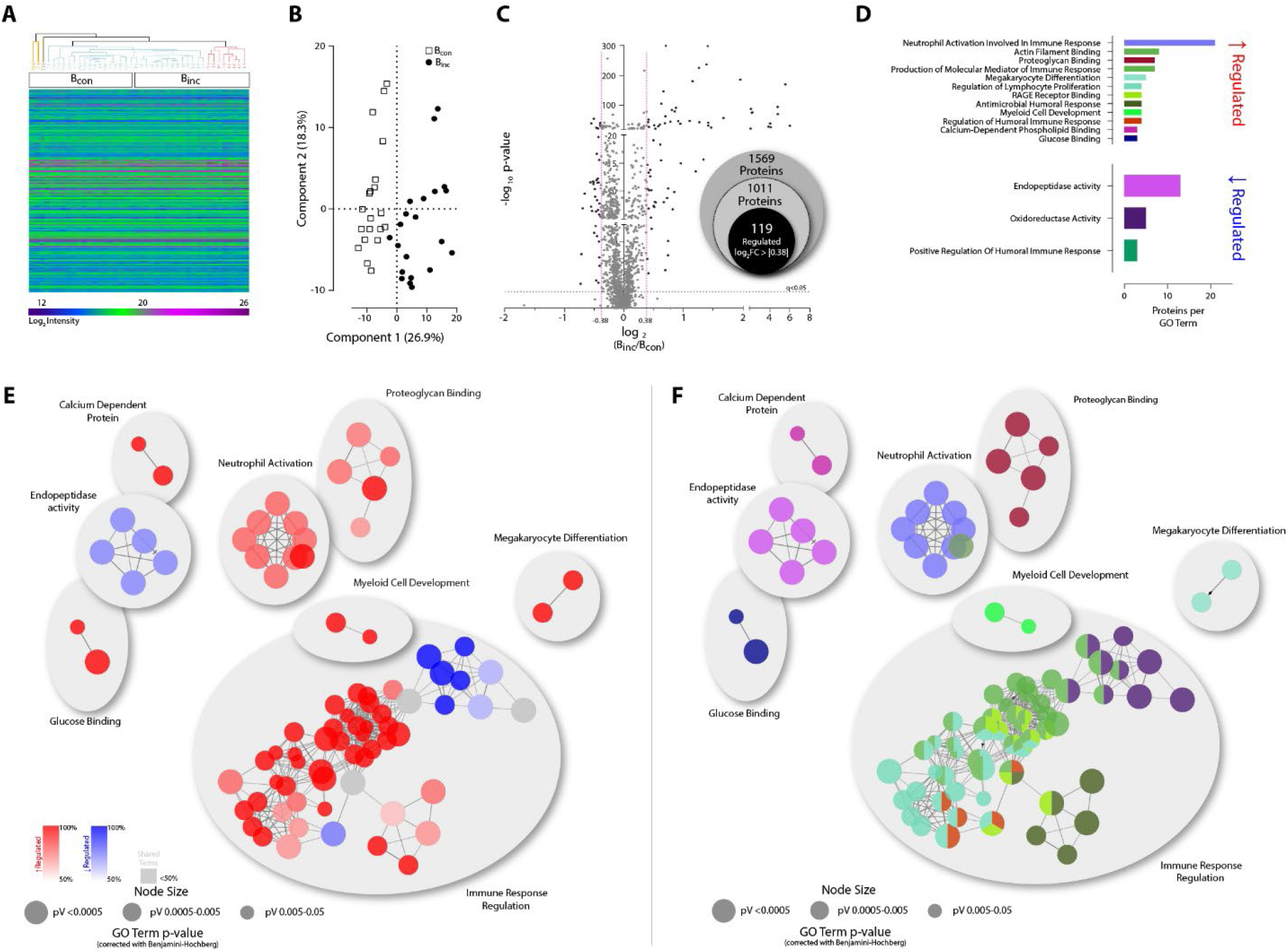
Incision injury alters unbiased protein profiling in human skin. (A) Unsupervised hierarchical clustering of Log_2_ protein intensities obtained by DIA-MS across all skin biopsies (contralateral (B_con_) and ipsilateral (B_ipsi_)). The legend color bar indicates the range of Log_2_ intensity within rows. (B) Principal component analysis separated the proteome signatures of skin biopsies (B) with incision (B_inc_) compared to the contralateral site (B_con_) with component 1 by 26.9% and component 2 by 18.3%. (C) In total, 1569 proteins (grey-colored) were quantified, among which 119 proteins (∼7.6 % of all quantified proteins, black colored) were regulated log_2_FC > |0.38|. Proteins considered as significantly regulated if they exhibited q < 0.05 (horizontal dotted line) and thresholded by log_2_FC > |0.38| (pink vertical dotted line) (D) Gene Ontology (GO) pathway (GO_molecular function AND GO_Immun system process analysis of regulated (upregulated in red; down-regulated in blue) proteins in ClueGO (v2.5.8). Only pathways are shown pV≤ 0.05. (E, F) Enriched term-term interaction (TTI) network analysis of regulated proteins was functionally grouped by ClueGO (v2.5.8) as functional clusters (AutoAnnonate 1.3). Each node represents a molecular function or immune system process. (E) Node colors associate with protein regulation (upregulated in red; down-regulated in blue) (F) or their function (description of the term see in D). Gray nodes contain proteins from both regulatory lists and therefore cannot be unambiguously assigned. TTI represents by edges. The size of nodes reflects the significant enrichment of the terms.

Regulated proteins were associated with distinct gene ontology terms for molecular function (GO_MF) and immune system processes (GO_ISP) such as *neutrophil activation in immune response*, *actin filament binding*, *endopeptidase activity*, and *oxidoreductase activity* (Fig. 2D). Accordingly, hub-related GO enrichment analysis (Fig. 2E, F) highlighted eight major clusters. Among them, the cluster *immune response regulation* was most prominent and included the GO-MF terms like *regulation of lymphocyte proliferation* (cyan), *oxidoreductase* (dark purple), and *antimicrobial humoral response* (dark green), to name a few (Fig. 2D and 2F). While this cluster was composed of up - (red) and down (blue)-regulated hubs (Fig. 2E), other clusters contained mainly down-regulated proteins (*endopeptidase activity*) or mainly upregulated proteins (all other clusters, e.g., *neutrophil activation*). In conclusion, our proteomics and GO-based analysis upon skin incision in human volunteers suggest significant alterations of pathways associated with the healing processes of incisional wounds and associated peripheral sensitization processes, resulting in pain and hyperalgesia.

Despite well-known limitations, translational pain research heavily relies on rodent models of painful diseases (18). In addition, mouse models are indispensable for back-translational research. In the context of incisional pain, the mouse model of unilateral hindpaw plantar incision injury is well-established (16,38). We employed this model to critically assess similarities and differences between humans and mice upon incision. We assessed non-evoked pain-related behavior at rest (figure S3), harvested punch biopsies of the incised skin (B_inc_M; M, Mouse), and sham-treated plantar aspect (B_sham_M), and analyzed relative proteome abundance changes in B_inc_M compared to B_sham_M similar to the way described for human volunteers above. In total, we measured samples from 12 mice/condition, of which three were pooled in one biological replicate (i.e., 3 biological replicates per condition, B_inc_M and B_sham_M, table S5). Unsupervised hierarchical clustering (Fig.3A) and PCA (Fig. 3B) clearly show the expected separation of proteome signatures of incision compared to sham-treated mice. In each sample, we quantified 1871 proteins. Upon statistical comparison (corrected for multiple testing using the Benjamini-Hochberg (BH) method, q-value (40–42)), we identified 435 proteins with significant and robust (log_2_ FC > |0.38|) abundance changes compared to sham (i.e., ∼23.3% of all quantified proteins, in bold font in table S7), with 175 being upregulated and 260 being downregulated (Fig. 3C and table S7). Upregulated proteins were predicted to be localized to myofill, mitochondrial envelope, inner membrane, or intermediate filament (for the whole list, please see figure S2B, red). Smooth endoplasmatic reticulum, cortical cytoskeleton, actin filament, or nucleosome are the most prominent sites to which downregulated proteins were predicted to be localized (for the whole list, please see figure S2B, blue). Functionally distinct GO_MF and GO_ISP were observed among regulated proteins, e.g., *negative regulation of endopeptidase activity, complement activation*, *regulation of ATPase activity*, and *negative kinase activity* (Fig. 3D). Hub-related GO enrichment analysis revealed 22 clusters (hubs without clustering were excluded), among which *complement activation* was highly prominent and mainly contained upregulated hubs (Fig. 3E, F).

**Fig 3.**
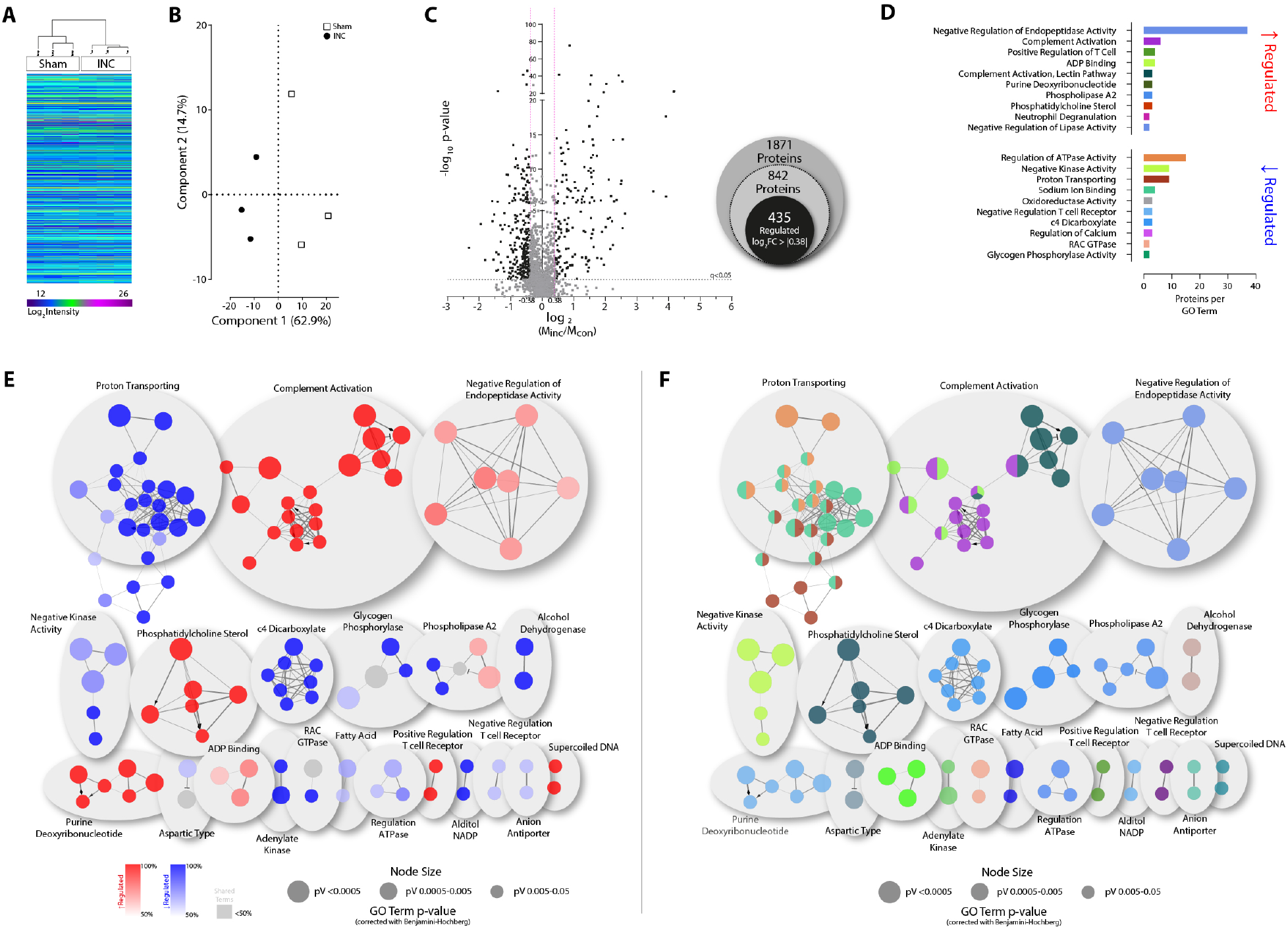
Unbiased protein profiling in mice skin modulates by incision injury. (A) Unsupervised hierarchical clustering of Log_2_ protein intensities obtained by quantitative DIA-MS of paw skin biopsies (B) from mice three biological replicates per condition, N=9 Sham, N=9 Incision, INC). The legend color bar indicates the range of Log_2_ intensity within rows. (B) Principal component analysis separated the proteome signatures of skin biopsies (B) with incision (B_inc_) compared to the contralateral site (B_con_) with component 1 by 62.9% and component 2 by 14.7%. (C) In total, 1871 proteins (grey-colored) were quantified, among which 435 proteins (∼23.3 % of all quantified proteins, black colored) were regulated more than log_2_FC > |0.38|. Proteins were considered as significantly regulated if they exhibited q < 0.05 (horizontal dotted line) and thresholded by log_2_ FC > |0.38| (pink vertical dotted line) (D) Gene Ontology (GO) pathway (GO_molecular function AND GO_Immun system process analysis of regulated (upregulated in red; down-regulated in blue) proteins in ClueGO (v2.5.8). Only pathways are shown pV≤ 0.05. (E, F) Enriched term-term interaction (TTI) network analysis of regulated proteins was functionally grouped by ClueGO (v2.5.8) as functional clusters (AutoAnnonate 1.3). Significantly (*p*-values ≤ 0.05) enriched GO terms visualized in a functionally grouped network that reflects the terms’ relationships. Each node represents a molecular function or immune system process. (E) Node colors were associated with protein regulation (upregulated in red; down-regulated in blue) (F) or their function. The degree of connectivity between terms (kappa score) was used for defining functional groups (highlighted in color see in D). Gray nodes contain proteins from both regulatory lists and therefore cannot be unambiguously assigned. TTI is represented by edges. The size of nodes reflects the significant enrichment of the terms.

Next, we aimed at assessing similarities and differences between protein signatures and their incision-induced dynamics in human and mouse skin (Fig. 4). This direct comparison is particularly relevant for translational pain research as previous studies have shown major differences in cutaneous repair mechanisms after incision injury between humans and mice (43). Overall, human and mouse skin proteomes showed an extensive compositional similarity, i.e., 1159 proteins were identified in both species (Fig. 4A). On the other hand, 712 mouse proteins (i.e., 38.1% of the quantified mouse skin proteome) were not reliably detected in all human volunteers and vice versa for 410 human proteins (i.e., 26.1% of the quantified human skin biopsy proteome, Fig. 4A). Note, however, that these discrepancies do not indicate the absence of respective proteins from one species. Given our stringent detection and quantification criteria (see above and methods for details), here reported species comparisons solely take those proteins into account that were quantified across all human (Fig. 2) or mouse skin biopsies (Fig. 3), respectively. Upon incision, we could discern 50 commonly regulated proteins in both species (Fig. 4B), amounting to 61% of all regulated human and 10% of all regulated mouse proteins. Notably, the direction of the regulation upon incision, i.e., up-or downregulated, was highly conserved in humans and mice with only a few exceptions such as ARPC3, AZGP1, COL1A1, SERPINB8, and HPX (Fig. 4C). Additionally, the top ten Reactome pathways, which were regulated upon incision, were largely comparable across species with few exceptions, e.g., *post-translational protein modification*, *metabolism of proteins*, and *metabolism of lipids*. Human skin biopsies exhibited noticeable changes in proteins annotated to *extracellular matrix organization*, a relatively underrepresented pathway in mouse skin biopsies (Fig. 4D). Vice versa, *post-translational protein modification,* and *metabolism of proteins* were enriched in mouse skin biopsies but not in human skin. We then used the web-based STRING interface (www.string-db.org, (44)) to visualize predicted species-dependent protein-protein interaction networks (PPI) across the 50 overlapping proteins (Fig. 4E). *Platelet aggregation*, *inflammatory activity*, and *actin polymerization* represent clusters that were prominently altered in both species. Annotated upregulated proteins, such as FN1, FERMT3, and LCP1, exhibited a high degree of connectivity and betweenness centrality, indicating that these proteins constitute to highly connected hubs in both species (Fig 4E, bold font). These data represent the first parallel comparison of post-incision protein alterations in human and mouse skin and reveal species-specific and overlapping changes in protein abundance. In particular, the knowledge of hitherto unknown disparities between human and mouse signatures upon incision is essential for successful bidirectional-translational research on the widely-used model for post-surgical pain across species.

**Fig 4.**
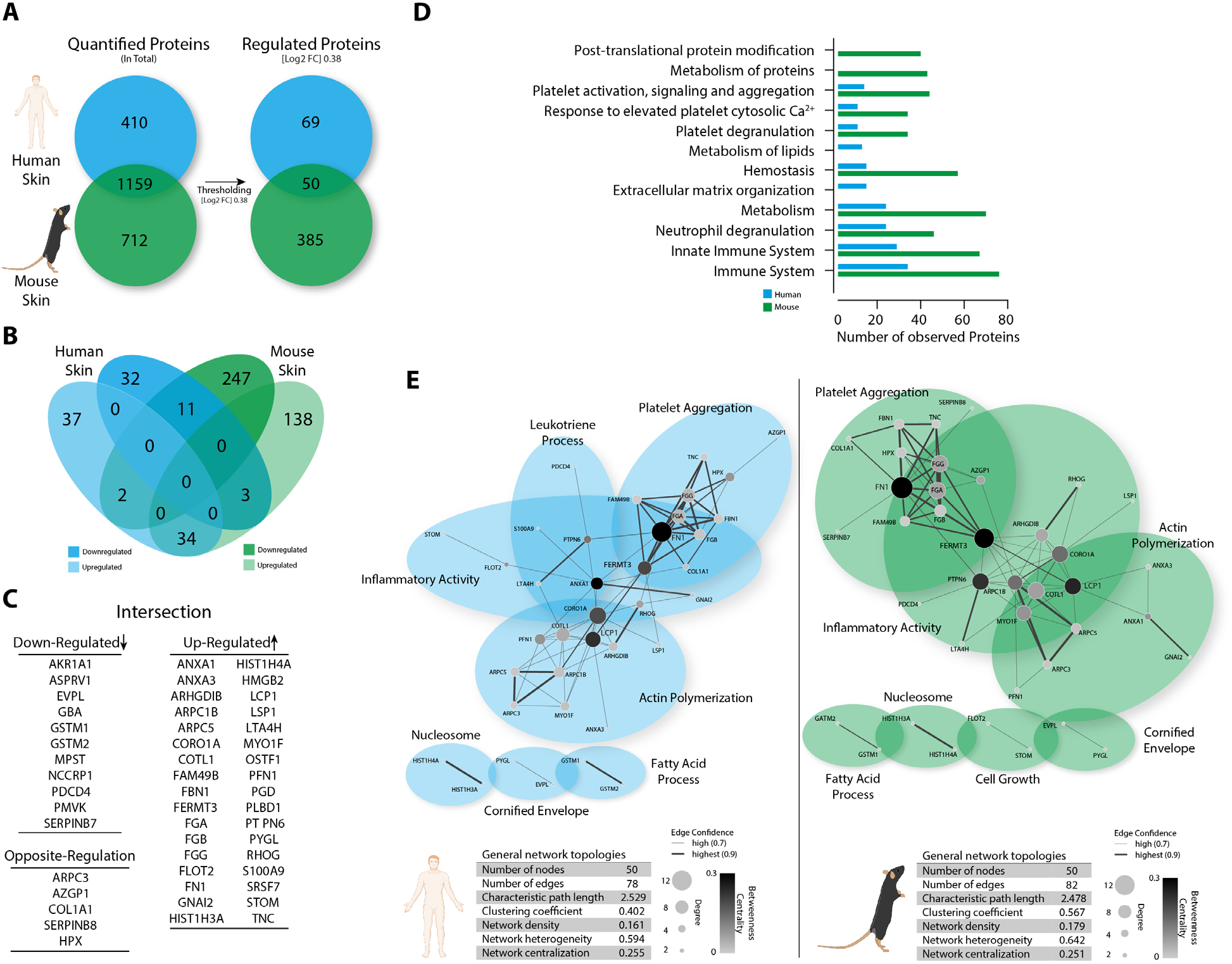
Incisional injury mediates distinct cutaneous protein signatures with a small amount of overlap in human and mouse biopsies. (A) After thresholding, 69 proteins were uniquely quantified in humans and 385 proteins in mice, while 50 proteins were overlapping in both species. (B) Venn diagrams display the detailed comparison of human and mice incision-induced protein signature regarding their regulation. Proteins regulated in both species (intersections) are listed in C. (C) List of incision-induced species overlapping 50 proteins regarding their regulation. (D) Gene Ontology (GO) pathway and REACTOME analysis revealed annotations of both species (human: blue, mice: green). Only the Top-Ten pathways, which exhibit significant enrichment (FDR < 0.05; assessed via the web-based interface STRING), display. (E) Enriched protein-protein interaction (PPI) network of incision-induced species overlapping skin proteins (FDR < 0.05; assessed via the web-based interface STRING). Network topology was analyzed by stringAPP in Cytoscape (3.8.2). Each node represents a single protein, including degree (size) and betweenness centrality (color). PPI is reflected by edges (thickness represents the confidence). Functional grouping was performed with AutoAnnonate 1.3.

### Pain-related phenotyping reveals human responder-types with distinct PPI signatures

While a comparison of molecular datasets from human and mouse incision models – as shown above- is mandatory for successful bidirectional translational research in the context of post-surgical pain, it may not sufficiently represent the diversity of human pain-related phenotypes. One way to overcome these limitations is to use, whenever ethically possible, human experimental models (18) in a so-called microtranslational approach. Microtranslation allows for identifying and investigating molecular mechanisms underlying distinct human phenotypes. Akin to patients after surgery, volunteers were undergoing the here-used skin incision model exhibit a wide variety of pain-related symptoms from acute pain right after an incision to developing hypersensitivity to mechanical or heat stimuli at the site of damage (primary hyperalgesia) and in the unaffected surrounding area (secondary mechanical hyperalgesia (45)) figure S1, figure S4A). Thus, we set out to test whether incision-related symptoms at given time points post-incision could be correlated with obtained snapshots of incision-induced alterations of the protein signature. Given that the size of the area of secondary mechanical hyperalgesia (HA) is considered a risk factor for the development of chronic pain after surgery (24), we then used this symptom to group human volunteers. We categorized volunteers according to the size of the HA, both 1 h and 24 h post-incision (HA time course, HATC) (Fig. 5A) and, separately, 24 h after incision (HA24) (figure S4A). Both HA parameters enabled us to group volunteers into High and Low responder types. This cohort’s cut-off values for HATC categorization were explicitly determined using the mean value (HA1, mean 68.65, HA24, mean 45.04 cm^2^, Fig 5A) and HA24 categorization mean and SD (mean 45.04 cm^2^ ± 49.62, figure S4A). We then repeated the proteome data analysis separately for each responder type and applied our cut-off (as explained above, Fig. 2) to highlight pronounced protein regulations. A comparison of responder types revealed overlapping proteins (e.g., 51 for HATC, Fig.5B and 54 for HA24, figure S4B) and, interestingly, responder type-specific proteins uniquely regulated either in High or Low responders. Focusing on HATC data, GO pathway analysis identified terms only present in HATC High responders, such as *oxidoreductase* and *serine-type endopeptidase activity* (Fig. 5C). Among shared GO pathway terms across both responder types were *myeloid leukocyte mediated immunity* and *granulocyte migration/chemotaxis*, to name a few (Fig. 5C). Enriched GO pathway analysis on regulated proteins predicted ten biological clusters (referred to as “cluster” with names in bold font, Fig. 5D) with their associated proteins (protein names are given in regular font; Fig. 5D). Among clusters were *immunity regulation, regulation of macrophage chemotaxis*, *oxidoreductase activity*, and *antibacterial response* harboring diverse GO terms, specifically in High responders (magenta circles). Cluster-associated proteins regulated in High responders, such as ELANE, MMP9, CYBB, and ITGAM (bold font), exhibited high cluster-cluster or cluster-protein interactions represented by magenta edges. Enhanced MMP9 or serine type-protease (CTSG, ELANE, LTF, and PRTN3) levels suggest a prevalent proteolytic environment and dysregulation upon incision, being in line with previous reports on chronic wounds associated with pain (46,47). While MMP9 and ELANE were prominently regulated in High responders (full magenta circles), CTSG, LTF, and PRNT3 were also regulated in HATC Low responders (magenta/cyan circles). Taken together, these data highlight previously unknown similarities and disparities between responder types based on pain-phenotyping both on the pathway and protein level.

**Fig 5.**
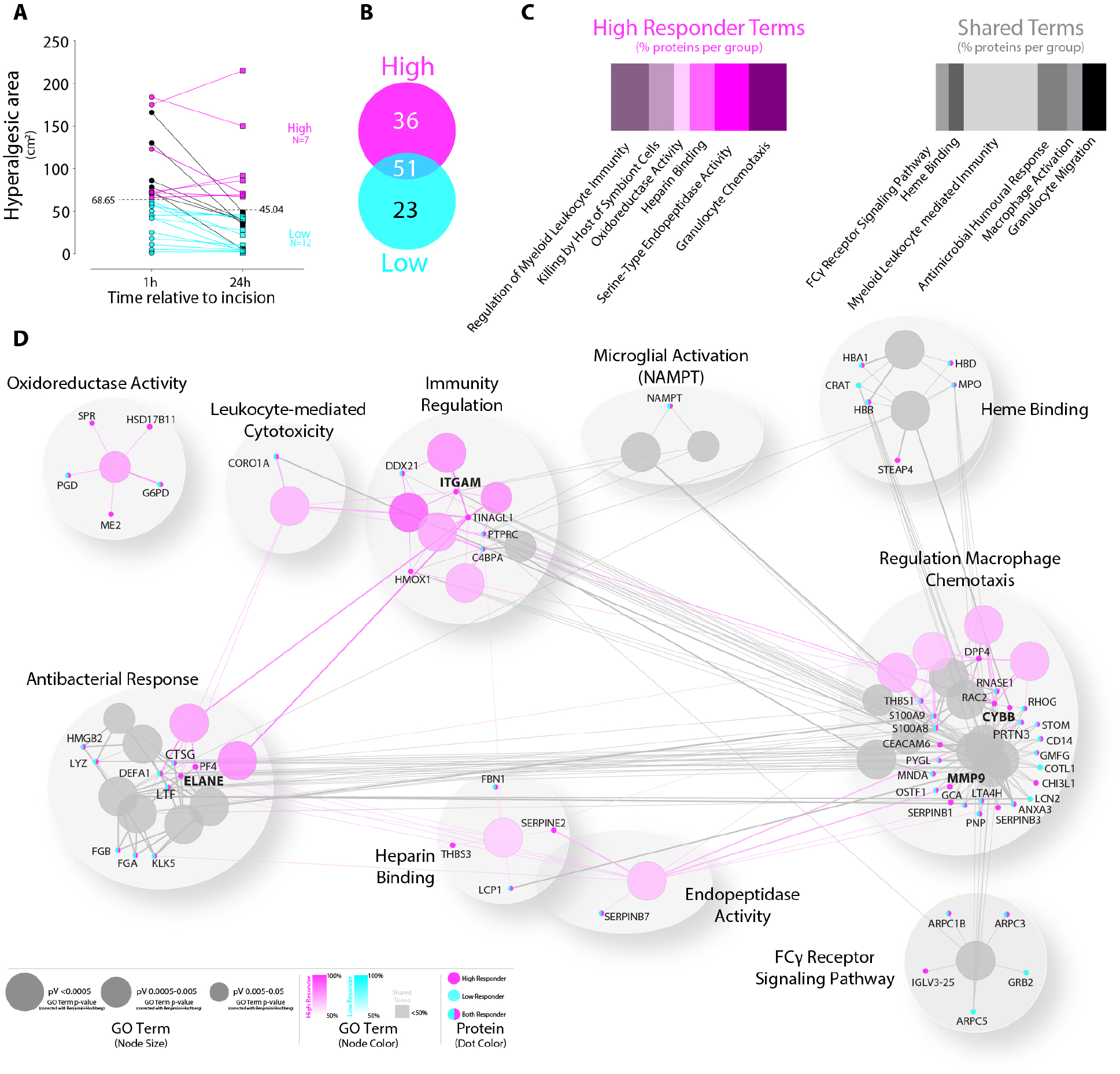
Dominance of High-responder enriched term-term interaction networks post-incision. (A) After a skin incision, a hyperalgesic area developed around the wound over time as expected. The total human cohort was phenotyped into High (n=7, magenta), Low Responder (n=12, cyan), and Undefined volunteers (n=7, black). Dimension of hyperalgesic area (cm^2^) at 1 and 24h post-incision (HATC) expressed as individual values connected by a line. HATC categorization was determined using the mean value (HA1, mean 68.65, HA24, mean 45.04 cm^2^, black dotted lines) (B) Thirty-six proteins that were uniquely regulated in High responders, 23 proteins for Low-responders, and 51 overlapping proteins were identified. (C) GO-analysis revealed annotations of regulated proteins for High Responder (magenta shades). Shared terms belong to both phenotyping groups (grey shades). In contrast, regulated proteins of the Low responders could not be significantly annotated to distinct pathways. (D) Enriched term-term interaction (TTI) network analysis with associated proteins was functionally grouped by ClueGO (v2.5.8) as functional clusters (AutoAnnonate 1.3). Significantly (*p*-values ≤ 0.05) enriched GO terms visualized in a functionally grouped network that reflects the terms’ relationships. Each node represents a molecular function or immune system process—node colours are associated with phenotyping (High Responder in magenta shades; Low Responder in cyan shades). Grey shades reflect unspecific terms, which belongs to both responder types. Proteins display as dots in responder colors (magenta= High Responder, cyan= Low Responder, color shared= responder independent). The degree of connectivity between terms (kappa score= 0.3) was used to define functional groups. TTI represents edges between nodes and dots. The size of nodes reflects the significant enrichment of the terms.

Next, we performed STRING PPI-topology-driven analysis to define responder type-specific network signatures and describe the role of each protein in the network. Those network signatures might indicate candidate proteins associated with or defining each responder type upon incision. For this purpose, merged networks from HATC and HA24 were used (Fig.6, figure S5). PPI architecture was analyzed by their topology features and diverse protein attributes (e.g., connectivity, characteristic path length). The High responder network contained seven highly connected (degree > 8) nodes (ELANE, LYZ, MMP9, MPO, ITGAM, CTSG, MNDA, magenta font) compared to only one such node in Low-responder networks (LYZ, cyan font) (Fig. 6A). The greater degree in conectivity in High responders is in accordance with differences in network density, number of edges, and, partially, with high betweenness centrality, particularly in the bottleneck hubs MMP9 and ITGAM. In High responders, serine-type endopeptidases (CTSG, LTF, PRTN3, magenta font in Fig. 6A) show a higher degree and shorter path length than in Low responders, suggesting a phenotype-specific role as part of the proteolytic environment (see above). This environment is substantiated when only the differential part of each responder type network is considered (Fig. 6B), i.e., when only responder type-specific PPI are plotted. As aforementioned, MMP9 represents the central hub for the PPI network in High responders. In addition to the proteolytic environment, other proteins, such as RAC2, NCF2, and CYBB (part of NADPH oxidase) (48), suggest alterations of the redox environment at the incision site. This may indicate elevated respiratory levels in phagocytotic cells and, consequently, increased reactive oxygen species (ROS) levels, likely promoting inflammatory responses. In contrast, the PPI network for Low responders (Fig. 6B) exhibited only a few nodes and edges (with decreased characteristic path length). These were related to anti-inflammatory functions (ALOX15B, PRTN3, LCN2 (49,50)) and adapter protein collagen deposition (GNB1, GRB2, COL3A1 (51)). While studies describing the function of the proteins ELANE and DPT in the cutaneous wound (secondary, excisional) healing could be found, these proteins have never been reported in the context of incisional wounds or pain symptoms. An exception is MMP9 which has been implicated in post-incision pain in the spinal cord (52,53), DRG and skin of mice (53).

**Fig. 6.**
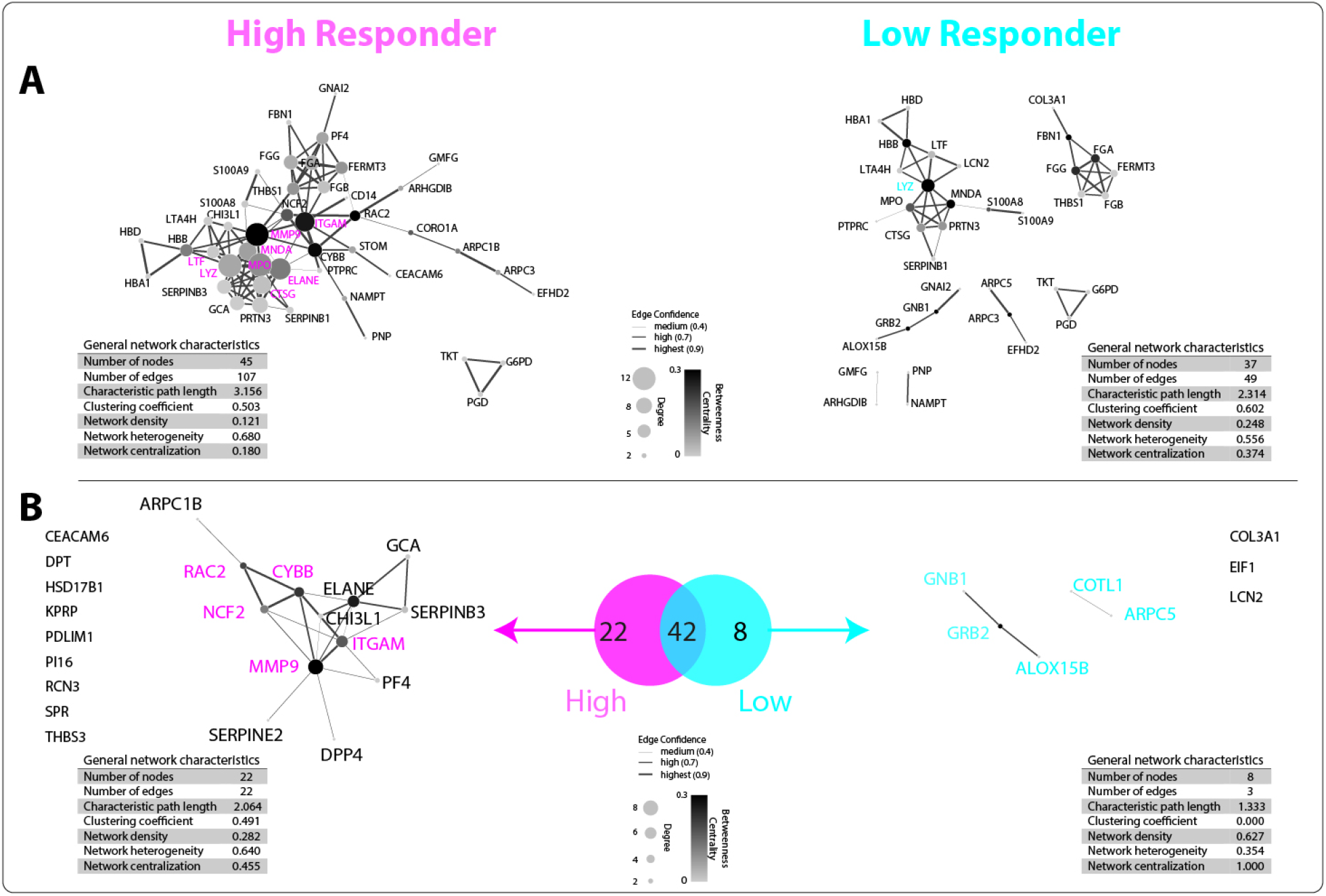
Skin incision induces distinct phenotypes protein-protein interactions network topology. (A) Enriched protein-protein interaction (PPI) networks for responder phenotypes post-incision (magenta, High Responder; cyan, Low Responder) resulting from merged networks (S5). (B) Differential PPI networks between High (magenta) and Low (cyan) Responder phenotypes. Unique PPI networks are displayed. Unlinked proteins are shown as names only, without a sphere. Enriched PPI network topologies of incision-induced phenotypes were analyzed (FDR < 0.05; assessed via the web-based interface STRING) by stringAPP in Cytoscape (3.8.2). Nodes represent a single protein, including degree (size) and betweenness centrality (color). PPI is reflected by edges (thickness represents the confidence).

These results suggest pronounced differences among responder types: While Low responder PPI signatures can be annotated to anti-inflammatory and cell migration processes, PPI signatures in High responders may be indicative for a proteolytic environment, associated with an imbalance of extracellular matrix (ECM) deposition and degradation, resulting in a hypercatabolic prolonged inflammatory state associated with an enlarged hyperalgesic area around the incision

Based on our data (Fig. 6), MMP9 and ELANE appear to play a central role in incision-induced protein networks involved in wound healing and peripheral sensitization processes. This is why we set out to study their localization in skin biopsies by immunohistochemistry. Upon incision, MMP9 was colocalized with CD66b^+^ cells (neutrophil granulocytes) within the incision wound (representative images in Fig. 7A and four additional volunteers (figure S6A)). In addition, MMP9^+^ cells were present in the epidermis (likely keratinocytes) and the dermis (likely being endothelial cells and mast cells) on ipsi- and contralateral sides (figure S6B). ELANE was colocalized in neutrophil granulocytes within the incision, absent in the epidermal layer (representative images in Figure 7B and four additional volunteers, figure S6B), and localized in neutrophil extracellular traps (NETs) (54,55).

**Fig. 7.**
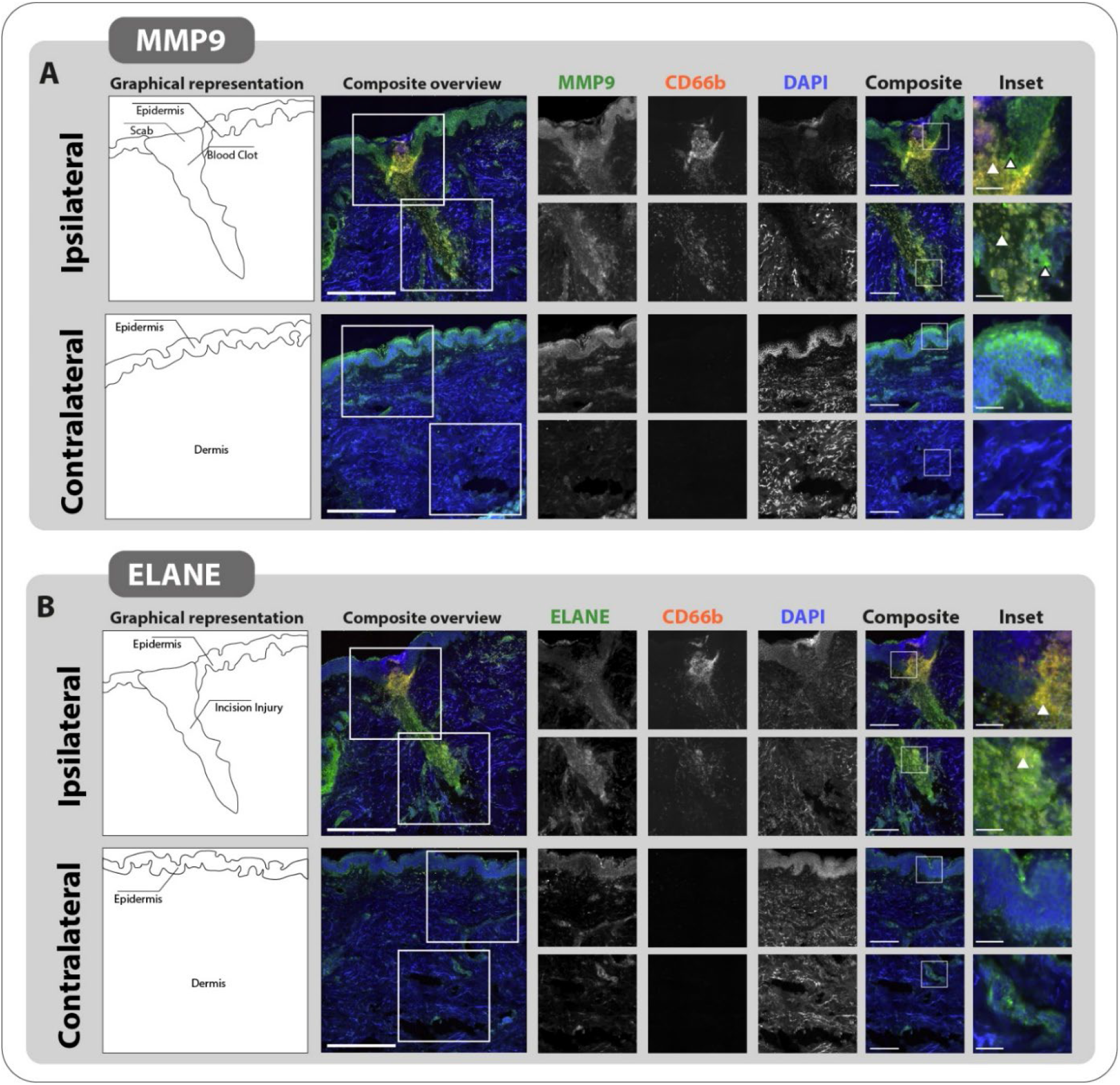
Localization of MMP9 and ELANE in human incised skin. (A) Upon incision, MMP9 colocalized with CD66b^+^ cells (neutrophils, see Inset, white arrow) within the incision injury and in the epidermis layer (see Inset, black/white arrow) on the ipsilateral side. Additionally, in the dermis layer, larger cells are MMP9^+^ (see Inset, black/white arrow). Marginal MMP9 and CD66b signals are detected on the contralateral side with the exception of the epidermis. (B) Upon incision, ELANE colocalized with CD66b^+^ cells (neutrophils, see Inset, white arrow) within the incision injury (in epidermis and dermis layer) on the ipsilateral side. Additionally, in the dermis layer, other cell types are ELANE^+^. Marginal ELANE and CD66b signals are detected on the contralateral side with the exception of the stratum-corneum epidermis. Scale bar = 500 µm(Composite overview), 200 µm (Composite), 50 µm (Inset). MMP9, matrix metalloproteinase 9; ELANE, neutrophil elastase; DAPI, 4′,6-diamidino-2-phenylindole.

## Discussion

Here, we used a translational approach by using parallel unbiased quantitative proteomics in human and mouse skin after an experimental surgical incision to reveal incision-induced protein networks and interspecies specificities as well as variations. This parallel analysis of protein networks reveals both species-specific differences and conserved overlap in the clusters of *platelet aggregation*, *inflammatory activity*, and *actin polymerization* after incision. Furthermore, we identified incision-induced responder types in humans based on phenotypic sensory and pain profiling. In particular, we have focused on the hyperalgesic area (HA) to mechanical stimulation - a typical and clinically relevant pain-related phenotype after skin incision, which is predictive of chronic pain after surgery in patients (24,26,27). Corresponding responder-type protein signatures in humans indicate first and unique insights into cutaneous PPI within 24 h post-incision and represent a protein snapshot, which correlates with incision-induced pain-related symptoms.

Many candidate molecules have been identified, which are presumably relevant for incision-induced nociceptor activation and sensitization causing pain and hyperalgesia, predominantely by using rodent incision models (15–17). Yet complex interplay between the incision-induced proteome for hyperalgesia is unknown, and translation from mice to humans is currently sparse. To address this, we used the well-established experimental incision models in mice and human volunteers and combined this with state-of-the-art proteomic analysis. We identified 435 (mouse) and 119 (human) proteins that were regulated after skin incision, with 50 proteins overlapping in both species. For example, we observed similarities in distinct protein candidates and altered protein networks related to *platelet aggregation*, *inflammatory activity*, and *actin polymerization*. The deciphered similarities in inflammatory activity in both species suggest that the interplay between inflammatory mediators and activation/sensitization of nociceptive terminals is to some extent evolutionarily conserved. This is particularly evident because pain and sensory phenomena such as mechanical hypersensitivity develop in the incision model in humans (39) and rodents (56).

However, our data highlight substantial species differences in PPI networks after incision. For example, *post-translational protein modification*, *metabolism of proteins*, and *metabolism of lipids* are predominant *Reactome* terms in mice after incision, whereas *extracellular matrix organization* is prevalent in human samples. These results indicate species-specific differences, especially in the healing strategy of the incisional wound in accordance with previous reports (57). Thus, relevant networks for mechanical hyperalgesia in humans beyond inflammatory mediators should rather be investigated in human volunteers and not in mice in the future. Only a few studies investigated differences between rodents and human tissue in an unbiased approach in the context of pain (19,58). Recent transcriptomic studies of pain-relevant structures such as peripheral nerves (58) and dorsal root ganglia (19) indicated species-specific similarities. However, these studies compared tissues at the transcriptomic (not proteomic) level and did not decipher signatures of chronic pain. Here, the PPI signatures revealed after incision in humans and mice may – in contrast to existing transcriptome data (19,58)– question a robust translatability (from mice to humans). This may partly explain the low success rate of developing analgesic drugs based on preclinical experiments mainly performed in rodents. In this context, it may be advantageous to research pigs (28), in which physiology, skin composition, and wound healing mechanisms are more similar to humans (16). In general, preclinical animal research lacks the possibility of multidimensional consideration, especially phenotyping and phenotype–related stratification. Overall, studies in humans are mandatory when aiming at deciphering pathological mechanisms underlying pathological pain.

Besides identification of species-specific protein networks after incision, our comprehensive translational approach integrates for the first time pain-related phenotyping with an unbiased proteomic approach in humans. The sensory profiling data were harnessed to stratify human volunteers into High and Low responders, thus, defining the functional relevance of a suite of proteins and their networks strongly associated with pain-related sensory profiles. Hyperalgesia after an incision is critical because it relates to the chronification of post-surgical pain (24,26,27). In addition, mechanical hyperalgesia is a general phenomenon inherent to many human pain models as well as pain patients, but clinically relevant biomarkers are lacking here (21). By stratifying for High versus Low responders related to the area of secondary mechanical hyperalgesia (HA), we revealed hitherto unknown protein signatures that correlate with the extent of HA and its time course. Thus, our results not only provide unprecedented mechanistic insights into incision injury (see below) but, importantly, serve as a stepping stone for phenotype-specific subgrouping of pain-related symptoms via protein signatures. We expect that such studies – e.g., profiling etiology- as well as symptom-specific PPI networks in humans and mice, will undoubtedly provide precious insights into biomarkers relevant for distinct pain-associated symptoms with high translational potential in the future.

Based on our protein network analysis in combination with sensory profiling stratification (High versus Low responder), we are able to propose mechanisms on the PPI level that are relevant for hyperalgesia development and maintenance after incision in humans. In particular, we predict a proteolytic environment in High responders associated with a hypercatabolic state resulting in a prolonged immune response (59). The proteolytic environment is represented by a higher abundance of proteases in central network positions, causing an imbalance of extracellular matrix (ECM) deposition and degradation. Colocalization of proteolytic proteins (MMP9, ELANE) with neutrophil granulocytes and other cells types at the incision site was revealed. Increased immigration of immune cells or possibly functional changes in High responders could indicate overactivation and prolongation of the inflammatory response, leading to a prolonged peripheral sensitization. In turn, this results in extended activation of nociceptive terminals that facilitate central sensitization processes causing large hyperalgesic areas and chronification of pain (45). A similar disbalance in proteases has been observed in chronic, unhealed, and painful wounds (46,47). In Low responders, an opposing picture was evident. Here, anti-inflammatory processes predominate, indicating a decrease in inflammation and the transition to the remodeling phase. The persistence of peripheral inflammation in the High responders and the predominance of the anti-inflammatory environment in the Low responders that correlate with the hyperalgesic area’s extent indicate potential networks targeted for future therapeutic intervention. Of note, modulation of central proteins within a PPI that affect essential, physiological aspects of cutaneous wound healing may compromise the integrity of the network and might therefore be primarily unsuitable because they possess dichromatic properties, such as MMP9 (53,60). Topological properties, eccentricity, and modularity should characterize potential drug target proteins and coreness to determine their ability to interfere with network-relevant proteins or transmit their biological stimulus to other related proteins in the network (61). Taken together, our phenotyping passed stratification and PPI analysis likely differentiate those networks more relevant for hyperalgesia and those relevant for other aspects not involved in pain-related sensations.

From a technical point of view, specific aspects and limitations need to be considered for interpreting our results. Skin samples were obtained within the incision site and contained proteins from several cell types: diverse cutaneous cells, nerve endings, and vascular and immune cells. Given this cellular complexity and heterogeneity, we cannot assign detected changes in protein abundance to specific cell types. This limitation is common to all “-omics” approaches not performed on the single-cell level. Thus far, RNA-seq is the method of choice for comprehensive molecular profiling on the single-cell level. Indeed, previous work has resulted in the successful and extensive characterization of skin cells by means of their transcriptome (62–64). However, the transcriptome and proteome correspondence is known to be limited, with consequential challenges to predict functional alterations on the mRNA level (33–35). Recent excellent work by Dyring-Andersen and colleagues has reported detailed molecular profiles of distinct human skin cell types on the protein level in healthy volunteers (65). Our work considerably extends these efforts by providing (i) inter-species proteome profiles, (ii) altered inter-species protein signatures upon incision-injury, and (iii) their correlation with pain-associated phenotypes in human volunteers. In summary, our findings show – for the first time –interspecies similarities as well as disparities of the cutaneous proteome signature upon incision injury, demonstrating the challenges that bidirectional translational approaches will have to overcome in the future. Based on these results, previous and future work aiming at identifying pain-related mechanisms should be critically assessed, e.g., if performed in rodents. Despite various challenges, ethically well-designed studies on human tissues are mandatory for successful analgesic drug development. In addition, parallel comparative studies in humans and rodents are required to assess the utility of preclinical rodent experiments by pre-determining species–related similarities before translational studies are considered. Hitherto unknown limitations of back-translation into the mouse, not only from an anatomical but also from a functional point of view, urgently need attention. Based on our findings revealing distinct incision-induced phenotypes, might be possible to identify and modulate multiple candidate proteins on the PPI network-level that could be harnessed to ameliorate post-surgical consequences (pain and wound healing) and, consequently, prevent pain chronification.

## Methods

### Human volunteers

In total, 26 male volunteers (age Mean 24± SEM 3.64 years; 2 left-handed) were recruited by notice boards around the university and medical campus (Fig 1). All volunteers enrolled passed our inclusion and exclusion criteria (see table S1). Each volunteer was informed about the study and experimental procedure prior study start and provided written informed consent. They have completed a set of psychological questionnaires, including the Beck’s Depression Inventory (BDI), the state-trait anxiety inventory (STAI), the pain catastrophizing scale (PCS), and the pain sensitivity questionnaire (PSQ), which is a self-rating instrument for the assessment of pain sensitivity, validated in healthy subjects and chronic pain patients (66–70) (table S2). The volunteers were asked to have the same breakfast three days in a row, and the testing was proceeded upright and seated with their forearms on armrests. The test and control sides were randomized beforehand.

### Quantitative sensory testing (QST)

Before incision, a comprehensive battery of quantitative sensory testing (QST) was applied to each volar aspect of the forearm to assess perception on non-painful and painful stimuli of different modalities established by the German Research Network in Neuropathic Pain (DFNS) (71). Briefly, thermal testing was performed by using a TSA-II NeuroSensory Analyzer (MEDOC, Ramat Yishai, Israel). With this device, thermal detection (warm, cold), pain thresholds (painful heat and cold), and pain to a defined suprathreshold painful stimulus will be determined (72). The Advanced Thermal Stimulator thermode (contact area of 16 x 16mm) was placed on the volar forearm. Testing starts at a neutral temperature (32°C) and increases (or decreases) with 1.5°C/s up to a maximum of 50°C (or down to a minimum of 0°C). First, thresholds of cold (CDT) and warm (WDT) detection were measured three times each, followed by an assessment of pain cold (CPT) and heat (HPT) thresholds. Volunteers were instructed to press a button when sensation changes to warm/cold or become painful; then, the thermode immediately returns to baseline temperature (32°C). If the cut-off temperature (50°C or 0°C) was reached, the device automatically returns to baseline temperature (32°C) to avoid tissue damage. The mean value from three measurements was defined as the heat and cold pain threshold. In addition, the pain intensity to a suprathreshold heat pain stimulus was assessed by applying three test stimuli (45, 46, and 47°C, using a ramp of 8°C/s and time at the target of 7 s). In addition, subjects were asked about paradoxical heat sensations (PHS) during the thermal sensory limen (TSL) procedure of alternating warm and cold stimuli.

Modified von Frey hairs (Optihair2-Set; Marstock Nervtest, Schriesheim, Germany) that exert forces between 0.25 and 512 mN (geometric progression by factor 2) was used to assess mechanical detection threshold (MDT). Using von Frey filaments (0.5 mm diameter) with rounded tip ends ensured undesired activation of nociceptors due to sharp edges. Threshold determinations were made using an adaptive method of limits by a series of alternating ascending and descending stimulus intensities (up-and-down rule), yielding 5 just suprathreshold and 5 subthreshold estimates. The geometric mean of these 10 determinations was represented the final threshold.

Standardized punctate probes performed the assessment of mechanical pain sensitivity and threshold with fixed intensities (8, 16, 32, 64, 128, 256, and 512 mN), and a contact area of 0.2 mm diameter, were used to determine mechanical pain sensitivity (MPS) and mechanical pain thresholds (MPT) (39,71). Stimulators were applied at a rate of 2 s on, 2 s off in ascending order until the first percept of sharpness was reached to assess MPT. The final threshold was the geometric mean of five ascending and descending stimuli series. To determine MPS the same stimuli were applied in a pseudorandom order with a 10 sec interval in 5 runs; subjects were asked to give a pain rating for each stimulus on a ‘0–100’ numerical rating scale (’0’ indicating “no pain”, and ‘100’ indicating “most intense pain imaginable”). MPS was calculated as the geometric mean of all numerical ratings for each pinprick stimulus.

QST parameters were measured in their physical dimension and were weighted by transformation to the standard normal distribution (Z-transformation). Z-scores indicate gain (above “0”) or a loss (below “0”) of function across QST-parameters.

### Experimental human skin incision

As previously described, the experimental incision was proceeded in one randomized ipsilateral volar forearm (23,39,73). Briefly, the skin was disinfected (70% ethanol) and incised 4 mm wide and 7 mm deep, perforating the muscular fascia by sterile scalpel (No. 11). Bleeding was stopped by a gauze swab and gentle press.

### Ongoing pain caused by incision

Using the numeric rating scale (NRS; 0-100), the volunteers were asked to rate the intensity of ongoing pain before the incision (baseline), during and after incision for one hour (first every minute for ten minutes, then every 5 minutes) and at 4 hrs and 24 hrs after the incision.

### Determination of mechanical hyperalgesia area and around the incision (73)

For mechanical hyperalgesia around the incision, MPS and MPT were assessed 5 mm apart from the incision (primary hyperalgesia) and 15 mm outside of the incision (secondary hyperalgesia).

Finally, the area, which is sensitive to a usually not painful punctate mechanical stimulation (area of punctate hyperalgesia), was determined (*2*). Briefly, a conventional von Frey filament with 116 mN bending force was applied to eight imaginary lines at a distance of 45° starting far out the assumed hyperalgesia region with centripetally directing to the incision. All 8 points are linked, depicted on a paper, and the area was determined by Image J (https://imagej.nih.gov/ij/). Testing was performed in random order at the incision site and the contralateral arm before as well as 1 h and 24 h after incision.

### Human skin biopsy

Both ipsilateral and contralateral skin samples were taken after an initial toponarcosis with 2% Xylocain and disinfection (Octenisept®, Schülke, Germany) 24 h after incision. Hence, a skin punch of 4 mm in diameter was used to extract the skin biopsies. The skin biopsies contained epidermins and dermis compartments and were stored at −80°C for mass spectrometry and 4°C for immunohistochemical staining.

### Plantar incison in mouse

Adult male C57BL/6J mice [total n = 24 mice, 10weeks, weight 25.4 ± (SD) 1.8 g] were kept in a 12/12 h day/night cycle with ad libitum access to food and water under standardized specific-pathogen-free (SPF) conditions. According to their experimental group, mice were housed together (five per cage) to minimize intergroup effects. Using the Microsoft® Excel randomization function, mice were randomly assigned to the groups. For paw incision in mice (14), all animals were intialley anesthetized with 5% isoflurane (in 100% oxygen) and maintained with 1,5-2% via a nose cone. Plantar aspect of the right hindpaw was disinfected by 100% ethanol and betadine® (povidoneiodine). A 0.5 cm longitudinal incision was performed through the skin and fascia of the plantar aspect of the right hind paw. The incision was started 2 mm from the proximal edge of the heel. The underlying muscle (musclus flexor digitorum brevis) was incised longitudinally and short-time retracted with forceps (Dumont#5). Skin incision was closed with one mattress suture of 6-0 prolene. Betadine® was used to disinfect after the procedure. Sham (control) mice were anesthetized (procedure see above) for the same duration, but no incision and sutures were performed.

### Determination of non-evoked pain behavior at rest

A novel assay to measure the unbalanced distribution of weight-bearing caused by guarding the incised hindpaw was used to assess non-evoked pain-related (38). Briefly, the green-illuminated hindpaw area was determined by ImageJ, and the ratio between contralateral (control) and the incised site was calculated from 10 images over 10 minutes. According to predefined exclusion criteria, images were excluded if grooming, rearing, or hind paw movement was detected. A decreased ratio represents the degree of guarding pain at rest in the incised hindpaw. Both groups did not overlap values or outliers (ROUT method, Q=5%); all mice were included for the skin biopsy.

### Mouse skin biopsy

Mice were euthanized by CO_2,_ and the incised skin (dermis and epidermis, without muscle tissue) was removed by 4 mm skin punch. Skin samples were snap-frozen in liquid nitrogen and stored at −-80°C.

### Tissue lysis and protein solubilization

Frozen skin tissue was manually homogenized, as previously described (74). Skin samples from three mice were pooled to obtain four replicates per experimental condition. Human skin biopsies were treated individually. Sample-related modifications to the published protocol are the following: human skin biopsies were washed in ice-cold PBS to remove exsessive blood and cut in quarters prior to tissue homogenization with a glass potter/pestel. Mouse skin biopsies were directly used for homogenization. Additional shearing of tissue lysates was done with the help of a 20G and a 25G canule and syringe. In order to remove potential DNA contaminations, samples were sonicated for 1 min. Afterwards, samples were solubilized for 15 min, 1100 rpm, at 70°C in a ThermoMixer. Removal of cell debris was achieved by centrifugation at 10000xg for 5 min at 18°C. Acetone precipitation of supernatants was done with five times excess of pre-cooled (- 20°C) acetone and incubation overnight at xym20°C. Precipitated proteins were centrifuged at 14,000xg for 30 min, washed with ice-cold 80% ethanol, and centrifugated at 14,000xg for 30 minutes. After complete removal of ethanol, proteins were air-dried and resuspended in 2% SDS lysis buffer. To ensure complete resuspension, samples were incubated for 15 min at 1100 rpm at 60°C in a ThermoMixer. Flash-frozen samples were stored at −80°C until further use.

### Singlepot, solid-phase-enhanced sample-preparation (SP3), and C18 cleanup

The SP3 protocol was based on the paramagnetic bead-based approach presented by Hughes et.al (75) with sample-dependent optimizations. For human and mouse skin samples, the protein concentration was adjusted to 25 µg in a total of 100 µL with freshly prepared SDS lysis buffer (2% SDS in 100mM Tris, 10mM DTT, 5% Glycerol, pH 7.5). The additional reducing agent was added to aim for the final molarity of 10 mM Dithiothreitol (DTT), counteracting for dilutions caused by sample and buffer mixing. DNA and RNA shearing was achieved by sonication (5 cycles, 30 s on and 60 s- off) to prevent negative influences in bead handling later on. Proteins were reduced at 60°C for 30 min at 1000 rpm in a ThermoMixer. Subsequent alkylation of reduced disulfides was done in the presence of 20 mM Iodoacetamide (IAA) in the dark. The reaction was quenched by the addition of a final of 5 mM DTT for 15 min. Paramagnetic beads (BD) were prepared due to the manufacturer’s instructions and lastly resuspended in SDS lysis buffer with a final concentration of 50 µg/µL. In between steps, the 1.5 mL Eppendorf tube was placed on a magnetic rack for two minutes to retain attached biomolecules by magnetic force while removing the corresponding working solution. Beads and proteins were carefully mixed in a 10:1 ratio (250 µg beads: 25 µg protein) to work in the optimal capacity range at neutral pH conditions. Binding was induced by buoyant mixing with 100% ethanol in a 1:1 ratio (vol/vol) for 5 min at 24°C and 1000 rpm agitation. Three off-magnet rinses with 80% ethanol aimed for removal of detergents and contaminants. After complete removal of washing solutions, 100 µL of 0.1 µg/µL Trypsin/rLys-C (Promega) in 100 mM ammonium bicarbonate (pH 8) was added. Adequate sonication steps of 15 s were performed to promote the disaggregation of beads. Proteolysis was performed overnight at 37°C in a ThermoMixer at 1000 rpm with a heatable in-house built lid to prevent condensation. After 16 h, centrifugation at 1000xg at 24°C for 1 min assisted in bead settling to avoid any carryover to subsequent processing steps. Elution of proteolytic peptides was accomplished by placing the Eppendorf tube on a magnetic rack and transferring the supernatant to a fresh tube. Peptides were acidified to a final concentration of 0.1% trifluoro-acetic acid (TFA) for enhanced reversed-phase purification with commercially available C18 columns (UltraMicroSpin, The Nest Group). While non-polar solutes such as peptides were retained, impurities were removed by centrifugations (110xg, 1 min) between each step. Samples were loaded to conditioned [100 µL 80% acetonitrile (ACN), 0.1% TFA] and equilibrated (100 µL 0.1% TFA) spin columns. After two washing steps with 50 µL 0.1% TFA, peptides were collected by pooling two elutions of 50 µL 50% ACN, 0.1% TFA into fresh Eppendorf tubes. Salt-free peptides were completely dried at 45°C in a centrifugal evaporator. For LC-MS analysis, samples were dissolved in 20uL buffer A (1% ACN, 0.1% FA) and afterward sonicated for 2 min. After centrifugation at 1000xg for 15min, the supernatant was transferred to a fresh Eppendorf tube, and peptide concentration was determined by a nanodrop spectrometer. Human skin samples were adjusted to a final concentration of 25 ng/µL and mouse skin samples to 50 ng/ µL in buffer A. For accurate prediction of retention times, indexed Retention Time (iRT) peptides (Biognosys)(76) in a 1:15 ratio were spiked in.

Exclusion of samples based on quality criteria: Due to low quality and high lipid content, samples from human volunteers, V8, V10, and V19 had to be excluded and were not measured by DIA-MS, marked in red). Poor mass spectrometry chromatograms caused the exclusion of two human biological replicates and two pooled mice samples (1xB_inc_M, 1xB_sham_M).

### Quantitative liquid chromatography-mass spectrometry (LC-MS)

Samples were analyzed on a Q-Exactive HF-X Orbitrap mass spectrometer coupled to an UltiMate 3000 UHPLC System (both Thermo Fisher Scientific). Peptides were separated on a 50cm uPAC column (PharmaFluidics) and directly injected (2 µL) to the mass spectrometer via a nano-electrospray ion source (Thermo Fisher). Flow rates for loading and equilibration were set to 600nl/min while peptide separation along the gradient was done at a constant flow rate of 300nl/min. Peptides of both sample types were loaded in buffer A and eluted in buffer B (85% ACN, 0.1% FA) with a gradient from 2% - 35% for 90 minutes and 35% - 56% for 20 minutes. The step gradient was followed by a 10 min wash of 95% buffer B. While MS acquisition was stopped after the main gradient, 2 short cycles of 10-minute linear gradients (2% - 95% buffer B) were programmed within the same method to reduce carryover between subsequent runs. Column temperature was constantly kept at 50°C. MS raw data were acquired in data-independent acquisition mode (DIA). A MS1 scan in the range of 350 to 1118 m/z (automatic gain control target value of 3e6 or 60ms injection time) at a resolution of 120,000 was followed by a MS2 scan at 30,000 resolution (automatic gain control target value of 3e6 and auto for injection time). In each DIA segment, 14 spectra of 18 Da windows with a stepped collision energy of 25, 27, and 30 were acquired. Thus, one MS cycle consisted of 3 full scans each followed by symmetrically segmented isolation windows. The spectra were recorded in profile mode. The default charge state was set to two.

### DIA-MS Data Analysis

The DIA raw files were analyzed by Spectronaut software version 13.15.200430.43655 (Biognosys(77), Switzerland), and spectra were searched against Uniprot FASTA databases ((HUMAN_Nov2018_UP000005640 or MOUSE_Nov2018_uniprot-proteome_UP000000589, respectively), a common contaminants database, and iRT peptides (78) by the Pulsar^TM^ search engine(79) being integrated into the directDIA^TM^ workflow or a library-based search. Cysteine carbamidomethylation was defined as fixed modification and acetylation of protein N-term and methionine oxidation as variable modification. Enzyme specificity was set to TrypsinP and LysC as protease, allowing two miscleavages within peptide lengths of 7 to 52 amino acids. The false discovery rate (FDR) for precursor, protein, and peptide identification was set to 0.01 per run by searching a decoy database (decoy method-mutated). Label-free quantification was based on the area under the curve between extracted ion currents (XIC) boundaries at MS2 level. TopN strategies for precursors per peptides (min: 1, max: 8) and peptides per protein (min: 2, max: 8) were applied for quantity calculations. The smallest quantitative unit was set to precursor ion. The topN ranking order was determined by a cross-run quality measure (global normalization) of the mean entities within a condition. All data analysis was performed in q-complete mode, i.e. peptides (more specifically precursors, see quantification settings above) were only considered if they were detected across all analyzed samples within one group (for complete DIA-MS datasets, please see table S6 and 7). To assess differential protein abundance between samples, mean log_2_ ratios were calculated in Spectronaut for each protein ID, and pairwise t-tests were performed. Obtained p-values were adjusted using the Benjamini-Hochberg (BH) procedure for multiple testing. Significantly regulated proteins were defined as having a BH-adjusted p-value of q < 0.05. In this way, the following dataset groups were compared: (i) human B_ipsi_ versus B_con_ across 23 samples (Fig. 2 and table S4; human B_ipsi_ versus B_con_ within responder types three somatosensory perceptions (Fig. 3 and table S1), and (iii) mouse B_incM_ versus B_shamM_ across all replicates (Fig. 5 and table S5).

All raw data have been deposited to the ProteomeXchange Consortium via the PRIDE partner repository (80) with the dataset identifier PXD030828.

### DIA-MS dataset post-analysis and comparisons

For all proteome comparisons, we used the following cut-off: proteins that i) exhibited q < 0.05 and ii) were more than 30 % up- or downregulated when comparing B_ipsi_ with B_con_ or, in the case of mouse samples, B_inc_M with B_sham_M. Proteome comparisons were made with respective gene names using http://molbiotools.com/listcompare.html and Excel (Microsoft Office, 2016). If two or more gene names are reported for a single protein, we used the first gene name, which was consistent across all datasets.

The following comparisons were performed:

1. Human B_ipsi_ versus B_con_ across all volunteers compared to mouse B_inc_M versus B_sham_M (Fig. 4).
2. Human B_ipsi_ versus B_con_ for each responder type within the hyperalgesic area, 24 post-incision, and time course (Fig. 5 and figure S4). Proteins that were exclusively regulated in only one responder type (i.e. only in High or only in Low) were defined as being log_2_ FC > |0.38| up- or downregulated in one responder type and not being regulated at all (q > 0.05) in the other responder type.

### Immunohistochemistry staining in human skin

The localization of proteases MMP and ELANE in incised skin was performed with immunohistochemistry (IHC) in a separate cohort of four human volunteers. Skin tissue sections (see “human skin biopsy”) for tissue sampling.

Frozen sections were allowed to dry and adjusted to room temperature. Permeabilization and section blocking was done in 5% donkey serum, 0,4% Triton X-100/PBS for 1h at room temperature. Primary antibodies were diluted in antibody master mix (1% donkey serum, 0,1% Triton X100/PBS) and incubated overnight at 4°C. After a serial of PBS washes (2-3 times with 1mL, three times in washing chamber for 5min with agitation), a fluorophore-conjugated secondary antibody in 1% donkey serum, 0,1% Triton X100/PBS was applied for 2h at room temperature. Unbound antibodies were rinsed with PBS using the above mentioned washing protocol. Sections were mounted in 4’,6-Diamidino-2-Phenylindole (DAPI)-containing SlowFade^TM^ Gold Antifade Medium. All incubation steps, except drying, were conducted in a dark, humified chamber. For antibody specifications, see table S8.

### IHC acquisition and analysis

All images were acquired on a Nikon Ti2 Eclipse inverted widefield fluorescence microscope equipped with a Photometrics Prime 95B 25mm camera system. NIS Image Software (version: 5.21.03) was used to obtain high-resolution data from stitched tiles (1.5mm x 1.5mm). Tissue sections were recorded with a 40×0.6 S Plan FLUOR air objective and excitation wavelength at 395nm, 470nm, and 555nm. Acquisition parameters were kept identical across all experimental groups, including negative controls (omitting primary antibodies).

For raw data processing, the open-source software FIJI (81) was utilized. Z-stacks (N=7, 1.0 µM) were combined to produce a maximum z-projection. Intensity settings of individual channels were adjusted to meet the average min/max value across all biological replicates within each technical procedure.

### Statistics

Sample sizes were a priori determined and are in line with standards in the field. All replicates were biological. Corresponding statistical tests are indicated in the respective figure legend and are two-sided. For proteome data and protein network analysis, we used the Benjamini-Hochberg (BH)-adjusted P-value, Q-value < 0.05. There were no missing data. Data were analyzed using GraphPad Prism 8.0.2 (San Diego, USA). All data are represented as mean ± SEM (standard error) unless indicated otherwise.

### Study approval

All human experiments were approved by the University Hospital Muenster of the local Ethics Committee of the Medical Faculty (registration no 2018-081-b-S), registered in German Clincial Trials Registry (DRKS-ID: DRKS00016641), and in accordance with the latest version of the Declaration of Helsinki.

The animal experiments were reviewed and approved by the Animal Ethics Committee of the State Agency for Nature, Environment, and Consumer Protection North Rhine-Westphalia (LANUV, Recklinghausen, Germany), with the recommendations of the ARRIVE guidelines 2.0 (82)and were in accordance with the ethical guidelines for the investigation of experimental pain in conscious animals (83).

## Author contributions

DS, MvdB, CK, DS-G, JS, SB, and BP performed most experiments and analyzed data unless indicated otherwise. DS, BP and EP-Z designed experiments on mouse behavior. MS and EP-Z conceived and directed the study, supervised experiments and data analysis. DS, MS and EP-Z wrote the manuscript and prepared figures with help from DG-V and PZ.

## Acknowledgments

This work was supported by the Emmy Noether-Program of the Deutsche Forschungsgemeinschaft (DFG), (SCHM 2533/2-1 to MS), a DFG research grant (PO1319/3-1 to EP-Z).

