## Supplemental Tables and Figures for "Species-specific cutaneous protein signatures upon incision injury and correlation with distinct pain-related phenotypes in humans"

**Supplement Table 1. In- and exclusion criteria**

| Criteria | Inclusion | Exclusion |
|---|---|---|
| **Gender** | Male | Female |
| **Age** | 18-40 y | > 40 y |
| **Health Status** | Healthy | Pre-existing diseases<br>• Diabetes<br>• cardiac diseases<br>• pruritus<br>• skin diseases (arm)<br>• neurological diseases |
| **Treatment** | No chronic use of medication | Regular analgesic treatment<br>(< 3days before experiment) |
| **QST** | Unobtrusive | Obtrusive one day before incision |

**Supplement Table 2. Questionnaires**

| Questionnaire | | Score Range | Median [95%CI] | Results |
|---|---|---|---|---|
| Beck-Depressions-Inventar (BDI-II)<br>0-9 clinically unremarkable<br>10-18 mild<br>19-29 moderate<br>>29 severe | | 0-45<br>(a lower score denoting a better outcome) | 4<br>[4 to 7.3] | Clinically unremarkable |
| Revised Life Orientation Test (LOT-R) | Optimism | 0-12<br>(a higher score denoting a better outcome) | 10<br>[8.2 to 10.1] | Clinically unremarkable |
| | Pessimism | 0-12<br>(a higher score denoting a better outcome) | 9<br>[7.8 to 9.5] | Clinically unremarkable |
| | Total | 0-24<br>(a higher score denoting a better outcome) | 19<br>[16.2 to 19.3] | Clinically unremarkable |
| Pain Catastrophizing Scale (PSC)<br>>30 severe | | 0-52<br>(a lower score denoting a better outcome) | 12<br>[9.1 to 14.9] | Clinically unremarkable |
| Pain Sensitivity Questionnaire (PSQ) | Minior | 0-10<br>(a lower score denoting a better outcome) | 2.2<br>[1.8 to 2.4] | Clinically unremarkable |
| | Moderate | 0-10<br>(a lower score denoting a better outcome) | 3.9<br>[3.3 to 4.4] | Clinically unremarkable |
| | Total | 0-10<br>(a lower score denoting a better outcome) | 3.1<br>[2.6 to 3.4] | Clinically unremarkable |
| State-Trait Anxiety Inventory (STAI) | X1 | 20-80<br>(a lower score denoting a better outcome) | 34<br>[31.3 to 36.1] | Clinically unremarkable |
| | X2 | 20-80<br>(a lower score denoting a better outcome) | 35<br>[33.4 to 39.6] | Clinically unremarkable |

### Supplement Table 3. Baseline QST raw values of volunteers

| QST | Control Area | Test Area |
|---|---|---|
| CDT, °C | -1.32 ± 0.53 | -1.53 ± 0.92 |
| WDT, °C | 2 ± 0.86 | 2.2 ± 0.69 |
| TSL, °C | 0.6 ± 0.9 | 0.8 ± 0.8 |
| CPT, °C | -13.38 ± 7.6 | -11.83 ± 8.49 |
| HPT, °C | 9.88 ± 3.9 | 10.20 ± 3.43 |
| MDT, mN | 1.33 ± 1.53 | 1.23 ± 1.36 |
| MPT, mN | 55.82 ± 32.11 | 74.4 ± 142.88 |
| WUR | 3.13 ± 3.1 | 3.1 ± 3.11 |
| VDT, /8 | 7.02 ± 0.38 | 7.11 ± 0.5 |
| PPT, kg | 4.45 ± 1.16 | 4.67 ± 1.16 |

Data are expressed as mean ± SD. Bold numbers indicates *P < 0.05, **P < 0.01, or ***P < 0.001 versus Control Area. Comparisons between areas (control area vs. test area) were performed using paired t-test. Abbreviations: CDT, cold detection threshold; CPT, cold pain threshold; HPT, heat pain threshold; MDT, mechanical detection threshold; MPT, mechanical pain threshold; PPT, pressure pain threshold; QST, quantitative sensory testing; TSL, thermal sensory limen; VDT, vibration detection threshold; WDT, warmth detection threshold; WUR, wind-up ratio.

**Supplement Table 4. Categorization of human volunteers**

(incl. = inclusion, excl. = exclusion)

| Pain Model | | | | | | Proteomics | | | | | | |
|---|---|---|---|---|---|---|---|---|---|---|---|---|
| No | In/Exclusion Criteria | HATC | | HA24 | | No | unbiased | HATC | | HA24 | | Notes |
| | | High | Low | High | Low | | | High | Low | High | Low | |
| 1 | incl. | | | | | 1 | incl. | | | | | |
| 2 | incl. | | | | | 2 | incl. | | | | | |
| 3 | incl. | incl. | | incl. | | 3 | incl. | incl. | | incl. | | |
| 4 | incl. | | incl. | | incl. | 4 | incl. | | incl. | | incl. | |
| 5 | incl. | incl. | | incl. | | 5 | incl. | incl. | | incl. | | |
| 6 | incl. | | | | | 6 | incl. | | | | | |
| 7 | incl. | | incl. | | incl. | 7 | incl. | | incl. | | incl. | |
| 8 | incl. | | incl. | | | 8 | excl. | | excl. | | | high lipid |
| 9 | incl. | incl. | | incl. | | 9 | incl. | incl. | | incl. | | |
| 10 | incl. | | incl. | | | 10 | excl. | | excl. | | | high lipid |
| 11 | incl. | | | | | 11 | incl. | | | | | |
| 12 | incl. | incl. | | incl. | | 12 | incl. | incl. | | incl. | | |
| 13 | incl. | | incl. | | | 13 | incl. | | incl. | | | |
| 14 | incl. | | | | incl. | 14 | incl. | | | | incl. | |
| 15 | incl. | incl. | | incl. | | 15 | incl. | incl. | | incl. | | |
| 16 | incl. | | incl. | | incl. | 16 | incl. | | incl. | | incl. | |
| 17 | incl. | | incl. | | incl. | 17 | incl. | | incl. | | incl. | |
| 18 | incl. | | | | | 18 | incl. | | | | | |
| 19 | incl. | | incl. | | incl. | 19 | excl. | | excl. | | excl. | no skin probe |
| 20 | incl. | incl. | | incl. | | 20 | incl. | incl. | | incl. | | |
| 21 | incl. | | | incl. | | 21 | excl. | | | excl. | | poor XIC |
| 22 | incl. | | incl. | | incl. | 22 | excl. | | excl. | | excl. | poor XIC |
| 23 | incl. | | incl. | | incl. | 23 | incl. | | incl. | | incl. | |
| 24 | incl. | | incl. | | incl. | 24 | incl. | | incl. | | incl. | |
| 25 | incl. | | incl. | | incl. | 25 | incl. | | incl. | | incl. | |
| 26 | incl. | incl. | | | | 26 | incl. | incl. | | | | |
| total | 26 | 7 | 12 | 7 | 10 | total | 21 | 7 | 8 | 6 | 8 | |

**Supplement Table 5. Categorization of mouse smaples**

(incl. = inclusion, excl. = exclusion)

**Pain Model**

| Incision Mice | |
|---|---|
| ID | Non-evoked pain-related behavior |
| Inc_1 | incl. |
| Inc_2 | incl. |
| Inc_3 | incl. |
| Inc_4 | incl. |
| Inc_5 | incl. |
| Inc_6 | incl. |
| Inc_7 | incl. |
| Inc_8 | incl. |
| Inc_9 | incl. |
| Inc_10 | incl. |
| Inc_11 | incl. |
| Inc_12 | incl. |
| total | 12 |
| **Sham Mice** | |
| ID | Non-evoked pain-related behavior |
| Sham_1 | incl. |
| Sham_2 | incl. |
| Sham_3 | incl. |
| Sham_4 | incl. |
| Sham_5 | incl. |
| Sham_6 | incl. |
| Sham_7 | incl. |
| Sham_8 | incl. |
| Sham_9 | incl. |
| Sham_10 | incl. |
| Sham_11 | incl. |
| Sham_12 | incl. |
| total | 12 |

**Proteomics**

| Incision Mice | | |
|---|---|---|
| ID | unbiased | Notes |
| Inc_pool_1 | excl. | poor XIC |
| Inc_pool_2 | incl. | |
| Inc_pool_3 | incl. | |
| Inc_pool_4 | incl. | |
| total | 3 | |
| **Sham Mice** | | |
| ID | unbiased | Notes |
| Sham_pool_1 | incl. | |
| Sham_pool_2 | incl. | |
| Sham_pool_3 | incl. | |
| Sham_pool_4 | excl. | poor XIC |
| total | 3 | |

### Supplement Table 6. Dataset of human DIA-MS

**(bold font represents log2FC > |0.38|)**

| UNIPROT ID | Gene Name | AVG Log2 (INC/con) | N (number of unique total peptides) | P-value | BH (Q-value) | Name |
|---|---|---|---|---|---|---|
| **Q9NTX5** | **ECHDC1** | **-0,729028192** | **2** | **0,03353199** | **0,019710561** | **Ethylmalonyl-CoA decarboxylase** |
| **P02461** | **COL3A1** | **-0,71540446** | **10** | **0,000547382** | **0,000509682** | **Collagen alpha-1(III) chain** |
| **O75635** | **SERPINB7** | **-0,692877544** | **2** | **1,44E-07** | **2,40E-07** | **Serpin B7** |
| **B7Z5J4** | **CPA4** | **-0,660213547** | **6** | **3,21E-10** | **7,41E-10** | **Carboxypeptidase A4** |
| **Q53RT3** | **ASPRV1** | **-0,562719166** | **6** | **2,86E-12** | **8,13E-12** | **Retroviral-like aspartic protease 1** |
| **P08493** | **MGP** | **-0,553585308** | **2** | **7,15E-08** | **1,23E-07** | **Matrix Gla protein** |
| **K7ERG9** | **CFD** | **-0,547964332** | **8** | **4,50E-39** | **5,99E-38** | **Complement factor D** |
| **Q92817** | **EVPL** | **-0,535639766** | **47** | **4,26E-07** | **6,70E-07** | **Envoplakin** |
| **Q96P63-2** | **SERPINB12** | **-0,51987398** | **15** | **1,97E-12** | **5,78E-12** | **Isoform 2 of Serpin B12** |
| **P02452** | **COL1A1** | **-0,512368553** | **37** | **2,35E-12** | **6,76E-12** | **Collagen alpha-1(I) chain** |
| **O60911** | **CTSV** | **-0,512104069** | **5** | **5,88E-12** | **1,62E-11** | **Cathepsin L2** |
| **P00738** | **HP** | **-0,506210684** | **24** | **8,52E-55** | **1,68E-53** | **Haptoglobin** |
| **M0QXX2** | **KLK5** | **-0,501386451** | **2** | **2,88E-05** | **3,27E-05** | **Kallikrein-5 (Fragment)** |
| **P48163** | **ME1** | **-0,488666518** | **7** | **4,82E-11** | **1,22E-10** | **NADP-dependent malic enzyme** |
| **A0A0A0MSS8** | **AKR1C3** | **-0,48361284** | **3** | **1,10E-09** | **2,38E-09** | **Aldo-keto reductase family 1 member C3** |
| **Q96CG8** | **CTHRC1** | **-0,476621298** | **2** | **1,12E-05** | **1,39E-05** | **Collagen triple helix repeat-containing protein 1** |
| **P05543** | **SERPINA7** | **-0,47082763** | **3** | **5,61E-07** | **8,54E-07** | **Thyroxine-binding globulin** |
| **P43155** | **CRAT** | **-0,465249211** | **7** | **0,021344591** | **0,013293087** | **Carnitine O-acetyltransferase** |
| **P31944** | **CASP14** | **-0,463083097** | **17** | **1,15E-24** | **7,34E-24** | **Caspase-14** |
| **Q8IW75** | **SERPINA12** | **-0,460228073** | **7** | **1,14E-12** | **3,40E-12** | **Serpin A12** |
| **E9PHK0** | **CLEC3B** | **-0,45659201** | **5** | **2,03E-17** | **8,59E-17** | **Tetranectin** |
| **Q15517** | **CDSN** | **-0,453016739** | **7** | **4,55E-05** | **5,06E-05** | **Corneodesmosin** |
| **O75342** | **ALOX12B** | **-0,451992993** | **7** | **4,63E-09** | **9,45E-09** | **Arachidonate 12-lipoxygenase, 12R-type** |
| **P36955** | **SERPINF1** | **-0,442648047** | **11** | **3,29E-12** | **9,24E-12** | **Pigment epithelium-derived factor** |
| **Q9NZT1** | **CALML5** | **-0,440051414** | **11** | **6,22E-13** | **1,92E-12** | **Calmodulin-like protein 5** |
| **Q15113** | **PCOLCE** | **-0,437100825** | **12** | **3,36E-22** | **1,94E-21** | **Procollagen C-endopeptidase enhancer 1** |
| **P25325** | **MPST** | **-0,435240634** | **5** | **2,82E-07** | **4,56E-07** | **3-mercaptopyruvate sulfurtransferase** |
| **Q6BCY4** | **CYB5R2** | **-0,430113581** | **3** | **0,029263472** | **0,017494081** | **NADH-cytochrome b5 reductase 2** |
| **Q53EL6** | **PDCD4** | **-0,430006656** | **11** | **1,86E-20** | **9,52E-20** | **Programmed cell death protein 4** |
| **Q96QA5** | **GSDMA** | **-0,425573941** | **13** | **3,02E-12** | **8,53E-12** | **Gasdermin-A** |
| **E9PHN6** | **GSTM2** | **-0,420273946** | **3** | **9,58E-05** | **0,000101569** | **Glutathione S-transferase** |
| **P52895** | **AKR1C2** | **-0,416069293** | **4** | **1,89E-06** | **2,73E-06** | **Aldo-keto reductase family 1 member C2** |
| **Q6ZVX7** | **NCCRP1** | **-0,41506648** | **6** | **2,60E-16** | **1,05E-15** | **F-box only protein 50** |
| **Q9BY32** | **ITPA** | **-0,413357333** | **3** | **0,000869818** | **0,000772473** | **Inosine triphosphate pyrophosphatase** |

| | | | | | | |
|---|---|---|---|---|---|---|
| P09488 | GSTM1 | -0,410947997 | 8 | 1,95E-08 | 3,62E-08 | Glutathione S-transferase Mu 1 |
| Q15126 | PMVK | -0,408623968 | 3 | 0,002785554 | 0,002171938 | Phosphomevalonate kinase |
| P25311 | AZGP1 | -0,399892781 | 12 | 4,52E-41 | 6,54E-40 | Zinc-alpha-2-glycoprotein |
| Q6UWP8 | SBSN | -0,399790922 | 30 | 1,33E-36 | 1,56E-35 | Suprabasin |
| P02790 | HPX | -0,399384315 | 21 | 2,26E-31 | 2,14E-30 | Hemopexin |
| P49862 | KLK7 | -0,393645256 | 5 | 4,28E-06 | 5,84E-06 | Kallikrein-7 |
| Q14914 | PTGR1 | -0,390654609 | 6 | 3,55E-11 | 9,25E-11 | Prostaglandin reductase 1 |
| P08253 | MMP2 | -0,386864374 | 4 | 9,20E-08 | 1,57E-07 | 72 kDa type IV collagenase |
| P22735 | TGM1 | -0,386459212 | 12 | 9,16E-12 | 2,50E-11 | Protein-glutamine gamma-glutamyltransferase K |
| P04062 | GBA | -0,385649025 | 5 | 3,20E-06 | 4,47E-06 | Lysosomal acid glucosylceramidase |
| Q08188 | TGM3 | -0,383331958 | 18 | 2,95E-34 | 3,09E-33 | Protein-glutamine gamma-glutamyltransferase E |
| P14550 | AKR1A1 | -0,380692627 | 7 | 5,77E-17 | 2,41E-16 | Alcohol dehydrogenase [NADP(+)] |
| Q9NTX5 | ECHDC1 | -0,729028192 | 2 | 0,03353199 | 0,019710561 | Ethylmalonyl-CoA decarboxylase |
| P02461 | COL3A1 | -0,71540446 | 10 | 0,000547382 | 0,000509682 | Collagen alpha-1(III) chain |
| O75635 | SERPINB7 | -0,692877544 | 2 | 1,44E-07 | 2,40E-07 | Serpin B7 |
| B7Z5J4 | CPA4 | -0,660213547 | 6 | 3,21E-10 | 7,41E-10 | Carboxypeptidase A4 |
| Q53RT3 | ASPRV1 | -0,562719166 | 6 | 2,86E-12 | 8,13E-12 | Retroviral-like aspartic protease 1 |
| P08493 | MGP | -0,553585308 | 2 | 7,15E-08 | 1,23E-07 | Matrix Gla protein |
| K7ERG9 | CFD | -0,547964332 | 8 | 4,50E-39 | 5,99E-38 | Complement factor D |
| Q92817 | EVPL | -0,535639766 | 47 | 4,26E-07 | 6,70E-07 | Envoplakin |
| Q96P63-2 | SERPINB12 | -0,51987398 | 15 | 1,97E-12 | 5,78E-12 | Isoform 2 of Serpin B12 |
| P02452 | COL1A1 | -0,512368553 | 37 | 2,35E-12 | 6,76E-12 | Collagen alpha-1(I) chain |
| O60911 | CTSV | -0,512104069 | 5 | 5,88E-12 | 1,62E-11 | Cathepsin L2 |
| P00738 | HP | -0,506210684 | 24 | 8,52E-55 | 1,68E-53 | Haptoglobin |
| M0QXX2 | KLK5 | -0,501386451 | 2 | 2,88E-05 | 3,27E-05 | Kallikrein-5 (Fragment) |
| P48163 | ME1 | -0,488666518 | 7 | 4,82E-11 | 1,22E-10 | NADP-dependent malic enzyme |
| A0A0A0MSS8 | AKR1C3 | -0,48361284 | 3 | 1,10E-09 | 2,38E-09 | Aldo-keto reductase family 1 member C3 |
| Q96CG8 | CTHRC1 | -0,476621298 | 2 | 1,12E-05 | 1,39E-05 | Collagen triple helix repeat-containing protein 1 |
| P05543 | SERPINA7 | -0,47082763 | 3 | 5,61E-07 | 8,54E-07 | Thyroxine-binding globulin |
| P43155 | CRAT | -0,465249211 | 7 | 0,021344591 | 0,013293087 | Carnitine O-acetyltransferase |
| P31944 | CASP14 | -0,463083097 | 17 | 1,15E-24 | 7,34E-24 | Caspase-14 |
| Q8IW75 | SERPINA12 | -0,460228073 | 7 | 1,14E-12 | 3,40E-12 | Serpin A12 |
| E9PHK0 | CLEC3B | -0,45659201 | 5 | 2,03E-17 | 8,59E-17 | Tetranectin |
| Q15517 | CDSN | -0,453016739 | 7 | 4,55E-05 | 5,06E-05 | Corneodesmosin |
| O75342 | ALOX12B | -0,451992993 | 7 | 4,63E-09 | 9,45E-09 | Arachidonate 12-lipoxygenase, 12R-type |
| P36955 | SERPINF1 | -0,442648047 | 11 | 3,29E-12 | 9,24E-12 | Pigment epithelium-derived factor |
| Q9NZT1 | CALML5 | -0,440051414 | 11 | 6,22E-13 | 1,92E-12 | Calmodulin-like protein 5 |

| | | | | | | |
|---|---|---|---|---|---|---|
| Q15113 | PCOLCE | -0,437100825 | 12 | 3,36E-22 | 1,94E-21 | Procollagen C-endopeptidase enhancer 1 |
| P25325 | MPST | -0,435240634 | 5 | 2,82E-07 | 4,56E-07 | 3-mercaptopyruvate sulfurtransferase |
| Q6BCY4 | CYB5R2 | -0,430113581 | 3 | 0,029263472 | 0,017494081 | NADH-cytochrome b5 reductase 2 |
| Q53EL6 | PDCD4 | -0,430006656 | 11 | 1,86E-20 | 9,52E-20 | Programmed cell death protein 4 |
| Q96QA5 | GSDMA | -0,425573941 | 13 | 3,02E-12 | 8,53E-12 | Gasdermin-A |
| E9PHN6 | GSTM2 | -0,420273946 | 3 | 9,58E-05 | 0,000101569 | Glutathione S-transferase |
| P52895 | AKR1C2 | -0,416069293 | 4 | 1,89E-06 | 2,73E-06 | Aldo-keto reductase family 1 member C2 |
| Q6ZVX7 | NCCRP1 | -0,41506648 | 6 | 2,60E-16 | 1,05E-15 | F-box only protein 50 |
| Q9BY32 | ITPA | -0,413357333 | 3 | 0,000869818 | 0,000772473 | Inosine triphosphate pyrophosphatase |
| P09488 | GSTM1 | -0,410947997 | 8 | 1,95E-08 | 3,62E-08 | Glutathione S-transferase Mu 1 |
| Q15126 | PMVK | -0,408623968 | 3 | 0,002785554 | 0,002171938 | Phosphomevalonate kinase |
| P25311 | AZGP1 | -0,399892781 | 12 | 4,52E-41 | 6,54E-40 | Zinc-alpha-2-glycoprotein |
| Q6UWP8 | SBSN | -0,399790922 | 30 | 1,33E-36 | 1,56E-35 | Suprabasin |
| P02790 | HPX | -0,399384315 | 21 | 2,26E-31 | 2,14E-30 | Hemopexin |
| P49862 | KLK7 | -0,393645256 | 5 | 4,28E-06 | 5,84E-06 | Kallikrein-7 |
| Q14914 | PTGR1 | -0,390654609 | 6 | 3,55E-11 | 9,25E-11 | Prostaglandin reductase 1 |
| P08253 | MMP2 | -0,386864374 | 4 | 9,20E-08 | 1,57E-07 | 72 kDa type IV collagenase |
| P22735 | TGM1 | -0,386459212 | 12 | 9,16E-12 | 2,50E-11 | Protein-glutamine gamma-glutamyltransferase K |
| P04062 | GBA | -0,385649025 | 5 | 3,20E-06 | 4,47E-06 | Lysosomal acid glucosylceramidase |
| Q08188 | TGM3 | -0,383331958 | 18 | 2,95E-34 | 3,09E-33 | Protein-glutamine gamma-glutamyltransferase E |
| P14550 | AKR1A1 | -0,380692627 | 7 | 5,77E-17 | 2,41E-16 | Alcohol dehydrogenase [NADP(+)] |
| Q7Z5L7 | PODN | -0,379877623 | 9 | 4,86E-07 | 7,59E-07 | Podcan |
| P00325 | ADH1B | -0,378115142 | 17 | 3,02E-24 | 1,91E-23 | Alcohol dehydrogenase 1B |
| Q9BW30 | TPPP3 | -0,375163268 | 3 | 0,00198236 | 0,001601513 | Tubulin polymerization-promoting protein family member 3 |
| Q00796 | SORD | -0,373032422 | 6 | 7,51E-05 | 8,12E-05 | Sorbitol dehydrogenase |
| Q9NR19 | ACSS2 | -0,372030044 | 5 | 0,055967767 | 0,030663854 | Acetyl-coenzyme A synthetase, cytoplasmic |
| P42357 | HAL | -0,370659528 | 12 | 5,28E-08 | 9,28E-08 | Histidine ammonia-lyase |
| Q13867 | BLMH | -0,368365898 | 17 | 1,15E-24 | 7,34E-24 | Bleomycin hydrolase |
| P21266 | GSTM3 | -0,368213778 | 10 | 3,60E-28 | 2,68E-27 | Glutathione S-transferase Mu 3 |
| P60981 | DSTN | -0,368051797 | 7 | 1,75E-06 | 2,53E-06 | Destrin |
| P06727 | APOA4 | -0,363021925 | 18 | 2,32E-29 | 1,97E-28 | Apolipoprotein A-IV |

| | | | | | | |
|---|---|---|---|---|---|---|
| Q9BUT1 | BDH2 | -0,361476142 | 2 | 8,89E-07 | 1,34E-06 | 3-hydroxybutyrate dehydrogenase type 2 |
| Q9UBW8 | COPS7A | -0,361281044 | 3 | 0,097134802 | 0,047802759 | COP9 signalosome complex subunit 7a |
| P28074 | PSMB5 | -0,360822894 | 6 | 2,49E-06 | 3,54E-06 | Proteasome subunit beta type-5 |
| P50440 | GATM | -0,36056588 | 3 | 7,88E-06 | 1,01E-05 | Glycine amidinotransferase, mitochondrial |
| P21399 | ACO1 | -0,359594822 | 15 | 5,08E-21 | 2,69E-20 | Cytoplasmic aconitate hydratase |
| P12277 | CKB | -0,355335987 | 15 | 1,09E-26 | 7,55E-26 | Creatine kinase B-type |
| A6NNA4 | MATN4 | -0,351660275 | 3 | 0,014670314 | 0,009435025 | Matrilin-4 |
| Q6P587 | FAHD1 | -0,350015814 | 3 | 0,00137436 | 0,00116785 | Acylpyruvase FAHD1, mitochondrial |
| O75223 | GGCT | -0,34788491 | 6 | 4,78E-10 | 1,08E-09 | Gamma-glutamylcyclotransferase |
| Q96JY6 | PDLIM2 | -0,346756441 | 3 | 1,51E-05 | 1,82E-05 | PDZ and LIM domain protein 2 |
| Q96D15 | RCN3 | -0,34658425 | 5 | 3,08E-10 | 7,14E-10 | Reticulocalbin-3 |
| A0A3B3IRL2 | CREG1 | -0,345615445 | 2 | 0,068070174 | 0,035666162 | Cellular repressor of E1A-stimulated genes 1, isoform CRA_a |
| P17900 | GM2A | -0,345294187 | 2 | 3,62E-06 | 5,04E-06 | Ganglioside GM2 activator |
| P35914 | HMGCL | -0,345274992 | 2 | 0,000629483 | 0,000579973 | Hydroxymethylglutaryl-CoA lyase, mitochondrial |
| P19652 | ORM2 | -0,344731553 | 7 | 1,00E-06 | 1,50E-06 | Alpha-1-acid glycoprotein 2 |
| A0A087WZF1 | LPP | -0,344440427 | 6 | 1,46E-10 | 3,53E-10 | Lipoma-preferred partner |
| P23142-4 | FBLN1 | -0,343903361 | 4 | 1,12E-07 | 1,89E-07 | Isoform C of Fibulin-1 |
| E7EVA0 | MAP4 | -0,343799263 | 14 | 7,95E-16 | 3,01E-15 | Microtubule-associated protein |
| P02763 | ORM1 | -0,342207222 | 6 | 5,72E-06 | 7,57E-06 | Alpha-1-acid glycoprotein 1 |
| P49327 | FASN | -0,33978491 | 105 | 7,64E-31 | 6,96E-30 | Fatty acid synthase |
| P01009 | SERPINA1 | -0,339405469 | 30 | 4,27E-29 | 3,50E-28 | Alpha-1-antitrypsin |
| Q9H1E1 | RNASE7 | -0,338411005 | 3 | 6,89E-05 | 7,50E-05 | Ribonuclease 7 |
| Q3ZCW2 | LGALSL | -0,337011816 | 2 | 0,026327852 | 0,015972014 | Galectin-related protein |
| P34913 | EPHX2 | -0,336901503 | 6 | 1,93E-05 | 2,26E-05 | Bifunctional epoxide hydrolase 2 |
| Q9H2C0 | GAN | -0,33443696 | 5 | 0,008705746 | 0,005965498 | Gigaxonin |

| | | | | | | |
|---|---|---|---|---|---|---|
| A0A0A0MS41 | SFXN3 | -0,332797992 | 3 | 4,77E-08 | 8,54E-08 | Sideroflexin |
| O75828 | CBR3 | -0,331699704 | 3 | 3,67E-07 | 5,84E-07 | Carbonyl reductase [NADPH] 3 |
| E7ETH0 | CFI | -0,32961309 | 6 | 0,000320712 | 0,00031 | Complement factor I |
| P16152 | CBR1 | -0,328533334 | 12 | 9,87E-42 | 1,52E-40 | Carbonyl reductase [NADPH] 1 |
| P01019 | AGT | -0,326107533 | 12 | 6,07E-11 | 1,52E-10 | Angiotensinogen |
| Q08257 | CRYZ | -0,325840602 | 6 | 4,62E-14 | 1,53E-13 | Quinone oxidoreductase |
| P21810 | BGN | -0,322071248 | 14 | 1,78E-26 | 1,22E-25 | Biglycan |
| P15121 | AKR1B1 | -0,320709443 | 5 | 5,01E-05 | 5,54E-05 | Aldose reductase |
| P04264 | KRT1 | -0,319970752 | 3 | 0,021985635 | 0,01364052 | Keratin, type II cytoskeletal 1 |
| Q14195-2 | DPYSL3 | -0,319061299 | 19 | 5,92E-29 | 4,70E-28 | Isoform LCRMP-4 of Dihydropyrimidinase-related protein 3 |
| P31937 | HIBADH | -0,318541594 | 6 | 1,03E-13 | 3,32E-13 | 3-hydroxyisobutyrate dehydrogenase, mitochondrial |
| Q9UEY8 | ADD3 | -0,317620838 | 5 | 6,62E-06 | 8,59E-06 | Gamma-adducin |
| P23142 | FBLN1 | -0,317367458 | 9 | 1,57E-10 | 3,77E-10 | Fibulin-1 |
| O76041-2 | NEBL | -0,316311673 | 2 | 0,001451584 | 0,001218736 | Isoform 2 of Nebulette |
| P51884 | LUM | -0,311773153 | 10 | 2,16E-19 | 1,06E-18 | Lumican |
| O43294 | TGFB1I1 | -0,310846328 | 4 | 0,024412583 | 0,014976298 | Transforming growth factor beta-1-induced transcript 1 protein |
| Q5T749 | KPRP | -0,3103826 | 4 | 7,67E-05 | 8,27E-05 | Keratinocyte proline-rich protein |
| Q63ZY3 | KANK2 | -0,309262194 | 3 | 1,11E-05 | 1,38E-05 | KN motif and ankyrin repeat domain-containing protein 2 |
| H0YFA4 | CRIP2 | -0,308422211 | 3 | 1,58E-15 | 5,75E-15 | Cysteine-rich protein 2 (Fragment) |
| A0A087WTA8 | COL1A2 | -0,308392222 | 29 | 1,94E-07 | 3,18E-07 | Collagen alpha-2(I) chain |
| B4DNG0 | OLFML3 | -0,304185703 | 13 | 3,64E-18 | 1,64E-17 | cDNA FLJ58142, highly similar to Olfactomedin-like protein 3 |
| P46821 | MAP1B | -0,304001244 | 6 | 1,60E-10 | 3,83E-10 | Microtubule-associated protein 1B |
| Q96C86 | DCPS | -0,301899358 | 3 | 0,006846454 | 0,004825867 | m7GpppX diphosphatase |
| Q15124 | PGM5 | -0,299475214 | 10 | 2,94E-08 | 5,41E-08 | Phosphoglucomutase-like protein 5 |
| P49189 | ALDH9A1 | -0,299137293 | 13 | 1,85E-30 | 1,63E-29 | 4-trimethylaminobutyraldehyde dehydrogenase |
| A0A087WT95 | ACOT2 | -0,29905766 | 5 | 0,002728689 | 0,002137763 | Acyl-coenzyme A thioesterase 2, mitochondrial |

| | | | | | | |
|---|---|---|---|---|---|---|
| D6R956 | UCHL1 | -0,298680467 | 3 | 0,002156199 | 0,001733415 | Ubiquitin carboxyl-terminal hydrolase |
| P36952 | SERPINB5 | -0,297635247 | 24 | 2,47E-27 | 1,76E-26 | Serpin B5 |
| P30086 | PEBP1 | -0,29760322 | 12 | 1,34E-20 | 6,92E-20 | Phosphatidylethanolamine-binding protein 1 |
| O00534 | VWA5A | -0,296608248 | 5 | 1,62E-11 | 4,37E-11 | von Willebrand factor A domain-containing protein 5A |
| O95833 | CLIC3 | -0,296417754 | 4 | 0,009384939 | 0,00637761 | Chloride intracellular channel protein 3 |
| Q08554 | DSC1 | -0,295460596 | 26 | 7,17E-16 | 2,76E-15 | Desmocollin-1 |
| A0A0A0MSE2 | HADH | -0,294341871 | 8 | 0,00192728 | 0,001562145 | Hydroxyacyl-coenzyme A dehydrogenase, mitochondrial |
| A0A1B0GTG2 | ALDH7A1 | -0,293969277 | 15 | 2,35E-11 | 6,24E-11 | Alpha-aminoadipic semialdehyde dehydrogenase |
| P05089 | ARG1 | -0,293292848 | 12 | 1,69E-10 | 3,99E-10 | Arginase-1 |
| O14791 | APOL1 | -0,292463136 | 4 | 0,045485873 | 0,025693513 | Apolipoprotein L1 |
| D6RF35 | GC | -0,292426617 | 21 | 1,27E-15 | 4,65E-15 | Vitamin D-binding protein |
| E9PR44 | CRYAB | -0,291791082 | 5 | 9,20E-09 | 1,80E-08 | Alpha-crystallin B chain (Fragment) |
| A0A0A0MS87 | NDRG2 | -0,289198822 | 5 | 3,10E-07 | 4,97E-07 | Protein NDRG2 |
| F6TLX2 | GLOD4 | -0,289010674 | 6 | 3,53E-12 | 9,81E-12 | Glyoxalase domain-containing protein 4 |
| P02766 | TTR | -0,288805185 | 11 | 3,46E-12 | 9,67E-12 | Transthyretin |
| Q16643 | DBN1 | -0,288151699 | 3 | 0,000153114 | 0,000155967 | Drebrin |
| P08670 | VIM | -0,286198281 | 46 | 1,62E-18 | 7,54E-18 | Vimentin |
| P06312 | IGKV4-1 | -0,285798208 | 2 | 0,037657583 | 0,021771481 | Immunoglobulin kappa variable 4-1 |
| P48637 | GSS | -0,285415099 | 7 | 1,33E-05 | 1,62E-05 | Glutathione synthetase |
| A0A0B4J2A4 | ACAA2 | -0,284503454 | 14 | 1,29E-07 | 2,17E-07 | 3-ketoacyl-CoA thiolase, mitochondrial |
| P49441 | INPP1 | -0,283814228 | 3 | 0,00149954 | 0,001248348 | Inositol polyphosphate 1-phosphatase |
| J3KNQ4 | PARVA | -0,282154643 | 5 | 0,000238389 | 0,000235991 | Alpha-parvin |
| Q9Y617 | PSAT1 | -0,280421564 | 9 | 7,86E-18 | 3,48E-17 | Phosphoserine aminotransferase |
| A0A3B3ISG5 | IDE | -0,279356308 | 17 | 6,49E-19 | 3,04E-18 | Insulin-degrading enzyme |
| Q96L46 | CAPNS2 | -0,278891184 | 4 | 2,53E-06 | 3,57E-06 | Calpain small subunit 2 |

| | | | | | | |
|---|---|---|---|---|---|---|
| P29373 | CRABP2 | -0,278701112 | 6 | 9,38E-16 | 3,50E-15 | Cellular retinoic acid-binding protein 2 |
| Q9UBG0 | MRC2 | -0,278556544 | 4 | 0,009704234 | 0,006556977 | C-type mannose receptor 2 |
| P08294 | SOD3 | -0,277964885 | 8 | 6,03E-08 | 1,05E-07 | Extracellular superoxide dismutase [Cu-Zn] |
| P80365 | HSD11B2 | -0,277831018 | 4 | 0,020155728 | 0,012664902 | Corticosteroid 11-beta-dehydrogenase isozyme 2 |
| Q09666 | AHNAK | -0,275926019 | 227 | 1,00E-257 | 1,64E-255 | Neuroblast differentiation-associated protein AHNAK |
| Q9BQ69 | MACROD1 | -0,274367224 | 3 | 0,000319232 | 0,000309177 | ADP-ribose glycohydrolase MACROD1 |
| P27482 | CALML3 | -0,273198064 | 7 | 2,92E-05 | 3,31E-05 | Calmodulin-like protein 3 |
| P07585 | DCN | -0,269346492 | 21 | 3,33E-17 | 1,40E-16 | Decorin |
| P23634 | ATP2B4 | -0,268048468 | 7 | 0,073586436 | 0,038111185 | Plasma membrane calcium-transporting ATPase 4 |
| P51888 | PRELP | -0,267752178 | 16 | 1,02E-09 | 2,22E-09 | Prolargin |
| I3L297 | CRK | -0,267245379 | 2 | 4,06E-06 | 5,60E-06 | Adapter molecule crk |
| P51178 | PLCD1 | -0,266714832 | 4 | 4,40E-06 | 5,99E-06 | 1-phosphatidylinositol 4,5-bisphosphate phosphodiesterase delta-1 |
| O95865 | DDAH2 | -0,266692286 | 6 | 1,08E-08 | 2,08E-08 | N(G),N(G)-dimethylarginine dimethylaminohydrolase 2 |
| P11766 | ADH5 | -0,266053477 | 9 | 5,45E-14 | 1,78E-13 | Alcohol dehydrogenase class-3 |
| H0YAC1 | KLKB1 | -0,265687632 | 5 | 0,026460777 | 0,016032885 | Plasma kallikrein (Fragment) |
| O60749 | SNX2 | -0,265673711 | 5 | 9,46E-06 | 1,20E-05 | Sorting nexin-2 |
| Q96CN7 | ISOC1 | -0,263758743 | 3 | 0,040985489 | 0,023323666 | Isochorismatase domain-containing protein 1 |
| P05120 | SERPINB2 | -0,263355394 | 5 | 1,35E-06 | 1,98E-06 | Plasminogen activator inhibitor 2 |
| P08185 | SERPINA6 | -0,262368981 | 6 | 0,07497217 | 0,038664893 | Corticosteroid-binding globulin |
| P02787 | TF | -0,260640323 | 47 | 8,03E-20 | 4,03E-19 | Serotransferrin |
| P04217 | A1BG | -0,260443025 | 13 | 6,63E-10 | 1,48E-09 | Alpha-1B-glycoprotein |
| P13489 | RNH1 | -0,260291793 | 16 | 3,13E-15 | 1,12E-14 | Ribonuclease inhibitor |
| Q9BV20 | MRI1 | -0,259654036 | 3 | 0,083625687 | 0,042401229 | Methylthioribose-1-phosphate isomerase |
| P04792 | HSPB1 | -0,256290568 | 16 | 7,84E-30 | 6,77E-29 | Heat shock protein beta-1 |
| Q9BWD1 | ACAT2 | -0,255976773 | 7 | 3,87E-05 | 4,34E-05 | Acetyl-CoA acetyltransferase, cytosolic |

| | | | | | | |
|---|---|---|---|---|---|---|
| Q99497 | PARK7 | -0,253518971 | 10 | 7,67E-10 | 1,71E-09 | Protein/nucleic acid deglycase DJ-1 |
| P30085 | CMPK1 | -0,253499078 | 8 | 3,76E-06 | 5,22E-06 | UMP-CMP kinase |
| F5H365 | SEC23A | -0,25348089 | 4 | 1,46E-05 | 1,77E-05 | Protein transport protein SEC23 |
| P11182 | DBT | -0,253230941 | 3 | 0,001636185 | 0,001346159 | Lipoamide acyltransferase component of branched-chain alpha-keto acid dehydrogenase complex, mitochondrial |
| Q6UWY5 | OLFML1 | -0,252708734 | 13 | 2,68E-16 | 1,07E-15 | Olfactomedin-like protein 1 |
| O60504 | SORBS3 | -0,252251158 | 2 | 9,41E-05 | 0,000100047 | Vinexin |
| A0A087WUI4 | TPSB2 | -0,252152767 | 10 | 2,72E-13 | 8,59E-13 | Tryptase beta-2 |
| B7Z9I1 | ACADM | -0,251016119 | 9 | 0,000175044 | 0,000175759 | Medium-chain-specific acyl-CoA dehydrogenase, mitochondrial |
| P18510 | IL1RN | -0,250988719 | 4 | 9,99E-05 | 0,00010543 | Interleukin-1 receptor antagonist protein |
| Q9UBE0 | SAE1 | -0,250511837 | 3 | 5,29E-07 | 8,13E-07 | SUMO-activating enzyme subunit 1 |
| Q96PD5 | PGLYRP2 | -0,250424104 | 7 | 1,06E-10 | 2,56E-10 | N-acetylmuramoyl-L-alanine amidase |
| P00966 | ASS1 | -0,250317649 | 9 | 1,07E-06 | 1,59E-06 | Argininosuccinate synthase |
| P02765 | AHSG | -0,24971036 | 7 | 4,90E-10 | 1,11E-09 | Alpha-2-HS-glycoprotein |
| H0YHN7 | CPM | -0,249560906 | 2 | 0,028345643 | 0,016986671 | Carboxypeptidase M (Fragment) |
| Q06828 | FMOD | -0,2490656 | 6 | 1,11E-05 | 1,38E-05 | Fibromodulin |
| Q07960 | ARHGAP1 | -0,248799438 | 12 | 5,08E-16 | 1,97E-15 | Rho GTPase-activating protein 1 |
| Q9HCY8 | S100A14 | -0,242749263 | 4 | 0,001274409 | 0,001094257 | Protein S100-A14 |
| P00352 | ALDH1A1 | -0,242590011 | 16 | 7,79E-16 | 2,97E-15 | Retinal dehydrogenase 1 |
| P42126 | ECI1 | -0,242493032 | 4 | 1,07E-06 | 1,59E-06 | Enoyl-CoA delta isomerase 1, mitochondrial |
| P17931 | LGALS3 | -0,242440796 | 5 | 0,001643595 | 0,001349998 | Galectin-3 |
| P02760 | AMBP | -0,241631345 | 12 | 2,52E-12 | 7,20E-12 | Protein AMBP |
| J3KS22 | DCXR | -0,24151388 | 7 | 8,02E-09 | 1,59E-08 | L-xylulose reductase (Fragment) |
| Q9BXN1 | ASPN | -0,241367878 | 15 | 1,19E-07 | 2,00E-07 | Asporin |
| Q96C23 | GALM | -0,241054992 | 3 | 0,000788855 | 0,000705666 | Aldose 1-epimerase |
| A0A2Q2TTZ9 | IGKV1D-33 | -0,239194767 | 3 | 0,002383043 | 0,00189718 | Immunoglobulin kappa variable 1-33 |
| P21291 | CSRP1 | -0,238783207 | 8 | 7,54E-06 | 9,68E-06 | Cysteine and glycine-rich protein 1 |

| | | | | | | |
|---|---|---|---|---|---|---|
| Q9UBQ7 | GRHPR | -0,23839815 | 10 | 1,79E-07 | 2,97E-07 | Glyoxylate reductase/hydroxypyruvate reductase |
| B3KSI3 | BCAT2 | -0,238126111 | 2 | 0,077227636 | 0,03962006 | Branched-chain-amino-acid aminotransferase |
| A0A024QZX5 | SERPINB6 | -0,237410742 | 11 | 1,94E-14 | 6,54E-14 | Serpin B6 |
| B7ZC38 | SH3GLB2 | -0,236424425 | 3 | 0,007243874 | 0,005084145 | Endophilin-B2 |
| A0A0C4DH67 | IGKV1-8 | -0,236164545 | 2 | 0,016376728 | 0,010464091 | Immunoglobulin kappa variable 1-8 |
| P55809 | OXCT1 | -0,235487481 | 4 | 3,59E-10 | 8,22E-10 | Succinyl-CoA:3-ketoacid coenzyme A transferase 1, mitochondrial |
| Q8IUX7 | AEBP1 | -0,235202751 | 13 | 7,55E-13 | 2,31E-12 | Adipocyte enhancer-binding protein 1 |
| A0A286YFJ8 | IGHG4 | -0,234972255 | 9 | 2,06E-06 | 2,96E-06 | Immunoglobulin heavy constant gamma 4 (Fragment) |
| Q6NVY1 | HIBCH | -0,234740337 | 3 | 4,00E-05 | 4,46E-05 | 3-hydroxyisobutyryl-CoA hydrolase, mitochondrial |
| P43652 | AFM | -0,234193433 | 10 | 2,24E-05 | 2,58E-05 | Afamin |
| H0Y8X4 | DNPH1 | -0,234034647 | 4 | 0,056430578 | 0,030848716 | 2'-deoxynucleoside 5'-phosphate N-hydrolase 1 (Fragment) |
| A0A286YF22 | PHGDH | -0,233646411 | 14 | 6,01E-17 | 2,48E-16 | D-3-phosphoglycerate dehydrogenase |
| Q14116 | IL18 | -0,232922843 | 4 | 5,02E-06 | 6,74E-06 | Interleukin-18 |
| A0A1C7CYX9 | DPYSL2 | -0,232351361 | 24 | 1,14E-28 | 8,90E-28 | Dihydropyrimidinase-related protein 2 |
| O75874 | IDH1 | -0,230676967 | 16 | 2,94E-19 | 1,42E-18 | Isocitrate dehydrogenase [NADP] cytoplasmic |
| P01042-2 | KNG1 | -0,230528081 | 12 | 7,35E-11 | 1,84E-10 | Isoform LMW of Kininogen-1 |
| O95302 | FKBP9 | -0,227814613 | 3 | 2,56E-08 | 4,73E-08 | Peptidyl-prolyl cis-trans isomerase FKBP9 |
| P05546 | SERPIND1 | -0,226486834 | 11 | 4,23E-11 | 1,08E-10 | Heparin cofactor 2 |
| P04066 | FUCA1 | -0,225092995 | 2 | 0,004820007 | 0,003528933 | Tissue alpha-L-fucosidase |
| P05155 | SERPING1 | -0,224028371 | 12 | 3,89E-05 | 4,35E-05 | Plasma protease C1 inhibitor |
| E9PHS0 | LANCL1 | -0,224013006 | 4 | 0,000206995 | 0,000205741 | LanC-like protein 1 (Fragment) |
| Q13642-1 | FHL1 | -0,223939481 | 7 | 2,85E-07 | 4,59E-07 | Isoform 1 of Four and a half LIM domains protein 1 |
| A0A087X142 | SEPTIN8 | -0,223898817 | 5 | 4,18E-05 | 4,65E-05 | Septin-8 |
| A0A087X0D5 | CTSH | -0,223841971 | 4 | 0,001069448 | 0,000932922 | Pro-cathepsin H |
| A6NMZ7 | COL6A6 | -0,223807564 | 30 | 1,09E-09 | 2,38E-09 | Collagen alpha-6(VI) chain |

| | | | | | | |
|---|---|---|---|---|---|---|
| Q01995 | TAGLN | -0,221796676 | 14 | 8,04E-05 | 8,59E-05 | Transgelin |
| P36269 | GGT5 | -0,221174475 | 5 | 0,050829894 | 0,028418532 | Glutathione hydrolase 5 proenzyme |
| Q16762 | TST | -0,219866759 | 2 | 0,004354058 | 0,003226199 | Thiosulfate sulfurtransferase |
| P38117 | ETFB | -0,219166739 | 7 | 9,82E-15 | 3,38E-14 | Electron transfer flavoprotein subunit beta |
| Q15075 | EEA1 | -0,218796577 | 6 | 6,26E-06 | 8,20E-06 | Early endosome antigen 1 |
| P00441 | SOD1 | -0,218210354 | 5 | 7,69E-05 | 8,28E-05 | Superoxide dismutase [Cu-Zn] |
| P48147 | PREP | -0,217663189 | 10 | 5,04E-08 | 8,96E-08 | Prolyl endopeptidase |
| B4DUC8 | MTAP | -0,217108959 | 4 | 0,000110643 | 0,000115331 | S-methyl-5'-thioadenosine phosphorylase |
| Q12805 | EFEMP1 | -0,21561572 | 4 | 0,014479057 | 0,009360967 | EGF-containing fibulin-like extracellular matrix protein 1 |
| A0A1B0GU03 | NaN | -0,215521074 | 10 | 1,53E-08 | 2,86E-08 | Uncharacterized protein |
| E7EM64 | COPS6 | -0,214391252 | 2 | 0,071343788 | 0,037222846 | COP9 signalosome complex subunit 6 |
| O75531 | BANF1 | -0,214118707 | 2 | 0,031093325 | 0,018409045 | Barrier-to-autointegration factor |
| P54727 | RAD23B | -0,213054404 | 5 | 2,26E-10 | 5,29E-10 | UV excision repair protein RAD23 homolog B |
| F5GWX2 | HEBP1 | -0,212912448 | 3 | 2,56E-06 | 3,59E-06 | Heme-binding protein 1 |
| Q9UIJ7 | AK3 | -0,211892321 | 6 | 3,07E-11 | 8,07E-11 | GTP:AMP phosphotransferase AK3, mitochondrial |
| P07686 | HEXB | -0,209044184 | 6 | 8,46E-09 | 1,66E-08 | Beta-hexosaminidase subunit beta |
| Q96TA1 | FAM129B | -0,2088103 | 17 | 7,21E-07 | 1,09E-06 | Niban-like protein 1 |
| P01008 | SERPINC1 | -0,208769235 | 21 | 4,06E-09 | 8,46E-09 | Antithrombin-III |
| Q16401 | PSMD5 | -0,208767919 | 8 | 1,23E-05 | 1,52E-05 | 26S proteasome non-ATPase regulatory subunit 5 |
| Q6NZI2 | CAVIN1 | -0,207826469 | 8 | 1,89E-07 | 3,13E-07 | Caveolae-associated protein 1 |
| Q13228 | SELENBP1 | -0,207658179 | 17 | 1,21E-08 | 2,30E-08 | Methanethiol oxidase |
| K7ES70 | MFAP4 | -0,206522412 | 4 | 0,003215353 | 0,00246027 | Microfibril-associated glycoprotein 4 |
| Q00577 | PURA | -0,206182167 | 3 | 2,19E-05 | 2,54E-05 | Transcriptional activator protein Pur-alpha |
| C9JF17 | APOD | -0,205635318 | 4 | 5,78E-05 | 6,34E-05 | Apolipoprotein D (Fragment) |
| A8K2U0 | A2ML1 | -0,204819632 | 16 | 2,66E-09 | 5,61E-09 | Alpha-2-macroglobulin-like protein 1 |
| Q96AC1 | FERMT2 | -0,20334193 | 12 | 3,33E-08 | 6,07E-08 | Fermitin family homolog 2 |

| | | | | | | |
|---|---|---|---|---|---|---|
| Q9UHY7 | ENOPH1 | -0,202649175 | 4 | 5,43E-06 | 7,26E-06 | Enolase-phosphatase E1 |
| A0A2R8Y4T1 | TNS1 | -0,202374452 | 9 | 1,03E-06 | 1,54E-06 | Tensin-1 |
| P27169 | PON1 | -0,202055058 | 5 | 0,022373779 | 0,013829019 | Serum paraoxonase/arylesterase 1 |
| P28300 | LOX | -0,200931897 | 5 | 0,013190191 | 0,008572753 | Protein-lysine 6-oxidase |
| P49588 | AARS | -0,200291429 | 9 | 1,62E-05 | 1,95E-05 | Alanine--tRNA ligase, cytoplasmic |
| O95171 | SCEL | -0,200080162 | 10 | 5,59E-06 | 7,43E-06 | Sciellin |
| P09110 | ACAA1 | -0,199699434 | 3 | 0,062544266 | 0,033593645 | 3-ketoacyl-CoA thiolase, peroxisomal |
| P17655 | CAPN2 | -0,199549251 | 13 | 3,28E-05 | 3,70E-05 | Calpain-2 catalytic subunit |
| P20774 | OGN | -0,19940106 | 16 | 3,22E-10 | 7,41E-10 | Mimecan |
| Q9NYL9 | TMOD3 | -0,198674934 | 2 | 1,32E-05 | 1,62E-05 | Tropomodulin-3 |
| Q99733 | NAP1L4 | -0,198295458 | 6 | 4,52E-09 | 9,31E-09 | Nucleosome assembly protein 1-like 4 |
| P62820 | RAB1A | -0,197562255 | 3 | 0,000106162 | 0,000111368 | Ras-related protein Rab-1A |
| P35580 | MYH10 | -0,197428019 | 6 | 4,06E-07 | 6,40E-07 | Myosin-10 |
| A0A0A0MRZ8 | IGKV3D-11 | -0,195956188 | 3 | 0,003178144 | 0,002438262 | Immunoglobulin kappa variable 3D-11 |
| Q02413 | DSG1 | -0,195300191 | 36 | 6,14E-13 | 1,91E-12 | Desmoglein-1 |
| P24752 | ACAT1 | -0,194865198 | 12 | 1,46E-12 | 4,33E-12 | Acetyl-CoA acetyltransferase, mitochondrial |
| Q9C0C2 | TNKS1BP1 | -0,194174291 | 13 | 4,39E-09 | 9,07E-09 | 182 kDa tankyrase-1-binding protein |
| P01619 | IGKV3-20 | -0,192049886 | 5 | 0,006030557 | 0,004337769 | Immunoglobulin kappa variable 3-20 |
| P01834 | IGKC | -0,191883853 | 7 | 6,42E-06 | 8,38E-06 | Immunoglobulin kappa constant |
| Q8IVF2 | AHNAK2 | -0,191556982 | 77 | 6,06E-18 | 2,71E-17 | Protein AHNAK2 |
| A0A286YEY4 | IGHG2 | -0,190888795 | 9 | 0,00246213 | 0,001947537 | Immunoglobulin heavy constant gamma 2 (Fragment) |
| P02647 | APOA1 | -0,190775534 | 27 | 7,44E-11 | 1,85E-10 | Apolipoprotein A-I |
| Q04828 | AKR1C1 | -0,189916804 | 14 | 7,48E-07 | 1,13E-06 | Aldo-keto reductase family 1 member C1 |
| P00167 | CYB5A | -0,189908595 | 2 | 0,001569251 | 0,001297599 | Cytochrome b5 |
| Q92820 | GGH | -0,189524016 | 6 | 0,012997308 | 0,008481002 | Gamma-glutamyl hydrolase |
| P49773 | HINT1 | -0,1886805 | 7 | 0,001548907 | 0,001285096 | Histidine triad nucleotide-binding protein 1 |

| | | | | | | |
|---|---|---|---|---|---|---|
| A0A087WT 99 | C11orf54 | - 0,1883461 52 | 3 | 8,03E-05 | 8,59E-05 | Ester hydrolase C11orf54 |
| Q13057 | COASY | - 0,1882126 38 | 5 | 2,17E-05 | 2,52E-05 | Bifunctional coenzyme A synthase |
| P20933 | AGA | - 0,1870992 53 | 2 | 0,006361033 | 0,0045488 78 | N(4)-(beta-N-acetylglucosaminyl)-L-asparaginase |
| P55263 | ADK | - 0,1869661 68 | 4 | 0,053073286 | 0,0293393 9 | Adenosine kinase |
| A0A087WXI 5 | CDH1 | - 0,1868467 05 | 11 | 1,46E-06 | 2,12E-06 | Cadherin-1 |
| P21964 | COMT | - 0,1866042 07 | 10 | 2,71E-05 | 3,11E-05 | Catechol O-methyltransferase |
| A0A0A0MR L6 | ABLIM1 | - 0,1863942 49 | 3 | 0,034279917 | 0,0201261 57 | Actin-binding LIM protein 1 |
| Q9NRX4 | PHPT1 | - 0,1862807 23 | 3 | 0,003326388 | 0,0025294 94 | 14 kDa phosphohistidine phosphatase |
| Q9NZN4 | EHD2 | - 0,1857847 65 | 25 | 5,79E-22 | 3,28E-21 | EH domain-containing protein 2 |
| P36405 | ARL3 | - 0,1854358 37 | 2 | 0,00737317 | 0,0051259 54 | ADP-ribosylation factor-like protein 3 |
| P12111 | COL6A3 | - 0,1842935 82 | 142 | 2,53E-68 | 5,65E-67 | Collagen alpha-3(VI) chain |
| A0A0C4DH 38 | IGHV5-51 | - 0,1842833 58 | 3 | 0,032310594 | 0,0190609 26 | Immunoglobulin heavy variable 5-51 |
| P30041 | PRDX6 | - 0,1828717 82 | 15 | 0,000739911 | 0,0006665 47 | Peroxiredoxin-6 |
| Q14315 | FLNC | - 0,1828214 53 | 16 | 0,000318548 | 0,0003091 24 | Filamin-C |
| O14896 | IRF6 | - 0,1827542 8 | 4 | 0,021004708 | 0,0131312 79 | Interferon regulatory factor 6 |
| K7EIK7 | EML2 | - 0,1825274 19 | 7 | 1,36E-08 | 2,57E-08 | Echinoderm microtubule-associated protein-like 2 |
| Q96IU4 | ABHD14B | - 0,1825086 62 | 6 | 0,003012024 | 0,0023295 16 | Protein ABHD14B |
| F5GXS0 | C4B | - 0,1823954 27 | 4 | 0,020470907 | 0,0128301 74 | Complement C4-B |
| A0A096LPI 6 | NaN | - 0,1823594 05 | 6 | 0,002388325 | 0,0018983 13 | Uncharacterized protein (Fragment) |
| P02545 | LMNA | - 0,1821476 08 | 51 | 3,88E-40 | 5,45E-39 | Prelamin-A/C |
| Q9BRA2 | TXNDC17 | - 0,1816096 59 | 3 | 0,000107751 | 0,0001127 95 | Thioredoxin domain-containing protein 17 |
| P40926 | MDH2 | - 0,1813817 77 | 15 | 3,04E-22 | 1,78E-21 | Malate dehydrogenase, mitochondrial |
| O15020 | SPTBN2 | - 0,1812003 18 | 42 | 0,0001393 | 0,0001421 9 | Spectrin beta chain, non-erythrocytic 2 |
| P45974 | USP5 | - 0,1811171 01 | 16 | 1,58E-12 | 4,66E-12 | Ubiquitin carboxyl-terminal hydrolase 5 |
| Q9Y6W5 | WASF2 | - 0,1800702 21 | 3 | 0,027558375 | 0,0166160 79 | Wiskott-Aldrich syndrome protein family member 2 |

| | | | | | | |
|---|---|---|---|---|---|---|
| P00734 | F2 | -0,178994326 | 15 | 7,50E-05 | 8,12E-05 | Prothrombin |
| Q9H2U2 | PPA2 | -0,178393576 | 8 | 1,69E-05 | 2,01E-05 | Inorganic pyrophosphatase 2, mitochondrial |
| Q13425 | SNTB2 | -0,178005953 | 3 | 0,004716268 | 0,003463289 | Beta-2-syntrophin |
| Q14247 | CTTN | -0,17798062 | 9 | 5,51E-08 | 9,65E-08 | Src substrate cortactin |
| Q96I99 | SUCLG2 | -0,176804578 | 8 | 9,72E-06 | 1,23E-05 | Succinate--CoA ligase [GDP-forming] subunit beta, mitochondrial |
| P60174 | TPI1 | -0,176774522 | 17 | 8,42E-18 | 3,70E-17 | Triosephosphate isomerase |
| Q9UBT2 | UBA2 | -0,174964183 | 3 | 0,00064629 | 0,000591211 | SUMO-activating enzyme subunit 2 |
| Q13442 | PDAP1 | -0,174368081 | 2 | 2,13E-06 | 3,06E-06 | 28 kDa heat- and acid-stable phosphoprotein |
| P17174 | GOT1 | -0,174324155 | 13 | 4,24E-09 | 8,79E-09 | Aspartate aminotransferase, cytoplasmic |
| Q9BR76 | CORO1B | -0,174156012 | 4 | 0,00416308 | 0,003108096 | Coronin-1B |
| Q15435 | PPP1R7 | -0,174150171 | 6 | 1,14E-08 | 2,19E-08 | Protein phosphatase 1 regulatory subunit 7 |
| P07741 | APRT | -0,174114089 | 9 | 2,34E-10 | 5,45E-10 | Adenine phosphoribosyltransferase |
| E7ENL6 | COL6A3 | -0,173734042 | 2 | 0,003878062 | 0,002921909 | Collagen alpha-3(VI) chain |
| D3YTG3 | ABI3BP | -0,173478145 | 17 | 2,83E-09 | 5,96E-09 | Target of Nesh-SH3 |
| P14923 | JUP | -0,173470035 | 38 | 3,49E-08 | 6,32E-08 | Junction plakoglobin |
| P00450 | CP | -0,173408373 | 41 | 3,57E-06 | 4,97E-06 | Ceruloplasmin |
| B3KUB4 | CA12 | -0,172983242 | 2 | 0,030747746 | 0,018248361 | Carbonic anhydrase 12 |
| A0A087X0S5 | COL6A1 | -0,172384502 | 44 | 9,14E-16 | 3,43E-15 | Collagen alpha-1(VI) chain |
| P10253 | GAA | -0,1720886 | 4 | 0,000135357 | 0,000139031 | Lysosomal alpha-glucosidase |
| P0DOY2 | IGLC2 | -0,17208094 | 6 | 0,001669394 | 0,001368903 | Immunoglobulin lambda constant 2 |
| P10768 | ESD | -0,171895222 | 11 | 0,00095676 | 0,000843595 | S-formylglutathione hydrolase |
| P05091 | ALDH2 | -0,171774271 | 18 | 1,07E-12 | 3,21E-12 | Aldehyde dehydrogenase, mitochondrial |
| A0A1B0GVI3 | KRT10 | -0,170582656 | 5 | 0,014586991 | 0,009406028 | Keratin, type I cytoskeletal 10 (Fragment) |
| G5EA31 | SEC24C | -0,170528598 | 4 | 0,047532722 | 0,026726971 | Protein transport protein Sec24C |
| Q16698 | DECR1 | -0,169665068 | 9 | 1,54E-19 | 7,65E-19 | 2,4-dienoyl-CoA reductase, mitochondrial |
| P33121 | ACSL1 | -0,169423659 | 20 | 0,034702698 | 0,020350092 | Long-chain-fatty-acid--CoA ligase 1 |

| | | | | | | |
|---|---|---|---|---|---|---|
| P55290 | CDH13 | -0,169358185 | 3 | 0,005622471 | 0,004062049 | Cadherin-13 |
| O60701 | UGDH | -0,168910585 | 7 | 0,097696993 | 0,047880263 | UDP-glucose 6-dehydrogenase |
| O00303 | EIF3F | -0,168874551 | 8 | 2,95E-05 | 3,34E-05 | Eukaryotic translation initiation factor 3 subunit F |
| Q9NQR4 | NIT2 | -0,168663508 | 8 | 0,000675425 | 0,000613117 | Omega-amidase NIT2 |
| O75569 | PRKRA | -0,168138304 | 4 | 0,002595763 | 0,00204666 | Interferon-inducible double-stranded RNA-dependent protein kinase activator A |
| P09758 | TACSTD2 | -0,168119701 | 6 | 0,003375129 | 0,002562598 | Tumor-associated calcium signal transducer 2 |
| P00505 | GOT2 | -0,167518005 | 11 | 1,86E-13 | 5,95E-13 | Aspartate aminotransferase, mitochondrial |
| Q15181 | PPA1 | -0,166826746 | 10 | 0,000444968 | 0,000418819 | Inorganic pyrophosphatase |
| Q8N474 | SFRP1 | -0,165604893 | 3 | 0,099167671 | 0,048499497 | Secreted frizzled-related protein 1 |
| A0A3B3ISS6 | GPNMB | -0,165473663 | 4 | 0,009814181 | 0,006614489 | Transmembrane glycoprotein NMB |
| A0A087WSY6 | IGKV3D-15 | -0,165396615 | 3 | 0,001692309 | 0,001385384 | Immunoglobulin kappa variable 3D-15 |
| P12955 | PEPD | -0,165298092 | 5 | 0,017208075 | 0,010938467 | Xaa-Pro dipeptidase |
| Q06830 | PRDX1 | -0,164129385 | 19 | 3,80E-10 | 8,65E-10 | Peroxiredoxin-1 |
| A6NFX8 | NUDT5 | -0,164004701 | 6 | 2,85E-05 | 3,25E-05 | ADP-sugar pyrophosphatase |
| A0A286YFF7 | PPT1 | -0,163612642 | 3 | 0,007323073 | 0,005113616 | Palmitoyl-protein thioesterase 1 |
| I3L2M4 | BAIAP2 | -0,163384882 | 2 | 0,011212801 | 0,007465086 | Brain-specific angiogenesis inhibitor 1-associated protein 2 (Fragment) |
| P37802 | TAGLN2 | -0,163361463 | 10 | 4,24E-06 | 5,80E-06 | Transgelin-2 |
| E7EQR4 | EZR | -0,16306643 | 15 | 1,65E-05 | 1,97E-05 | Ezrin |
| P46939 | UTRN | -0,162046339 | 2 | 0,018549496 | 0,011745627 | Utrophin |
| Q6YHK3 | CD109 | -0,160600268 | 14 | 2,31E-11 | 6,17E-11 | CD109 antigen |
| P50402 | EMD | -0,160301666 | 3 | 0,001399125 | 0,001186844 | Emerin |
| P06396 | GSN | -0,158818311 | 18 | 2,44E-19 | 1,19E-18 | Gelsolin |
| E9PHI4 | SUN1 | -0,158376518 | 6 | 0,000138988 | 0,000142166 | SUN domain-containing protein 1 |
| Q12955 | ANK3 | -0,157720846 | 3 | 0,092795568 | 0,04597726 | Ankyrin-3 |
| A0A1B0GTM3 | ASAH1 | -0,157199029 | 5 | 1,84E-05 | 2,18E-05 | Acid ceramidase |

| | | | | | | |
|---|---|---|---|---|---|---|
| P01701 | IGLV1-51 | -0,157124367 | 3 | 0,011394394 | 0,007565509 | Immunoglobulin lambda variable 1-51 |
| P68402 | PAFAH1B2 | -0,156719198 | 4 | 0,102147143 | 0,049709589 | Platelet-activating factor acetylhydrolase IB subunit beta |
| P62330 | ARF6 | -0,156198088 | 5 | 2,47E-11 | 6,54E-11 | ADP-ribosylation factor 6 |
| P10155 | TROVE2 | -0,156184106 | 6 | 0,005784124 | 0,004166602 | 60 kDa SS-A/Ro ribonucleoprotein |
| P00739 | HPR | -0,155725101 | 4 | 0,041006039 | 0,023323666 | Haptoglobin-related protein |
| P0DPH7 | TUBA3C | -0,155666309 | 3 | 0,020261242 | 0,012714963 | Tubulin alpha-3C chain |
| A0A0A0MS08 | IGHG1 | -0,155440631 | 18 | 2,14E-06 | 3,06E-06 | Immunoglobulin heavy constant gamma 1 (Fragment) |
| D6R9P4 | GNPDA1 | -0,155377643 | 6 | 0,001206581 | 0,001039646 | Glucosamine-6-phosphate isomerase |
| F5GY80 | C8B | -0,155013619 | 6 | 0,067734766 | 0,035566174 | Complement component C8 beta chain |
| A2A274 | ACO2 | -0,154646724 | 20 | 3,23E-25 | 2,15E-24 | Aconitate hydratase, mitochondrial |
| O00264 | PGRMC1 | -0,154300046 | 3 | 0,05253456 | 0,029139801 | Membrane-associated progesterone receptor component 1 |
| P47929 | LGALS7 | -0,154176919 | 10 | 0,003029845 | 0,002336495 | Galectin-7 |
| Q5T6H7 | XPNPEP1 | -0,154084216 | 4 | 0,000819819 | 0,000732035 | Xaa-Pro aminopeptidase 1 |
| P62258 | YWHAE | -0,153969707 | 18 | 2,22E-12 | 6,47E-12 | 14-3-3 protein epsilon |
| P10599 | TXN | -0,153960396 | 7 | 0,013020543 | 0,00848491 | Thioredoxin |
| Q9UHG3 | PCYOX1 | -0,153284506 | 12 | 1,38E-06 | 2,01E-06 | Prenylcysteine oxidase 1 |
| P15088 | CPA3 | -0,153235081 | 8 | 0,025596814 | 0,015605492 | Mast cell carboxypeptidase A |
| Q07507 | DPT | -0,153069352 | 8 | 0,032247982 | 0,019046827 | Dermatopontin |
| Q16658 | FSCN1 | -0,152924368 | 17 | 1,39E-08 | 2,61E-08 | Fascin |
| P02753 | RBP4 | -0,152567672 | 5 | 0,00183593 | 0,001493021 | Retinol-binding protein 4 |
| O15400 | STX7 | -0,152560584 | 3 | 0,034786564 | 0,020357988 | Syntaxin-7 |
| Q9Y5Z4 | HEBP2 | -0,151905978 | 4 | 4,89E-08 | 8,72E-08 | Heme-binding protein 2 |
| P54652 | HSPA2 | -0,151288761 | 7 | 3,67E-09 | 7,68E-09 | Heat shock-related 70 kDa protein 2 |
| O60784 | TOM1 | -0,15095606 | 5 | 0,01275316 | 0,008356368 | Target of Myb protein 1 |
| Q13885 | TUBB2A | -0,150686615 | 6 | 0,010608238 | 0,007101025 | Tubulin beta-2A chain |

| | | | | | | |
|---|---|---|---|---|---|---|
| A0A1W2PNX8 | UNC45A | -0,150672903 | 8 | 0,001432666 | 0,001209043 | Protein unc-45 homolog A |
| P40925 | MDH1 | -0,150658046 | 8 | 0,000762327 | 0,000684425 | Malate dehydrogenase, cytoplasmic |
| A0A3B3ISQ4 | EPS8L2 | -0,150251311 | 4 | 0,002039017 | 0,001644584 | Epidermal growth factor receptor kinase substrate 8-like protein 2 |
| Q06124 | PTPN11 | -0,150064941 | 3 | 0,074734896 | 0,038582968 | Tyrosine-protein phosphatase non-receptor type 11 |
| E9PB61 | ALYREF | -0,149566942 | 6 | 0,018541519 | 0,011745627 | THO complex subunit 4 |
| P02748 | C9 | -0,148810593 | 9 | 0,034917337 | 0,020403005 | Complement component C9 |
| Q9UI09 | NDUFA12 | -0,148791274 | 2 | 0,08649853 | 0,043449514 | NADH dehydrogenase [ubiquinone] 1 alpha subcomplex subunit 12 |
| P16083 | NQO2 | -0,147961188 | 3 | 0,000407588 | 0,00038863 | Ribosyldihydronicotinamide dehydrogenase [quinone] |
| Q3LXA3 | TKFC | -0,147728931 | 6 | 0,000296357 | 0,0002893 | Triokinase/FMN cyclase |
| C9J9W2 | LASP1 | -0,14758328 | 6 | 5,75E-06 | 7,59E-06 | LIM and SH3 domain protein 1 (Fragment) |
| A0A140T9R1 | FLOT1 | -0,146293233 | 6 | 0,000919434 | 0,000815065 | Flotillin-1 (Fragment) |
| P27348 | YWHAQ | -0,145919732 | 8 | 5,01E-07 | 7,78E-07 | 14-3-3 protein theta |
| P07360 | C8G | -0,144932482 | 3 | 0,058179115 | 0,03169892 | Complement component C8 gamma chain |
| E9PHY5 | EPB41L2 | -0,14445557 | 6 | 0,005181533 | 0,003754478 | Band 4.1-like protein 2 |
| Q7Z6Z7 | HUWE1 | -0,143387403 | 6 | 0,000621612 | 0,000573796 | E3 ubiquitin-protein ligase HUWE1 |
| P12532 | CKMT1A | -0,142452725 | 9 | 0,000548012 | 0,000509682 | Creatine kinase U-type, mitochondrial |
| J3KR44 | OTUB1 | -0,142413879 | 11 | 0,001129852 | 0,000980401 | Ubiquitin thioesterase |
| P28072 | PSMB6 | -0,141970546 | 4 | 0,000853467 | 0,000759323 | Proteasome subunit beta type-6 |
| P01700 | IGLV1-47 | -0,141873453 | 2 | 0,074054659 | 0,038271945 | Immunoglobulin lambda variable 1-47 |
| P29622 | SERPINA4 | -0,14175025 | 11 | 5,55E-05 | 6,10E-05 | Kallistatin |
| Q96KP4 | CNDP2 | -0,141737468 | 16 | 3,98E-07 | 6,31E-07 | Cytosolic non-specific dipeptidase |
| H3BLU7 | AKR7A2 | -0,141724335 | 4 | 0,025665663 | 0,015628102 | Aflatoxin B1 aldehyde reductase member 2 (Fragment) |
| P51648 | ALDH3A2 | -0,140545086 | 9 | 0,004122248 | 0,003082289 | Fatty aldehyde dehydrogenase |
| Q8N1G4 | LRRC47 | -0,140376554 | 6 | 8,57E-06 | 1,09E-05 | Leucine-rich repeat-containing protein 47 |
| Q96HC4 | PDLIM5 | -0,140347211 | 5 | 0,000424817 | 0,000403494 | PDZ and LIM domain protein 5 |

| | | | | | | |
|---|---|---|---|---|---|---|
| Q92506 | HSD17B8 | -0,139333231 | 4 | 0,029055804 | 0,017391065 | Estradiol 17-beta-dehydrogenase 8 |
| B4DJV2 | CS | -0,13838299 | 12 | 0,003473085 | 0,002632909 | Citrate synthase |
| Q07954 | LRP1 | -0,138286992 | 9 | 0,011192124 | 0,007461416 | Prolow-density lipoprotein receptor-related protein 1 |
| Q13409-2 | DYNC1I2 | -0,137505333 | 9 | 5,88E-05 | 6,44E-05 | Isoform 2B of Cytoplasmic dynein 1 intermediate chain 2 |
| Q15056 | EIF4H | -0,1373224 | 5 | 0,006084497 | 0,004363808 | Eukaryotic translation initiation factor 4H |
| P35858 | IGFALS | -0,136611288 | 6 | 0,009932365 | 0,006684984 | Insulin-like growth factor-binding protein complex acid labile subunit |
| Q13509 | TUBB3 | -0,136468765 | 6 | 0,035152529 | 0,020491759 | Tubulin beta-3 chain |
| P68371 | TUBB4B | -0,136354704 | 18 | 0,00133475 | 0,001138123 | Tubulin beta-4B chain |
| P00568 | AK1 | -0,136329295 | 5 | 0,067353577 | 0,035457643 | Adenylate kinase isoenzyme 1 |
| A0A087X253 | AP2B1 | -0,135889347 | 6 | 0,101001757 | 0,049200856 | AP complex subunit beta |
| P31947 | SFN | -0,135216139 | 17 | 4,69E-05 | 5,20E-05 | 14-3-3 protein sigma |
| P11216 | PYGB | -0,135165226 | 32 | 0,012608813 | 0,008282425 | Glycogen phosphorylase, brain form |
| P27338 | MAOB | -0,134820957 | 5 | 0,01217916 | 0,00804315 | Amine oxidase [flavin-containing] B |
| P22314 | UBA1 | -0,134248006 | 33 | 9,51E-13 | 2,89E-12 | Ubiquitin-like modifier-activating enzyme 1 |
| Q9NSE4 | IARS2 | -0,133994783 | 6 | 0,000277234 | 0,000272254 | Isoleucine--tRNA ligase, mitochondrial |
| Q9H223 | EHD4 | -0,133804969 | 11 | 4,16E-06 | 5,70E-06 | EH domain-containing protein 4 |
| P98095 | FBLN2 | -0,133758207 | 17 | 4,04E-07 | 6,39E-07 | Fibulin-2 |
| A0A0G2JPR0 | C4A | -0,133747191 | 60 | 7,06E-05 | 7,66E-05 | Complement C4-A |
| Q14166 | TTLL12 | -0,133423065 | 6 | 0,002727861 | 0,002137763 | Tubulin--tyrosine ligase-like protein 12 |
| A0A1W2PPU6 | SCARB2 | -0,132362139 | 4 | 0,064884855 | 0,034437269 | Lysosome membrane protein 2 |
| Q13011 | ECH1 | -0,132290918 | 9 | 0,031041574 | 0,018400547 | Delta(3,5)-Delta(2,4)-dienoyl-CoA isomerase, mitochondrial |
| Q99714 | HSD17B10 | -0,132168929 | 15 | 0,000286218 | 0,000280516 | 3-hydroxyacyl-CoA dehydrogenase type-2 |
| E7EX90 | DCTN1 | -0,131815647 | 10 | 6,41E-08 | 1,11E-07 | Dynactin subunit 1 |
| A1L0T0 | ILVBL | -0,131785614 | 7 | 0,001182828 | 0,001020967 | Acetolactate synthase-like protein |
| J3QRN6 | MYO1D | -0,131509196 | 9 | 0,007376373 | 0,005125954 | Unconventional myosin-Id |
| P50213 | IDH3A | -0,131257232 | 5 | 2,09E-05 | 2,43E-05 | Isocitrate dehydrogenase [NAD] subunit alpha, mitochondrial |

| | | | | | | |
|---|---|---|---|---|---|---|
| P31939 | ATIC | -0,130941615 | 14 | 6,49E-06 | 8,45E-06 | Bifunctional purine biosynthesis protein PURH |
| P00367 | GLUD1 | -0,130838434 | 23 | 1,29E-06 | 1,89E-06 | Glutamate dehydrogenase 1, mitochondrial |
| P61970 | NUTF2 | -0,13066509 | 4 | 0,053269579 | 0,029359379 | Nuclear transport factor 2 |
| P34896 | SHMT1 | -0,130588812 | 3 | 0,004684532 | 0,003445127 | Serine hydroxymethyltransferase, cytosolic |
| M0R261 | PGLS | -0,129196974 | 4 | 0,09932014 | 0,048525828 | 6-phosphogluconolactonase (Fragment) |
| A0A0B4J1V0 | IGHV3-15 | -0,129003165 | 2 | 0,092666572 | 0,04595963 | Immunoglobulin heavy variable 3-15 |
| P49411 | TUFM | -0,127566136 | 14 | 7,95E-05 | 8,55E-05 | Elongation factor Tu, mitochondrial |
| E7EPK1 | SEPTIN7 | -0,127394169 | 10 | 1,00E-09 | 2,20E-09 | Septin-7 |
| P53597 | SUCLG1 | -0,127331509 | 3 | 0,002763069 | 0,002157826 | Succinate--CoA ligase [ADP/GDP-forming] subunit alpha, mitochondrial |
| C9JFR7 | CYCS | -0,127205696 | 5 | 0,000670966 | 0,000610195 | Cytochrome c (Fragment) |
| A0A024R442 | DNPEP | -0,127039198 | 6 | 0,041241683 | 0,023430609 | Aspartyl aminopeptidase, isoform CRA_b |
| P08559 | PDHA1 | -0,126989114 | 4 | 0,084488025 | 0,042677729 | Pyruvate dehydrogenase E1 component subunit alpha, somatic form, mitochondrial |
| Q92747 | ARPC1A | -0,126716058 | 6 | 0,040556395 | 0,023148198 | Actin-related protein 2/3 complex subunit 1A |
| A0A0C4DGB5 | CAST | -0,125811688 | 18 | 6,45E-09 | 1,29E-08 | Calpastatin |
| P12110 | COL6A2 | -0,125243229 | 36 | 0,001460611 | 0,001224226 | Collagen alpha-2(VI) chain |
| Q01082 | SPTBN1 | -0,12389569 | 75 | 1,47E-17 | 6,28E-17 | Spectrin beta chain, non-erythrocytic 1 |
| P09417 | QDPR | -0,122118932 | 6 | 0,001292467 | 0,001105902 | Dihydropteridine reductase |
| Q9BRX8 | PRXL2A | -0,121580625 | 3 | 0,05163756 | 0,028772004 | Peroxiredoxin-like 2A |
| P68363 | TUBA1B | -0,121381984 | 27 | 0,000179346 | 0,000179711 | Tubulin alpha-1B chain |
| P15924 | DSP | -0,121175727 | 189 | 7,93E-11 | 1,95E-10 | Desmoplakin |
| Q15019 | SEPTIN2 | -0,120625422 | 13 | 5,38E-06 | 7,22E-06 | Septin-2 |
| Q14574 | DSC3 | -0,12049878 | 21 | 0,021035698 | 0,013133964 | Desmocollin-3 |
| P30084 | ECHS1 | -0,119742838 | 10 | 8,91E-11 | 2,17E-10 | Enoyl-CoA hydratase, mitochondrial |
| A0A087WYR3 | TPD52L2 | -0,119603817 | 2 | 0,006710535 | 0,004743654 | Tumor protein D54 |
| O43852 | CALU | -0,11952921 | 4 | 0,024401397 | 0,014976298 | Calumenin |

| | | | | | | |
|---|---|---|---|---|---|---|
| A0A024R571 | EHD1 | -0,119514833 | 14 | 5,55E-12 | 1,54E-11 | EH domain-containing protein 1 |
| P01024 | C3 | -0,118681403 | 95 | 2,04E-09 | 4,37E-09 | Complement C3 |
| Q9GZP4 | PITHD1 | -0,118656488 | 3 | 0,052179964 | 0,028975781 | PITH domain-containing protein 1 |
| A0A0A0MS15 | IGHV3-49 | -0,118220176 | 3 | 0,076404127 | 0,039320953 | Immunoglobulin heavy variable 3-49 |
| A0A3B3ISX9 | TNXB | -0,117771875 | 114 | 1,43E-34 | 1,56E-33 | Tenascin-X |
| K7EKI8 | PPL | -0,117520546 | 51 | 1,44E-14 | 4,93E-14 | Periplakin |
| Q8NBS9 | TXNDC5 | -0,117484963 | 8 | 0,03891398 | 0,022392606 | Thioredoxin domain-containing protein 5 |
| Q99536 | VAT1 | -0,116685505 | 17 | 1,86E-05 | 2,19E-05 | Synaptic vesicle membrane protein VAT-1 homolog |
| E7EXA3 | NIF3L1 | -0,116452593 | 2 | 0,027703186 | 0,016682947 | NIF3-like protein 1 (Fragment) |
| P07355 | ANXA2 | -0,116423845 | 28 | 1,28E-08 | 2,42E-08 | Annexin A2 |
| A0A0C4DGQ5 | CAPNS1 | -0,115260856 | 7 | 0,021242104 | 0,013246027 | Calpain small subunit 1 |
| P61086 | UBE2K | -0,115215719 | 3 | 0,084256018 | 0,042604276 | Ubiquitin-conjugating enzyme E2 K |
| A0A024R4E5 | HDLBP | -0,114506021 | 13 | 0,000436264 | 0,000412773 | High density lipoprotein binding protein (Vigilin), isoform CRA_a |
| P09211 | GSTP1 | -0,113793088 | 12 | 2,61E-07 | 4,24E-07 | Glutathione S-transferase P |
| A0A2R8Y543 | CTNNB1 | -0,113723075 | 23 | 0,009076255 | 0,006176373 | Catenin beta-1 |
| P36957 | DLST | -0,113417444 | 8 | 5,96E-06 | 7,84E-06 | Dihydrolipoyllysine-residue succinyltransferase component of 2-oxoglutarate dehydrogenase complex, mitochondrial |
| K7EN45 | PIN1 | -0,113352899 | 2 | 0,067701291 | 0,035566174 | Peptidyl-prolyl cis-trans isomerase (Fragment) |
| P28838 | LAP3 | -0,113067263 | 16 | 0,002601326 | 0,002047764 | Cytosol aminopeptidase |
| P10301 | RRAS | -0,112872602 | 4 | 0,012571858 | 0,008270231 | Ras-related protein R-Ras |
| B4DJ81 | NDUFS1 | -0,111638737 | 13 | 3,38E-05 | 3,80E-05 | NADH-ubiquinone oxidoreductase 75 kDa subunit, mitochondrial |
| A0A087WW40 | SH3GLB1 | -0,11150432 | 2 | 0,092218626 | 0,045783617 | Endophilin-B1 |
| Q13596 | SNX1 | -0,111450747 | 5 | 3,01E-08 | 5,51E-08 | Sorting nexin-1 |
| A6NKB8 | RNPEP | -0,111089836 | 12 | 2,37E-08 | 4,38E-08 | Aminopeptidase B |
| P02649 | APOE | -0,111027803 | 10 | 0,000314751 | 0,000306042 | Apolipoprotein E |
| O75368 | SH3BGRL | -0,110980064 | 6 | 0,004118439 | 0,003082289 | SH3 domain-binding glutamic acid-rich-like protein |

| | | | | | | |
|---|---|---|---|---|---|---|
| P49748 | ACADVL | -0,110366598 | 16 | 3,52E-15 | 1,23E-14 | Very long-chain specific acyl-CoA dehydrogenase, mitochondrial |
| A0A0G2JMB2 | IGHA2 | -0,110067298 | 10 | 0,040494996 | 0,023139998 | Immunoglobulin heavy constant alpha 2 (Fragment) |
| Q6YN16 | HSDL2 | -0,110055315 | 4 | 0,008753029 | 0,005989555 | Hydroxysteroid dehydrogenase-like protein 2 |
| F5H6E2 | MYO1C | -0,109848797 | 26 | 3,79E-05 | 4,26E-05 | Unconventional myosin-Ic |
| O00151 | PDLIM1 | -0,109393145 | 9 | 0,00219978 | 0,00176269 | PDZ and LIM domain protein 1 |
| Q92597 | NDRG1 | -0,109347192 | 11 | 0,000131925 | 0,000136647 | Protein NDRG1 |
| Q01813 | PFKP | -0,10885937 | 7 | 0,005189117 | 0,003754478 | ATP-dependent 6-phosphofructokinase, platelet type |
| Q9H845 | ACAD9 | -0,108790197 | 8 | 0,00242558 | 0,001924815 | Acyl-CoA dehydrogenase family member 9, mitochondrial |
| H0YLU7 | ETFA | -0,10871974 | 9 | 1,17E-06 | 1,73E-06 | Electron transfer flavoprotein subunit alpha, mitochondrial (Fragment) |
| Q9Y446 | PKP3 | -0,108558573 | 26 | 0,064029236 | 0,034130427 | Plakophilin-3 |
| P00488 | F13A1 | -0,108417056 | 20 | 1,88E-05 | 2,21E-05 | Coagulation factor XIII A chain |
| Q9UKG1 | APPL1 | -0,107340197 | 5 | 0,085437239 | 0,043068772 | DCC-interacting protein 13-alpha |
| P07954 | FH | -0,107130282 | 11 | 0,004795116 | 0,003515942 | Fumarate hydratase, mitochondrial |
| Q13561 | DCTN2 | -0,107019755 | 8 | 0,002689704 | 0,002113953 | Dynactin subunit 2 |
| O15173 | PGRMC2 | -0,106948712 | 3 | 0,000663328 | 0,000604365 | Membrane-associated progesterone receptor component 2 |
| Q86U42 | PABPN1 | -0,106667895 | 3 | 0,050198605 | 0,028097513 | Polyadenylate-binding protein 2 |
| Q05707 | COL14A1 | -0,1064308 | 50 | 1,30E-07 | 2,17E-07 | Collagen alpha-1(XIV) chain |
| K7EQA1 | PDCD5 | -0,106430454 | 2 | 0,091817255 | 0,045700347 | Programmed cell death protein 5 |
| P58107 | EPPK1 | -0,106297837 | 110 | 0,022868133 | 0,014084561 | Epiplakin |
| Q8WUF5 | PPP1R13L | -0,106176245 | 5 | 0,063743615 | 0,034015031 | RelA-associated inhibitor |
| Q9H0U4 | RAB1B | -0,106102571 | 7 | 8,12E-07 | 1,23E-06 | Ras-related protein Rab-1B |
| Q15582 | TGFBI | -0,105726498 | 24 | 0,001125281 | 0,00097816 | Transforming growth factor-beta-induced protein ig-h3 |
| P63104 | YWHAZ | -0,105201448 | 18 | 1,09E-08 | 2,09E-08 | 14-3-3 protein zeta/delta |
| P30838 | ALDH3A1 | -0,104573169 | 10 | 9,92E-08 | 1,69E-07 | Aldehyde dehydrogenase, dimeric NADP-preferring |
| P63241 | EIF5A | -0,103685547 | 9 | 0,099065494 | 0,048497734 | Eukaryotic translation initiation factor 5A-1 |
| P62937 | PPIA | -0,102837469 | 13 | 0,042500124 | 0,024117718 | Peptidyl-prolyl cis-trans isomerase A |

| | | | | | | |
|---|---|---|---|---|---|---|
| A0A087WYF7 | APOOL | -0,10274002 | 2 | 0,027841183 | 0,016725106 | MICOS complex subunit |
| A0A087WX97 | BCL2L13 | -0,102639619 | 2 | 0,086633737 | 0,043449514 | Bcl-2-like protein 13 |
| Q9GZZ9 | UBA5 | -0,102568322 | 3 | 0,039604217 | 0,022763171 | Ubiquitin-like modifier-activating enzyme 5 |
| P04181 | OAT | -0,102316235 | 4 | 0,096086097 | 0,047369098 | Ornithine aminotransferase, mitochondrial |
| E9PLK3 | NPEPPS | -0,101886662 | 14 | 2,98E-07 | 4,80E-07 | Aminopeptidase |
| O43707 | ACTN4 | -0,100952619 | 34 | 0,007832294 | 0,005404613 | Alpha-actinin-4 |
| B7ZKJ8 | ITIH4 | -0,10084745 | 25 | 0,002204194 | 0,001763355 | ITIH4 protein |
| Q969G5 | CAVIN3 | -0,100783821 | 7 | 0,001953237 | 0,00158058 | Caveolae-associated protein 3 |
| A0A075B6R9 | IGKV2D-24 | -0,100313341 | 2 | 0,067312274 | 0,035457643 | Immunoglobulin kappa variable 2D-24 (non-functional) (Fragment) |
| D6RAX7 | COPS4 | -0,100100697 | 3 | 0,097192807 | 0,047802759 | COP9 constitutive photomorphogenic-like protein subunit 4 isoform 2 |
| Q13813 | SPTAN1 | -0,099271819 | 103 | 4,39E-29 | 3,54E-28 | Spectrin alpha chain, non-erythrocytic 1 |
| O60664 | PLIN3 | -0,099216393 | 13 | 0,00301606 | 0,002329516 | Perilipin-3 |
| G3V0I5 | NDUFV1 | -0,099107725 | 10 | 0,05182187 | 0,028809446 | NADH dehydrogenase [ubiquinone] flavoprotein 1, mitochondrial |
| O75306 | NDUFS2 | -0,099042651 | 5 | 0,006545767 | 0,004647211 | NADH dehydrogenase [ubiquinone] iron-sulfur protein 2, mitochondrial |
| Q03135 | CAV1 | -0,098793356 | 4 | 0,070493597 | 0,036870638 | Caveolin-1 |
| P15374 | UCHL3 | -0,097487943 | 5 | 0,070518841 | 0,036870638 | Ubiquitin carboxyl-terminal hydrolase isozyme L3 |
| C9JJP5 | TFG | -0,097218309 | 4 | 0,024578372 | 0,015059227 | Protein TFG (Fragment) |
| P07437 | TUBB | -0,097135758 | 5 | 0,07327607 | 0,038029353 | Tubulin beta chain |
| A0MZ66 | SHTN1 | -0,097133376 | 2 | 0,018217332 | 0,011565067 | Shootin-1 |
| Q8WUM4 | PDCD6IP | -0,09668976 | 15 | 0,000133928 | 0,000137851 | Programmed cell death 6-interacting protein |
| P00387 | CYB5R3 | -0,09615426 | 14 | 0,004217894 | 0,003139492 | NADH-cytochrome b5 reductase 3 |
| E9PQP6 | FNTA | -0,096136484 | 4 | 0,007016394 | 0,004931523 | Protein farnesyltransferase/geranylgeranyltransferase type-1 subunit alpha (Fragment) |
| P13861 | PRKAR2A | -0,09596438 | 8 | 1,24E-06 | 1,83E-06 | cAMP-dependent protein kinase type II-alpha regulatory subunit |
| F6QDS0 | hCG_2043426 | -0,094983299 | 4 | 0,003311016 | 0,002521703 | HCG2043426, isoform CRA_b |
| P13671 | C6 | -0,093971509 | 3 | 0,071959867 | 0,037504507 | Complement component C6 |

| | | | | | | |
|---|---|---|---|---|---|---|
| Q14980 | NUMA1 | -0,093965786 | 22 | 0,002320401 | 0,001850303 | Nuclear mitotic apparatus protein 1 |
| P78347 | GTF2I | -0,093331804 | 7 | 0,040244935 | 0,023023847 | General transcription factor II-I |
| O00204 | SULT2B1 | -0,092841623 | 5 | 0,00046792 | 0,000437674 | Sulfotransferase family cytosolic 2B member 1 |
| P02549 | SPTA1 | -0,092436074 | 57 | 5,26E-07 | 8,11E-07 | Spectrin alpha chain, erythrocytic 1 |
| P06756 | ITGAV | -0,089895657 | 2 | 0,090519434 | 0,045182902 | Integrin alpha-V |
| P05455 | SSB | -0,089866075 | 8 | 0,019452857 | 0,012286015 | Lupus La protein |
| O75746 | SLC25A12 | -0,089604566 | 8 | 0,000433456 | 0,000410907 | Calcium-binding mitochondrial carrier protein Aralar1 |
| I3L1P8 | SLC25A11 | -0,089214554 | 5 | 0,100592889 | 0,049050249 | Mitochondrial 2-oxoglutarate/malate carrier protein (Fragment) |
| O95479 | H6PD | -0,088554938 | 5 | 0,00062013 | 0,000573504 | GDH/6PGL endoplasmic bifunctional protein |
| Q9NZL9 | MAT2B | -0,088273142 | 3 | 0,000137341 | 0,000140774 | Methionine adenosyltransferase 2 subunit beta |
| P55084 | HADHB | -0,08805678 | 16 | 7,28E-13 | 2,24E-12 | Trifunctional enzyme subunit beta, mitochondrial |
| Q15149-4 | PLEC | -0,086477535 | 174 | 0,000688305 | 0,000622511 | Isoform 4 of Plectin |
| O94973 | AP2A2 | -0,086325419 | 11 | 0,04011292 | 0,023001815 | AP-2 complex subunit alpha-2 |
| Q86VP6 | CAND1 | -0,085991222 | 20 | 0,06347081 | 0,034015031 | Cullin-associated NEDD8-dissociated protein 1 |
| P08758 | ANXA5 | -0,085762499 | 18 | 0,001044083 | 0,000914037 | Annexin A5 |
| J3KQ32 | OLA1 | -0,08475982 | 5 | 0,054371007 | 0,029922299 | Obg-like ATPase 1 |
| Q96CX2 | KCTD12 | -0,084523327 | 8 | 0,002449843 | 0,001940938 | BTB/POZ domain-containing protein KCTD12 |
| C9JGI3 | TYMP | -0,084150561 | 11 | 0,014573282 | 0,009406028 | Thymidine phosphorylase (Fragment) |
| Q9NR45 | NANS | -0,08282405 | 7 | 0,058635719 | 0,031877098 | Sialic acid synthase |
| P11177 | PDHB | -0,082440483 | 9 | 0,000167919 | 0,000168949 | Pyruvate dehydrogenase E1 component subunit beta, mitochondrial |
| Q03252 | LMNB2 | -0,081163398 | 14 | 0,087157494 | 0,043667502 | Lamin-B2 |
| Q99961 | SH3GL1 | -0,080512108 | 3 | 0,000165913 | 0,000167273 | Endophilin-A2 |
| E7EMB3 | CALM2 | -0,080382976 | 7 | 0,000161193 | 0,000163519 | Calmodulin-2 |
| P41250 | GARS | -0,079681916 | 9 | 0,012573441 | 0,008270231 | Glycine--tRNA ligase |
| Q9UNZ2 | NSFL1C | -0,079412762 | 6 | 3,42E-08 | 6,21E-08 | NSFL1 cofactor p47 |

| | | | | | | |
|---|---|---|---|---|---|---|
| P04275 | VWF | -0,078091387 | 7 | 0,051764216 | 0,028809446 | von Willebrand factor |
| O43776 | NARS | -0,078075143 | 10 | 0,000400672 | 0,000383523 | Asparagine--tRNA ligase, cytoplasmic |
| P22695 | UQCRC2 | -0,077471762 | 15 | 0,002871247 | 0,00223168 | Cytochrome b-c1 complex subunit 2, mitochondrial |
| Q96G03 | PGM2 | -0,077385327 | 12 | 0,018849293 | 0,011920119 | Phosphoglucomutase-2 |
| Q15063-3 | POSTN | -0,077078882 | 24 | 2,64E-07 | 4,28E-07 | Isoform 3 of Periostin |
| Q14011 | CIRBP | -0,076987341 | 5 | 0,00114374 | 0,000990704 | Cold-inducible RNA-binding protein |
| P43034 | PAFAH1B1 | -0,076969564 | 6 | 0,003651149 | 0,002763639 | Platelet-activating factor acetylhydrolase IB subunit alpha |
| A0A286YEY1 | IGHA1 | -0,076270134 | 5 | 0,082825797 | 0,042140943 | Immunoglobulin heavy constant alpha 1 (Fragment) |
| Q13363 | CTBP1 | -0,075698373 | 3 | 0,083682101 | 0,042401229 | C-terminal-binding protein 1 |
| P30044 | PRDX5 | -0,074574199 | 10 | 0,000398801 | 0,000382476 | Peroxiredoxin-5, mitochondrial |
| P25705 | ATP5F1A | -0,073210323 | 23 | 1,42E-05 | 1,72E-05 | ATP synthase subunit alpha, mitochondrial |
| Q9P2B2 | PTGFRN | -0,072170795 | 2 | 0,035021225 | 0,020439434 | Prostaglandin F2 receptor negative regulator |
| Q9BUF5 | TUBB6 | -0,072022986 | 11 | 0,037765141 | 0,021790621 | Tubulin beta-6 chain |
| P18085 | ARF4 | -0,070994565 | 5 | 0,006531114 | 0,00464567 | ADP-ribosylation factor 4 |
| Q13564 | NAE1 | -0,070542337 | 4 | 0,097464727 | 0,047856932 | NEDD8-activating enzyme E1 regulatory subunit |
| Q9NZM1 | MYOF | -0,069540785 | 16 | 0,016785008 | 0,010697181 | Myoferlin |
| P54709 | ATP1B3 | -0,069127368 | 8 | 0,046361913 | 0,026098468 | Sodium/potassium-transporting ATPase subunit beta-3 |
| P31150 | GDI1 | -0,068486587 | 8 | 0,001415166 | 0,001198385 | Rab GDP dissociation inhibitor alpha |
| E9PCR7 | OGDH | -0,067842729 | 17 | 0,029838318 | 0,017751454 | 2-oxoglutarate dehydrogenase, mitochondrial |
| Q96LJ7 | DHRS1 | -0,066515908 | 4 | 0,051192338 | 0,02855627 | Dehydrogenase/reductase SDR family member 1 |
| A0A1B0GUU9 | IGHM | -0,065346104 | 16 | 0,025424469 | 0,01553893 | Immunoglobulin heavy constant mu (Fragment) |
| P30038 | ALDH4A1 | -0,059976109 | 5 | 0,002556279 | 0,002018763 | Delta-1-pyrroline-5-carboxylate dehydrogenase, mitochondrial |
| P48735 | IDH2 | -0,059263471 | 16 | 0,047775163 | 0,026832626 | Isocitrate dehydrogenase [NADP], mitochondrial |
| G3V4U0 | FBLN5 | -0,055716544 | 5 | 0,058957987 | 0,031966107 | Fibulin-5 |
| A0A0B4J231 | IGLL5 | -0,055426614 | 3 | 0,040731591 | 0,023221255 | Immunoglobulin lambda-like polypeptide 5 |

| | | | | | | |
|---|---|---|---|---|---|---|
| A0A087X2I1 | PSMC6 | -0,054107301 | 6 | 0,065781795 | 0,034875693 | 26S proteasome regulatory subunit 10B |
| D6REX3 | SEC31A | -0,053857525 | 8 | 0,090641309 | 0,045182902 | Protein transport protein Sec31A |
| P04080 | CSTB | -0,051442296 | 4 | 0,05453679 | 0,029980001 | Cystatin-B |
| B9A067 | IMMT | -0,05115634 | 10 | 0,025822057 | 0,015703896 | MICOS complex subunit MIC60 |
| P50502 | ST13 | -0,050131338 | 6 | 0,00736601 | 0,005125954 | Hsc70-interacting protein |
| H0YFD6 | HADHA | -0,04988804 | 23 | 0,025290875 | 0,015476506 | Trifunctional enzyme subunit alpha, mitochondrial |
| P05387 | RPLP2 | -0,049458314 | 7 | 0,010714826 | 0,007162628 | 60S acidic ribosomal protein P2 |
| O75369 | FLNB | -0,047593231 | 75 | 0,086634092 | 0,043449514 | Filamin-B |
| P08603 | CFH | -0,047062734 | 41 | 0,080348902 | 0,04096545 | Complement factor H |
| P22234 | PAICS | -0,046458666 | 6 | 0,095675254 | 0,047235448 | Multifunctional protein ADE2 |
| P06576 | ATP5F1B | -0,044758548 | 28 | 0,007298409 | 0,005107848 | ATP synthase subunit beta, mitochondrial |
| Q9UJZ1 | STOML2 | -0,044746346 | 4 | 0,077307435 | 0,03962006 | Stomatin-like protein 2, mitochondrial |
| P25786 | PSMA1 | -0,043280156 | 8 | 0,029560617 | 0,017650271 | Proteasome subunit alpha type-1 |
| A0A0G2JIW1 | HSPA1B | -0,04254013 | 28 | 0,044939956 | 0,02541432 | Heat shock 70 kDa protein 1B |
| P23526 | AHCY | -0,039635236 | 10 | 0,072822864 | 0,037874048 | Adenosylhomocysteinase |
| O14818 | PSMA7 | -0,037619013 | 8 | 0,064367603 | 0,034236606 | Proteasome subunit alpha type-7 |
| P30043 | BLVRB | -0,034934407 | 13 | 0,016857891 | 0,010729731 | Flavin reductase (NADPH) |
| A0A286YES1 | IGHG3 | -0,034687934 | 9 | 0,005187758 | 0,003754478 | Immunoglobulin heavy constant gamma 3 (Fragment) |
| P54136 | RARS | -0,03187902 | 8 | 0,075716702 | 0,039007976 | Arginine--tRNA ligase, cytoplasmic |
| Q14204 | DYNC1H1 | -0,029257383 | 89 | 0,001778131 | 0,001450814 | Cytoplasmic dynein 1 heavy chain 1 |
| D6RGG3 | COL12A1 | -0,027894755 | 33 | 1,74E-09 | 3,76E-09 | Collagen alpha-1(XII) chain |
| A0A0A0MR02 | VDAC2 | -0,025186907 | 9 | 0,085316508 | 0,043052023 | Voltage-dependent anion-selective channel protein 2 (Fragment) |
| A0A087WZN1 | IDH3B | -0,021344715 | 4 | 0,024262275 | 0,014921299 | Isocitrate dehydrogenase [NAD] subunit, mitochondrial |
| P04114 | APOB | -0,017950608 | 148 | 8,59E-18 | 3,74E-17 | Apolipoprotein B-100 |
| F5H2B9 | UACA | -0,010826532 | 2 | 0,061772338 | 0,033288051 | Uveal autoantigen with coiled-coil domains and ankyrin repeats |

| | | | | | | |
|---|---|---|---|---|---|---|
| P05141 | SLC25A5 | -0,008264416 | 7 | 0,064172824 | 0,034169945 | ADP/ATP translocase 2 |
| P19367 | HK1 | 0,00117115 | 21 | 0,044592936 | 0,025276181 | Hexokinase-1 |
| A0A2R8Y891 | PFKM | 0,001258373 | 7 | 0,072982254 | 0,037916863 | ATP-dependent 6-phosphofructokinase |
| P62736 | ACTA2 | 0,006523831 | 16 | 0,036256881 | 0,021035832 | Actin, aortic smooth muscle |
| P00558 | PGK1 | 0,014565491 | 22 | 0,022550967 | 0,013921049 | Phosphoglycerate kinase 1 |
| P14618 | PKM | 0,021113733 | 33 | 2,94E-06 | 4,12E-06 | Pyruvate kinase PKM |
| P14625 | HSP90B1 | 0,023744766 | 27 | 0,004315819 | 0,003202689 | Endoplasmin |
| P00918 | CA2 | 0,024347229 | 16 | 0,006573495 | 0,004660172 | Carbonic anhydrase 2 |
| P18669 | PGAM1 | 0,025152206 | 16 | 0,008996161 | 0,00613885 | Phosphoglycerate mutase 1 |
| P40121 | CAPG | 0,027268827 | 12 | 0,001441649 | 0,001212792 | Macrophage-capping protein |
| P61088 | UBE2N | 0,028042406 | 6 | 0,067383936 | 0,035457643 | Ubiquitin-conjugating enzyme E2 N |
| K7ELL7 | PRKCSH | 0,028093335 | 6 | 0,077158001 | 0,03962006 | Glucosidase 2 subunit beta |
| P09972 | ALDOC | 0,028285115 | 6 | 0,022305212 | 0,013821366 | Fructose-bisphosphate aldolase C |
| A0A087WVQ6 | CLTC | 0,028930256 | 71 | 0,084159321 | 0,042599163 | Clathrin heavy chain |
| H0YL69 | PSMA4 | 0,030307912 | 8 | 0,000713556 | 0,000644164 | Proteasome subunit alpha type (Fragment) |
| P12236 | SLC25A6 | 0,030328322 | 9 | 0,007647751 | 0,005307043 | ADP/ATP translocase 3 |
| Q15029 | EFTUD2 | 0,031753888 | 8 | 0,006702948 | 0,004743654 | 116 kDa U5 small nuclear ribonucleoprotein component |
| P62829 | RPL23 | 0,034409554 | 5 | 0,071054798 | 0,037111423 | 60S ribosomal protein L23 |
| P55072 | VCP | 0,034453025 | 31 | 0,01647685 | 0,01051441 | Transitional endoplasmic reticulum ATPase |
| Q6XQN6 | NAPRT | 0,034499144 | 11 | 0,012988129 | 0,008481002 | Nicotinate phosphoribosyltransferase |
| A0A087X0X3 | HNRNPM | 0,035262927 | 19 | 0,064736024 | 0,034395382 | Heterogeneous nuclear ribonucleoprotein M |
| P16615 | ATP2A2 | 0,035308101 | 13 | 1,43E-05 | 1,73E-05 | Sarcoplasmic/endoplasmic reticulum calcium ATPase 2 |
| P06733 | ENO1 | 0,03568959 | 24 | 0,001482238 | 0,001238134 | Alpha-enolase |
| P50990 | CCT8 | 0,03591272 | 19 | 2,15E-06 | 3,06E-06 | T-complex protein 1 subunit theta |
| H3BPE1 | MACF1 | 0,038431223 | 14 | 0,013362763 | 0,008673456 | Microtubule-actin cross-linking factor 1, isoforms 1/2/3/5 |
| Q9UHD8 | SEPTIN9 | 0,03883977 | 10 | 0,002183039 | 0,001752129 | Septin-9 |
| B0QY89 | EIF3L | 0,039610175 | 11 | 0,082937824 | 0,042154349 | Eukaryotic translation initiation factor 3 subunit L |
| F8VVA7 | COPZ1 | 0,040464921 | 2 | 0,079665321 | 0,040701286 | Coatomer subunit zeta-1 |
| P23229 | ITGA6 | 0,041046736 | 14 | 0,015764509 | 0,010099138 | Integrin alpha-6 |
| O00299 | CLIC1 | 0,041064389 | 12 | 0,054790703 | 0,030085966 | Chloride intracellular channel protein 1 |
| P53621 | COPA | 0,041440772 | 15 | 0,03034387 | 0,018030416 | Coatomer subunit alpha |
| P63220 | RPS21 | 0,045461959 | 4 | 0,059611543 | 0,032229538 | 40S ribosomal protein S21 |
| P22626 | HNRNPA2B1 | 0,046387005 | 15 | 0,006488986 | 0,004626929 | Heterogeneous nuclear ribonucleoproteins A2/B1 |
| Q15084 | PDIA6 | 0,046553809 | 14 | 0,01159917 | 0,007691094 | Protein disulfide-isomerase A6 |
| P13797 | PLS3 | 0,046580226 | 28 | 4,59E-09 | 9,42E-09 | Plastin-3 |
| H3BNC9 | NaN | 0,046770854 | 8 | 0,015211463 | 0,009757549 | Uncharacterized protein |
| P39019 | RPS19 | 0,046902371 | 7 | 0,067274116 | 0,035457643 | 40S ribosomal protein S19 |
| P02042 | HBD | 0,047842485 | 6 | 8,52E-09 | 1,67E-08 | Hemoglobin subunit delta |

| | | | | | | |
|---|---|---|---|---|---|---|
| Q9UQ80 | PA2G4 | 0,048179583 | 9 | 0,077447724 | 0,039650656 | Proliferation-associated protein 2G4 |
| Q9Y265 | RUVBL1 | 0,048282784 | 9 | 0,000193171 | 0,000192779 | RuvB-like 1 |
| F5H7S7 | IQGAP2 | 0,048712379 | 17 | 0,001322639 | 0,001129754 | Ras GTPase-activating-like protein IQGAP2 |
| C9JIG9 | OXSR1 | 0,049019269 | 3 | 0,052998451 | 0,029330976 | Serine/threonine-protein kinase OSR1 |
| P32119 | PRDX2 | 0,049052913 | 14 | 0,00040578 | 0,000387658 | Peroxiredoxin-2 |
| P49368 | CCT3 | 0,049631643 | 14 | 1,77E-05 | 2,10E-05 | T-complex protein 1 subunit gamma |
| F8VVM2 | SLC25A3 | 0,049885489 | 7 | 0,001352713 | 0,001151445 | Phosphate carrier protein, mitochondrial |
| P35606 | COPB2 | 0,050450235 | 13 | 0,058736613 | 0,031896704 | Coatomer subunit beta' |
| P16989 | YBX3 | 0,050863607 | 8 | 0,058994361 | 0,031966107 | Y-box-binding protein 3 |
| P43304 | GPD2 | 0,052131548 | 11 | 0,001562037 | 0,001293808 | Glycerol-3-phosphate dehydrogenase, mitochondrial |
| P51148 | RAB5C | 0,05216619 | 9 | 0,004574192 | 0,003369016 | Ras-related protein Rab-5C |
| Q15046 | KARS | 0,052202364 | 7 | 0,085906865 | 0,043261185 | Lysine--tRNA ligase |
| B5ME19 | EIF3CL | 0,053042264 | 8 | 0,046013464 | 0,025949552 | Eukaryotic translation initiation factor 3 subunit C-like protein |
| P55884 | EIF3B | 0,053387978 | 15 | 0,059268299 | 0,032079211 | Eukaryotic translation initiation factor 3 subunit B |
| Q99832 | CCT7 | 0,054084077 | 20 | 0,013496272 | 0,008748571 | T-complex protein 1 subunit eta |
| P61247 | RPS3A | 0,054231603 | 11 | 0,006781374 | 0,004786853 | 40S ribosomal protein S3a |
| P09874 | PARP1 | 0,054849313 | 4 | 0,067849381 | 0,035588375 | Poly [ADP-ribose] polymerase 1 |
| P49755 | TMED10 | 0,055211503 | 6 | 0,009693873 | 0,006556977 | Transmembrane emp24 domain-containing protein 10 |
| P23396 | RPS3 | 0,055752483 | 14 | 0,066650732 | 0,035260387 | 40S ribosomal protein S3 |
| P63173 | RPL38 | 0,055885638 | 3 | 0,056731418 | 0,030978754 | 60S ribosomal protein L38 |
| D6RFH4 | CYB5B | 0,056212287 | 2 | 0,051006102 | 0,028484679 | Cytochrome b5 type B |
| P63244 | RACK1 | 0,056352303 | 16 | 2,01E-05 | 2,35E-05 | Receptor of activated protein C kinase 1 |
| Q92900 | UPF1 | 0,056353529 | 2 | 0,095718988 | 0,047235448 | Regulator of nonsense transcripts 1 |
| A0A087WUQ6 | GPX1 | 0,057516363 | 5 | 0,06139115 | 0,03311891 | Glutathione peroxidase |
| P35749-4 | MYH11 | 0,057585166 | 100 | 1,87E-05 | 2,21E-05 | Isoform 4 of Myosin-11 |
| P62701 | RPS4X | 0,057694806 | 12 | 0,001760467 | 0,001438787 | 40S ribosomal protein S4, X isoform |
| Q9BS26 | ERP44 | 0,057816846 | 4 | 0,073905494 | 0,038235019 | Endoplasmic reticulum resident protein 44 |
| P18206 | VCL | 0,061670591 | 33 | 0,008270518 | 0,005691042 | Vinculin |
| E9PEB5 | FUBP1 | 0,062514167 | 5 | 0,021409639 | 0,013316741 | Far upstream element-binding protein 1 |
| H7C3G9 | NAGK | 0,062758579 | 11 | 0,060920169 | 0,032900904 | N-acetyl-D-glucosamine kinase |
| P04075 | ALDOA | 0,062825899 | 27 | 0,014649294 | 0,009433838 | Fructose-bisphosphate aldolase A |
| P18124 | RPL7 | 0,063237168 | 11 | 0,025520553 | 0,015578303 | 60S ribosomal protein L7 |
| E9PAV3 | NACA | 0,06443394 | 4 | 0,009646752 | 0,006537468 | Nascent polypeptide-associated complex subunit alpha, muscle-specific form |
| P20073 | ANXA7 | 0,065382694 | 8 | 1,16E-05 | 1,43E-05 | Annexin A7 |
| Q9Y277 | VDAC3 | 0,065464184 | 7 | 0,0006768 | 0,000613233 | Voltage-dependent anion-selective channel protein 3 |
| P55268 | LAMB2 | 0,065501218 | 29 | 1,61E-05 | 1,94E-05 | Laminin subunit beta-2 |
| P62280 | RPS11 | 0,065805629 | 5 | 0,052822707 | 0,029266635 | 40S ribosomal protein S11 |
| F8VQE1 | LIMA1 | 0,066444506 | 6 | 0,007327437 | 0,005113616 | LIM domain and actin-binding protein 1 |

| | | | | | | |
|---|---|---|---|---|---|---|
| Q16543 | CDC37 | 0,066852409 | 4 | 0,029704741 | 0,017714828 | Hsp90 co-chaperone Cdc37 |
| Q06323 | PSME1 | 0,067935834 | 9 | 0,053182434 | 0,029359379 | Proteasome activator complex subunit 1 |
| Q9NT62 | ATG3 | 0,068062636 | 2 | 0,097706879 | 0,047880263 | Ubiquitin-like-conjugating enzyme ATG3 |
| P23381 | WARS | 0,068921679 | 6 | 0,097257239 | 0,047802759 | Tryptophan--tRNA ligase, cytoplasmic |
| P31946 | YWHAB | 0,06908459 | 9 | 0,055615868 | 0,030505024 | 14-3-3 protein beta/alpha |
| A0A2R8Y6J3 | RPL5 | 0,069274411 | 9 | 8,72E-05 | 9,30E-05 | 60S ribosomal protein L5 (Fragment) |
| K7ELC7 | RPL27 | 0,069755743 | 4 | 0,019536948 | 0,012323305 | 60S ribosomal protein L27 (Fragment) |
| P39023 | RPL3 | 0,070124026 | 11 | 0,000269156 | 0,000264849 | 60S ribosomal protein L3 |
| P02730 | SLC4A1 | 0,071108251 | 24 | 0,005771269 | 0,004163438 | Band 3 anion transport protein |
| P30050 | RPL12 | 0,071750051 | 6 | 2,86E-05 | 3,26E-05 | 60S ribosomal protein L12 |
| Q7KZF4 | SND1 | 0,071794565 | 16 | 0,006202003 | 0,004441609 | Staphylococcal nuclease domain-containing protein 1 |
| F8VPD4 | CAD | 0,07309158 | 4 | 0,0060681 | 0,004358402 | CAD protein |
| P50914 | RPL14 | 0,074359561 | 4 | 0,012755351 | 0,008356368 | 60S ribosomal protein L14 |
| Q9HB71 | CACYBP | 0,074558026 | 3 | 0,028088394 | 0,016853037 | Calcyclin-binding protein |
| P04406 | GAPDH | 0,074605691 | 18 | 0,003227819 | 0,002465974 | Glyceraldehyde-3-phosphate dehydrogenase |
| P62851 | RPS25 | 0,075153336 | 4 | 0,001893162 | 0,001537022 | 40S ribosomal protein S25 |
| C9JNW5 | RPL24 | 0,075246221 | 3 | 0,063160099 | 0,033887425 | 60S ribosomal protein L24 |
| P62942 | FKBP1A | 0,075886155 | 3 | 0,063682061 | 0,034015031 | Peptidyl-prolyl cis-trans isomerase FKBP1A |
| B0YIW6 | ARCN1 | 0,075977554 | 6 | 0,007830316 | 0,005404613 | Archain 1, isoform CRA_a |
| Q13347 | EIF3I | 0,0760406 | 3 | 0,03855595 | 0,022212561 | Eukaryotic translation initiation factor 3 subunit I |
| P62136 | PPP1CA | 0,076464976 | 9 | 0,092950795 | 0,046007838 | Serine/threonine-protein phosphatase PP1-alpha catalytic subunit |
| P62979 | RPS27A | 0,076644723 | 5 | 0,004513904 | 0,003329596 | Ubiquitin-40S ribosomal protein S27a |
| J3KTL2 | SRSF1 | 0,076963675 | 5 | 0,053288466 | 0,029359379 | Serine/arginine-rich-splicing factor 1 |
| A0A075B7D9 | TAF15 | 0,07705801 | 4 | 0,004195457 | 0,003127523 | TATA-binding protein-associated factor 2N |
| A0A3B3ISU0 | DSC2 | 0,07729933 | 2 | 0,086234174 | 0,043381609 | Desmocollin-2 |
| P21281 | ATP6V1B2 | 0,077466325 | 12 | 0,000338368 | 0,000326426 | V-type proton ATPase subunit B, brain isoform |
| Q14152 | EIF3A | 0,077760497 | 18 | 0,00179536 | 0,001462445 | Eukaryotic translation initiation factor 3 subunit A |
| P22087 | FBL | 0,077789365 | 7 | 0,044698753 | 0,025307004 | rRNA 2'-O-methyltransferase fibrillarin |
| Q00839 | HNRNPU | 0,078815161 | 21 | 0,000294129 | 0,000287697 | Heterogeneous nuclear ribonucleoprotein U |
| Q15067-2 | ACOX1 | 0,078815378 | 8 | 0,076656022 | 0,039409365 | Isoform 2 of Peroxisomal acyl-coenzyme A oxidase 1 |
| P62249 | RPS16 | 0,078959332 | 6 | 0,000419224 | 0,000398952 | 40S ribosomal protein S16 |
| P12814-2 | ACTN1 | 0,079332503 | 43 | 2,15E-35 | 2,46E-34 | Isoform 2 of Alpha-actinin-1 |
| Q02878 | RPL6 | 0,080363991 | 8 | 0,063598737 | 0,034015031 | 60S ribosomal protein L6 |
| Q13838 | DDX39B | 0,080916862 | 12 | 6,07E-06 | 7,96E-06 | Spliceosome RNA helicase DDX39B |
| Q09028 | RBBP4 | 0,080970244 | 6 | 0,039741426 | 0,022815381 | Histone-binding protein RBBP4 |
| P21333 | FLNA | 0,081653097 | 102 | 4,24E-21 | 2,27E-20 | Filamin-A |
| P56537 | EIF6 | 0,081708725 | 5 | 0,002310599 | 0,001845478 | Eukaryotic translation initiation factor 6 |
| P61026 | RAB10 | 0,081875962 | 5 | 0,040164987 | 0,023004859 | Ras-related protein Rab-10 |

| | | | | | | |
|---|---|---|---|---|---|---|
| P62277 | RPS13 | 0,081914136 | 8 | 0,001158003 | 0,001001296 | 40S ribosomal protein S13 |
| M0R117 | RPL18A | 0,08204511 | 4 | 0,048296999 | 0,027094782 | 60S ribosomal protein L18a |
| E9PPG9 | RAE1 | 0,082463717 | 2 | 0,032854819 | 0,019338845 | mRNA export factor |
| P62906 | RPL10A | 0,082616907 | 9 | 0,007770147 | 0,005384383 | 60S ribosomal protein L10a |
| H0Y8G5 | HNRNPD | 0,083658151 | 9 | 4,59E-06 | 6,22E-06 | Heterogeneous nuclear ribonucleoprotein D0 (Fragment) |
| P49458 | SRP9 | 0,084569639 | 2 | 0,021798037 | 0,013541205 | Signal recognition particle 9 kDa protein |
| Q53GQ0 | HSD17B12 | 0,084733002 | 6 | 0,011375735 | 0,007563327 | Very-long-chain 3-oxoacyl-CoA reductase |
| P04843 | RPN1 | 0,08516553 | 14 | 6,28E-09 | 1,26E-08 | Dolichyl-diphosphooligosaccharide--protein glycosyltransferase subunit 1 |
| J3QRP6 | SLC9A3R1 | 0,085180764 | 3 | 0,020674564 | 0,01294133 | Na(+)/H(+) exchange regulatory cofactor NHE-RF1 (Fragment) |
| A0A0C4DGS1 | DDOST | 0,085287024 | 11 | 0,002840757 | 0,002211475 | Dolichyl-diphosphooligosaccharide--protein glycosyltransferase 48 kDa subunit |
| P07305 | H1F0 | 0,085670826 | 4 | 0,026112717 | 0,015861058 | Histone H1.0 |
| P11171 | EPB41 | 0,08633796 | 12 | 0,078748277 | 0,040274587 | Protein 4.1 |
| Q04917 | YWHAH | 0,087224097 | 5 | 0,004086383 | 0,003064787 | 14-3-3 protein eta |
| A0A0A0MRA5 | HNRNPUL1 | 0,087377882 | 6 | 0,019579911 | 0,012329916 | Heterogeneous nuclear ribonucleoprotein U-like protein 1 |
| H0Y8C6 | IPO5 | 0,087980769 | 10 | 0,013041176 | 0,008487115 | Importin-5 (Fragment) |
| A0A2R8Y811 | RPS14 | 0,088420517 | 8 | 0,072277258 | 0,037630065 | 40S ribosomal protein S14 (Fragment) |
| A0A2U3TZL5 | CD59 | 0,088818317 | 2 | 0,00039845 | 0,000382476 | CD59 glycoprotein (Fragment) |
| F8W1A4 | AK2 | 0,089019828 | 9 | 0,049169947 | 0,027553091 | Adenylate kinase 2, mitochondrial |
| P07900 | HSP90AA1 | 0,089700003 | 17 | 5,06E-07 | 7,84E-07 | Heat shock protein HSP 90-alpha |
| P00747 | PLG | 0,089825754 | 29 | 1,96E-10 | 4,62E-10 | Plasminogen |
| P60953 | CDC42 | 0,090002918 | 5 | 0,092095623 | 0,045768734 | Cell division control protein 42 homolog |
| F6SDV2 | TINAGL1 | 0,090381925 | 7 | 0,010734287 | 0,007165901 | Tubulointerstitial nephritis antigen-like 1, isoform CRA_a |
| Q5T123 | SH3BGRL3 | 0,090952392 | 3 | 0,012944864 | 0,008469246 | SH3 domain-binding glutamic acid-rich-like protein 3 |
| P78527 | PRKDC | 0,091283332 | 32 | 1,96E-09 | 4,20E-09 | DNA-dependent protein kinase catalytic subunit |
| Q9NYU2 | UGGT1 | 0,091383115 | 6 | 0,089972323 | 0,044986161 | UDP-glucose:glycoprotein glucosyltransferase 1 |
| P60866 | RPS20 | 0,091671185 | 3 | 0,008499561 | 0,005832335 | 40S ribosomal protein S20 |
| P08236 | GUSB | 0,092610527 | 3 | 0,006534154 | 0,00464567 | Beta-glucuronidase |
| P62316 | SNRPD2 | 0,09328834 | 3 | 0,000849702 | 0,000757343 | Small nuclear ribonucleoprotein Sm D2 |
| P02747 | C1QC | 0,094058368 | 3 | 0,080254483 | 0,040959757 | Complement C1q subcomponent subunit C |
| B7Z7P8 | ETF1 | 0,095259855 | 6 | 0,013778945 | 0,008920054 | Eukaryotic peptide chain release factor subunit 1 |
| Q96BM9 | ARL8A | 0,095728365 | 6 | 0,099714203 | 0,048670028 | ADP-ribosylation factor-like protein 8A |
| P19827 | ITIH1 | 0,097600005 | 25 | 0,000192053 | 0,000192053 | Inter-alpha-trypsin inhibitor heavy chain H1 |
| A0A0A0MSM0 | HSPH1 | 0,097632704 | 5 | 0,063652603 | 0,034015031 | Heat shock protein 105 kDa |
| A0A087WZP6 | NID2 | 0,098417826 | 5 | 0,008469606 | 0,005819897 | Nidogen-2 |
| H0Y2P0 | CD44 | 0,098716675 | 4 | 0,000659122 | 0,000601648 | CD44 antigen (Fragment) |
| P62266 | RPS23 | 0,098771482 | 4 | 0,032860314 | 0,019338845 | 40S ribosomal protein S23 |
| P24844 | MYL9 | 0,098891783 | 9 | 0,004981875 | 0,003636621 | Myosin regulatory light polypeptide 9 |
| Q06210 | GFPT1 | 0,098918013 | 4 | 0,016145559 | 0,010329798 | Glutamine--fructose-6-phosphate aminotransferase [isomerizing] 1 |

| | | | | | | |
|---|---|---|---|---|---|---|
| M0R0F0 | RPS5 | 0,099493413 | 5 | 0,01027699 | 0,006898062 | 40S ribosomal protein S5 (Fragment) |
| P15880 | RPS2 | 0,099613187 | 10 | 0,004244869 | 0,003154797 | 40S ribosomal protein S2 |
| Q9BSJ8 | ESYT1 | 0,099701582 | 12 | 0,027799042 | 0,016720206 | Extended synaptotagmin-1 |
| P53396 | ACLY | 0,101233761 | 15 | 0,001577772 | 0,001302456 | ATP-citrate synthase |
| A0A0D9SEM4 | SRSF4 | 0,101393925 | 2 | 0,08328034 | 0,042284755 | Serine/arginine-rich-splicing factor 4 (Fragment) |
| A0A087X271 | CNN2 | 0,102195563 | 2 | 0,058635511 | 0,031877098 | Calponin (Fragment) |
| P62753 | RPS6 | 0,103035716 | 5 | 0,009415264 | 0,006389393 | 40S ribosomal protein S6 |
| P11277 | SPTB | 0,103401491 | 53 | 0,00074106 | 0,000666547 | Spectrin beta chain, erythrocytic |
| P26641 | EEF1G | 0,103548227 | 13 | 0,00386564 | 0,002917017 | Elongation factor 1-gamma |
| H0Y2Y8 | ZYX | 0,105195261 | 7 | 0,066208367 | 0,035064065 | Zyxin (Fragment) |
| P10644 | PRKAR1A | 0,10574802 | 6 | 0,026800679 | 0,016198936 | cAMP-dependent protein kinase type I-alpha regulatory subunit |
| P42224 | STAT1 | 0,107019297 | 9 | 0,00898601 | 0,00613885 | Signal transducer and activator of transcription 1-alpha/beta |
| A0A0A0MQS9 | LAMA4 | 0,107851309 | 8 | 0,009715521 | 0,006556977 | Laminin subunit alpha-4 |
| A0A087WY82 | F11R | 0,108047302 | 2 | 0,009964144 | 0,006697212 | Junctional adhesion molecule A |
| Q9BVK6 | TMED9 | 0,108128759 | 3 | 0,009016817 | 0,006144424 | Transmembrane emp24 domain-containing protein 9 |
| Q2TAY7 | SMU1 | 0,108479478 | 3 | 0,06244052 | 0,033593645 | WD40 repeat-containing protein SMU1 |
| G8JLB6 | HNRNPH1 | 0,108761994 | 10 | 0,000636709 | 0,000584442 | Heterogeneous nuclear ribonucleoprotein H |
| E9PJD9 | RPL27A | 0,108930092 | 2 | 0,066830299 | 0,035317408 | 60S ribosomal protein L27a |
| Q15233 | NONO | 0,108931703 | 8 | 0,03534169 | 0,020577647 | Non-POU domain-containing octamer-binding protein |
| Q9UBS4 | DNAJB11 | 0,109028132 | 2 | 0,034798918 | 0,020357988 | DnaJ homolog subfamily B member 11 |
| A0A0B4J2C3 | TPT1 | 0,109564985 | 6 | 0,003965874 | 0,002978947 | Translationally-controlled tumor protein |
| P30040 | ERP29 | 0,109665163 | 7 | 0,000110595 | 0,000115331 | Endoplasmic reticulum resident protein 29 |
| P07237 | P4HB | 0,109893957 | 20 | 0,000998416 | 0,00087718 | Protein disulfide-isomerase |
| P62241 | RPS8 | 0,110398143 | 8 | 0,002994052 | 0,002319801 | 40S ribosomal protein S8 |
| P19338 | NCL | 0,110480292 | 15 | 4,64E-06 | 6,26E-06 | Nucleolin |
| E5RHG8 | ELOC | 0,112474237 | 2 | 0,000395614 | 0,000380905 | Elongin-C (Fragment) |
| O43747 | AP1G1 | 0,11249371 | 7 | 0,036065331 | 0,020949401 | AP-1 complex subunit gamma-1 |
| P16144 | ITGB4 | 0,112589305 | 20 | 4,13E-06 | 5,67E-06 | Integrin beta-4 |
| P60842 | EIF4A1 | 0,115218204 | 22 | 1,20E-22 | 7,20E-22 | Eukaryotic initiation factor 4A-I |
| Q9BXP5 | SRRT | 0,116306504 | 2 | 0,022367143 | 0,013829019 | Serrate RNA effector molecule homolog |
| P62424 | RPL7A | 0,116908366 | 9 | 4,64E-06 | 6,26E-06 | 60S ribosomal protein L7a |
| Q9Y230 | RUVBL2 | 0,11734587 | 8 | 0,000132838 | 0,000137049 | RuvB-like 2 |
| A0A087WUC6 | SPCS2 | 0,117425629 | 2 | 0,000457866 | 0,000429743 | Signal peptidase complex subunit 2 |
| P36578 | RPL4 | 0,118291399 | 16 | 1,04E-05 | 1,31E-05 | 60S ribosomal protein L4 |
| E5RI99 | RPL30 | 0,120058654 | 5 | 0,000117072 | 0,000121775 | 60S ribosomal protein L30 (Fragment) |
| B2R5W2 | HNRNPC | 0,120290157 | 10 | 5,71E-06 | 7,57E-06 | Heterogeneous nuclear ribonucleoproteins C1/C2 |
| P24534 | EEF1B2 | 0,121803679 | 4 | 1,06E-05 | 1,33E-05 | Elongation factor 1-beta |
| O14929 | HAT1 | 0,122429922 | 2 | 0,001516381 | 0,001260236 | Histone acetyltransferase type B catalytic subunit |

| | | | | | | |
|---|---|---|---|---|---|---|
| P56192 | MARS | 0,1233866<br>19 | 4 | 0,011955168 | 0,0079164<br>77 | Methionine--tRNA ligase, cytoplasmic |
| P16930 | FAH | 0,1239591<br>1 | 3 | 0,037779268 | 0,0217906<br>21 | Fumarylacetoacetase |
| P04844 | RPN2 | 0,1250649<br>31 | 14 | 0,004947158 | 0,0036166<br>44 | Dolichyl-diphosphooligosaccharide--protein glycosyltransferase subunit 2 |
| P04179 | SOD2 | 0,1262628<br>95 | 7 | 1,07E-05 | 1,33E-05 | Superoxide dismutase [Mn], mitochondrial |
| P62304 | SNRPE | 0,1269889<br>1 | 3 | 0,088798548 | 0,0444444<br>41 | Small nuclear ribonucleoprotein E |
| P31153 | MAT2A | 0,1274781<br>15 | 5 | 0,003126046 | 0,0024031<br>48 | S-adenosylmethionine synthase isoform type-2 |
| A0A024R4<br>M0 | RPS9 | 0,1279767<br>7 | 8 | 0,000603109 | 0,0005598<br>67 | 40S ribosomal protein S9 |
| P62318 | SNRPD3 | 0,1280556<br>71 | 3 | 0,00044506 | 0,0004188<br>19 | Small nuclear ribonucleoprotein Sm D3 |
| J3KND3 | MYL6 | 0,1285647<br>68 | 8 | 0,012383374 | 0,0081670<br>51 | Myosin light polypeptide 6 |
| P00915 | CA1 | 0,1288919<br>65 | 14 | 2,35E-15 | 8,46E-15 | Carbonic anhydrase 1 |
| P05198 | EIF2S1 | 0,1291320<br>14 | 6 | 4,88E-07 | 7,61E-07 | Eukaryotic translation initiation factor 2 subunit 1 |
| A0A0C4DG<br>Z5 | YARS | 0,1292586<br>03 | 3 | 0,037269947 | 0,0215727<br>22 | Tyrosine--tRNA ligase |
| Q9Y4G6 | TLN2 | 0,1294595<br>48 | 7 | 0,005157346 | 0,0037496<br>78 | Talin-2 |
| P00390 | GSR | 0,1302243<br>41 | 7 | 0,00308524 | 0,0023754<br>9 | Glutathione reductase, mitochondrial |
| Q9NS69 | TOMM22 | 0,1303563<br>46 | 2 | 0,014841133 | 0,0095324<br>25 | Mitochondrial import receptor subunit TOM22 homolog |
| P11047 | LAMC1 | 0,1309492<br>54 | 22 | 6,95E-06 | 8,98E-06 | Laminin subunit gamma-1 |
| A0A087X0K<br>0 | COL15A1 | 0,1313656<br>65 | 8 | 8,69E-06 | 1,11E-05 | Collagen alpha-1(XV) chain |
| E9PIX6 | PRMT1 | 0,1332672<br>45 | 4 | 0,026721863 | 0,0161711<br>65 | Protein arginine N-methyltransferase 1 (Fragment) |
| P62491 | RAB11A | 0,1352190<br>57 | 8 | 0,000955749 | 0,0008435<br>95 | Ras-related protein Rab-11A |
| P14866 | HNRNPL | 0,1355382<br>2 | 13 | 6,23E-08 | 1,08E-07 | Heterogeneous nuclear ribonucleoprotein L |
| P12004 | PCNA | 0,1358144<br>02 | 3 | 0,035557484 | 0,0206788<br>2 | Proliferating cell nuclear antigen |
| A0A3B3IUB<br>5 | HM13 | 0,1366801<br>31 | 2 | 0,000955726 | 0,0008435<br>95 | Minor histocompatibility antigen H13 |
| P11142 | HSPA8 | 0,1377279<br>21 | 31 | 8,05E-21 | 4,21E-20 | Heat shock cognate 71 kDa protein |
| Q16853 | AOC3 | 0,1393864<br>62 | 13 | 0,012001884 | 0,0079367<br>3 | Membrane primary amine oxidase |
| E9PRY8 | EEF1D | 0,1404270<br>64 | 12 | 0,000240417 | 0,0002375<br>21 | Elongation factor 1-delta |
| E9PK25 | CFL1 | 0,1414151<br>63 | 15 | 3,75E-08 | 6,74E-08 | Cofilin-1 |
| O95573 | ACSL3 | 0,1420276<br>59 | 2 | 0,03718916 | 0,0215513<br>16 | Long-chain-fatty-acid--CoA ligase 3 |
| J3QLE5 | SNRPN | 0,1422048<br>56 | 4 | 0,07358867 | 0,0381111<br>85 | Small nuclear ribonucleoprotein-associated protein N (Fragment) |
| P11021 | HSPA5 | 0,1431948<br>58 | 25 | 9,99E-37 | 1,20E-35 | Endoplasmic reticulum chaperone BiP |
| Q6IBS0 | TWF2 | 0,1438988<br>7 | 4 | 0,000160306 | 0,0001629<br>55 | Twinfilin-2 |
| P08238 | HSP90AB1 | 0,1444238<br>05 | 33 | 1,90E-23 | 1,17E-22 | Heat shock protein HSP 90-beta |
| O75533 | SF3B1 | 0,1445196<br>17 | 10 | 0,003276932 | 0,0024996<br>13 | Splicing factor 3B subunit 1 |
| Q9Y2V2 | CARHSP1 | 0,1447240<br>41 | 2 | 0,01035442 | 0,0069405<br>65 | Calcium-regulated heat-stable protein 1 |
| P49756 | RBM25 | 0,1449374<br>83 | 3 | 0,046044632 | 0,0259495<br>52 | RNA-binding protein 25 |
| P52597 | HNRNPF | 0,1452652<br>96 | 6 | 2,52E-06 | 3,56E-06 | Heterogeneous nuclear ribonucleoprotein F |
| P63000 | RAC1 | 0,1460969<br>7 | 4 | 0,031613385 | 0,0186944<br>54 | Ras-related C3 botulinum toxin substrate 1 |
| P23246 | SFPQ | 0,1471577<br>61 | 13 | 6,58E-15 | 2,28E-14 | Splicing factor, proline- and glutamine-rich |
| P13473 | LAMP2 | 0,1472732 | 4 | 0,027343288 | 0,0165066<br>23 | Lysosome-associated membrane glycoprotein 2 |

| | | | | | | |
|---|---|---|---|---|---|---|
| A0A087X054 | HYOU1 | 0,147396784 | 9 | 0,000198216 | 0,000197413 | Hypoxia up-regulated protein 1 |
| J3KP15 | SRSF2 | 0,14762078 | 2 | 0,000989658 | 0,000871041 | Serine/arginine-rich-splicing factor 2 (Fragment) |
| Q9Y6C2 | EMILIN1 | 0,148480978 | 7 | 0,093761489 | 0,046362465 | EMILIN-1 |
| O75367 | H2AFY | 0,148919304 | 10 | 3,37E-15 | 1,18E-14 | Core histone macro-H2A.1 |
| A0A0D9SF53 | DDX3X | 0,149081943 | 16 | 0,022873098 | 0,014084561 | ATP-dependent RNA helicase DDX3X |
| Q13283 | G3BP1 | 0,151413126 | 6 | 0,000163688 | 0,000165394 | Ras GTPase-activating protein-binding protein 1 |
| A0A2R8Y5S7 | RDX | 0,152388948 | 12 | 3,17E-07 | 5,06E-07 | Radixin |
| P23284 | PPIB | 0,152733656 | 8 | 4,54E-13 | 1,42E-12 | Peptidyl-prolyl cis-trans isomerase B |
| P11166 | SLC2A1 | 0,152914264 | 3 | 0,080666531 | 0,041084817 | Solute carrier family 2, facilitated glucose transporter member 1 |
| P61769 | B2M | 0,153458366 | 2 | 0,001616603 | 0,001332276 | Beta-2-microglobulin |
| Q15393 | SF3B3 | 0,153729951 | 8 | 0,062493811 | 0,033593645 | Splicing factor 3B subunit 3 |
| O75534 | CSDE1 | 0,153967181 | 4 | 0,02976775 | 0,017730912 | Cold shock domain-containing protein E1 |
| P08697 | SERPINF2 | 0,15445362 | 9 | 0,002119691 | 0,001706854 | Alpha-2-antiplasmin |
| A0A087X2D0 | SRSF3 | 0,156958816 | 4 | 0,000132871 | 0,000137049 | Serine/arginine-rich-splicing factor 3 |
| P06748 | NPM1 | 0,157633787 | 8 | 1,43E-08 | 2,68E-08 | Nucleophosmin |
| F8W6I7 | HNRNPA1 | 0,15786451 | 18 | 3,39E-11 | 8,87E-11 | Heterogeneous nuclear ribonucleoprotein A1 |
| P16403 | HIST1H1C | 0,160634495 | 10 | 0,001421775 | 0,001201913 | Histone H1.2 |
| P62879 | GNB2 | 0,167555442 | 8 | 5,79E-09 | 1,16E-08 | Guanine nucleotide-binding protein G(I)/G(S)/G(T) subunit beta-2 |
| P40763 | STAT3 | 0,168738354 | 16 | 9,65E-05 | 0,000102147 | Signal transducer and activator of transcription 3 |
| P27797 | CALR | 0,169510021 | 14 | 5,09E-19 | 2,41E-18 | Calreticulin |
| I3L0N3 | NSF | 0,170256339 | 8 | 2,40E-05 | 2,76E-05 | Vesicle-fusing ATPase |
| Q9UH99 | SUN2 | 0,171005947 | 6 | 0,003748683 | 0,002833106 | SUN domain-containing protein 2 |
| P55265 | ADAR | 0,171613007 | 3 | 0,003920816 | 0,002949605 | Double-stranded RNA-specific adenosine deaminase |
| Q9Y490 | TLN1 | 0,172846404 | 62 | 3,87E-22 | 2,21E-21 | Talin-1 |
| P26373 | RPL13 | 0,174425289 | 4 | 6,82E-05 | 7,44E-05 | 60S ribosomal protein L13 |
| J3KTE4 | RPL19 | 0,174958368 | 3 | 0,001475405 | 0,001234523 | Ribosomal protein L19 |
| P14543 | NID1 | 0,175641705 | 11 | 2,16E-05 | 2,52E-05 | Nidogen-1 |
| P16452 | EPB42 | 0,176790933 | 8 | 0,056067372 | 0,030684257 | Erythrocyte membrane protein band 4.2 |
| P53999 | SUB1 | 0,177409551 | 5 | 5,79E-09 | 1,16E-08 | Activated RNA polymerase II transcriptional coactivator p15 |
| P09661 | SNRPA1 | 0,179781386 | 2 | 0,001115888 | 0,000971712 | U2 small nuclear ribonucleoprotein A' |
| O15260 | SURF4 | 0,180665829 | 3 | 0,091865129 | 0,045700347 | Surfeit locus protein 4 |
| Q08211 | DHX9 | 0,182459695 | 24 | 9,85E-14 | 3,19E-13 | ATP-dependent RNA helicase A |
| A0A087WSW9 | TXNRD1 | 0,185370668 | 4 | 0,008202933 | 0,005652441 | Thioredoxin reductase 1, cytoplasmic |
| P11940 | PABPC1 | 0,190185573 | 13 | 2,60E-09 | 5,51E-09 | Polyadenylate-binding protein 1 |
| P00492 | HPRT1 | 0,190548374 | 6 | 0,001442039 | 0,001212792 | Hypoxanthine-guanine phosphoribosyltransferase |
| P32926 | DSG3 | 0,191457421 | 15 | 1,41E-11 | 3,85E-11 | Desmoglein-3 |
| P02792 | FTL | 0,191485924 | 4 | 0,000103801 | 0,000109124 | Ferritin light chain |
| P38919 | EIF4A3 | 0,192839623 | 7 | 3,89E-06 | 5,37E-06 | Eukaryotic initiation factor 4A-III |

| | | | | | | |
|---|---|---|---|---|---|---|
| B0QZ18 | CPNE1 | 0,192856279 | 4 | 0,090560591 | 0,045182902 | Copine-1 |
| A0A0S2Z4L3 | PROS1 | 0,193530497 | 5 | 1,69E-05 | 2,01E-05 | Protein S isoform 2 (Fragment) |
| B1AK87 | CAPZB | 0,193538649 | 8 | 1,21E-17 | 5,22E-17 | Capping protein (Actin filament) muscle Z-line, beta, isoform CRA_a |
| P61586 | RHOA | 0,200228829 | 2 | 0,003181635 | 0,002438262 | Transforming protein RhoA |
| Q7Z406-6 | MYH14 | 0,200706045 | 43 | 0,001493481 | 0,001245411 | Isoform 6 of Myosin-14 |
| P69905 | HBA1 | 0,202572047 | 13 | 3,69E-16 | 1,45E-15 | Hemoglobin subunit alpha |
| C9J6B1 | RALB | 0,202744352 | 3 | 1,27E-05 | 1,55E-05 | Ras-related protein Ral-B (Fragment) |
| Q9Y6A9 | SPCS1 | 0,202787517 | 2 | 5,49E-06 | 7,32E-06 | Signal peptidase complex subunit 1 |
| Q9NP72 | RAB18 | 0,205801219 | 2 | 0,001023491 | 0,000897607 | Ras-related protein Rab-18 |
| Q7L1Q6 | BZW1 | 0,205826297 | 6 | 0,000264148 | 0,000260442 | Basic leucine zipper and W2 domain-containing protein 1 |
| P08134 | RHOC | 0,20755882 | 9 | 4,60E-11 | 1,17E-10 | Rho-related GTP-binding protein RhoC |
| A0A087WYS1 | UGP2 | 0,209202441 | 10 | 5,51E-05 | 6,08E-05 | UTP--glucose-1-phosphate uridylyltransferase |
| P30460 | HLA-B | 0,209740487 | 2 | 0,000781085 | 0,000699989 | HLA class I histocompatibility antigen, B-8 alpha chain |
| Q9Y6N5 | SQOR | 0,210387411 | 9 | 4,09E-16 | 1,60E-15 | Sulfide:quinone oxidoreductase, mitochondrial |
| Q9BTT0 | ANP32E | 0,214613092 | 2 | 0,007270193 | 0,005095349 | Acidic leucine-rich nuclear phosphoprotein 32 family member E |
| P0C0S5 | H2AFZ | 0,222930118 | 3 | 2,22E-05 | 2,56E-05 | Histone H2A.Z |
| H0Y449 | YBX1 | 0,224791493 | 5 | 0,000103568 | 0,000109112 | Nuclease-sensitive element-binding protein 1 (Fragment) |
| B1AKG0 | CFHR1 | 0,225582784 | 4 | 0,002743003 | 0,00214556 | Complement factor H-related protein 1 |
| B1AP13 | CD55 | 0,229616438 | 4 | 6,02E-05 | 6,58E-05 | Complement decay-accelerating factor |
| O75083 | WDR1 | 0,231308534 | 17 | 2,02E-18 | 9,20E-18 | WD repeat-containing protein 1 |
| P10319 | HLA-B | 0,231333108 | 8 | 0,000617888 | 0,000572507 | HLA class I histocompatibility antigen, B-58 alpha chain |
| P39748 | FEN1 | 0,23267397 | 2 | 4,70E-09 | 9,55E-09 | Flap endonuclease 1 |
| A0A087WWU8 | TPM3 | 0,232854334 | 11 | 6,94E-22 | 3,88E-21 | Tropomyosin alpha-3 chain |
| P07738 | BPGM | 0,23312909 | 5 | 0,019597549 | 0,012329916 | Bisphosphoglycerate mutase |
| D6RBV2 | LMAN2 | 0,234035059 | 2 | 0,005159618 | 0,003749678 | Vesicular integral-membrane protein VIP36 |
| A0A0G2JJA7 | PSMB9 | 0,235352012 | 3 | 8,82E-06 | 1,12E-05 | Proteasome subunit beta |
| E7EMS2 | NPC2 | 0,237105218 | 2 | 7,34E-06 | 9,45E-06 | NPC intracellular cholesterol transporter 2 |
| A0A0G2JH68 | DIAPH1 | 0,244384274 | 8 | 0,004513287 | 0,003329596 | Protein diaphanous homolog 1 |
| Q00013 | MPP1 | 0,24442417 | 2 | 0,057782471 | 0,031517712 | 55 kDa erythrocyte membrane protein |
| P63261 | ACTG1 | 0,244500713 | 33 | 1,73E-34 | 1,85E-33 | Actin, cytoplasmic 2 |
| Q13177 | PAK2 | 0,245916513 | 2 | 0,006444611 | 0,004601957 | Serine/threonine-protein kinase PAK 2 |
| P37837 | TALDO1 | 0,246403881 | 15 | 2,31E-11 | 6,17E-11 | Transaldolase |
| P07195 | LDHB | 0,247567448 | 14 | 7,97E-10 | 1,77E-09 | L-lactate dehydrogenase B chain |
| P51149 | RAB7A | 0,248091197 | 9 | 2,69E-32 | 2,70E-31 | Ras-related protein Rab-7a |
| P08754 | GNAI3 | 0,248594012 | 4 | 0,006930216 | 0,00487792 | Guanine nucleotide-binding protein G(k) subunit alpha |
| H0YHA7 | RPL18 | 0,248727472 | 6 | 7,18E-08 | 1,24E-07 | 60S ribosomal protein L18 (Fragment) |
| E9PGZ4 | SACM1L | 0,252504888 | 2 | 0,000445208 | 0,000418819 | Phosphatidylinositide phosphatase SAC1 |
| P61006 | RAB8A | 0,253193347 | 4 | 1,00E-05 | 1,27E-05 | Ras-related protein Rab-8A |

| | | | | | | |
|---|---|---|---|---|---|---|
| A0A2R8YC S7 | SAMHD1 | 0,2561892 06 | 8 | 1,43E-05 | 1,73E-05 | Deoxynucleoside triphosphate triphosphohydrolase SAMHD1 |
| H0Y6E7 | RBMX | 0,2586429 95 | 4 | 0,000163713 | 0,0001653 94 | RNA-binding motif protein, X chromosome (Fragment) |
| P98160 | HSPG2 | 0,2598345 56 | 68 | 1,18E-37 | 1,45E-36 | Basement membrane-specific heparan sulfate proteoglycan core protein |
| O15230 | LAMA5 | 0,2611434 24 | 11 | 0,000313103 | 0,0003050 43 | Laminin subunit alpha-5 |
| P53634 | CTSC | 0,2648390 51 | 4 | 1,15E-06 | 1,70E-06 | Dipeptidyl peptidase 1 |
| J3QRS3 | MYL12A | 0,2659215 33 | 3 | 7,58E-09 | 1,50E-08 | Myosin regulatory light chain 12A |
| P12268 | IMPDH2 | 0,2774247 51 | 9 | 8,06E-10 | 1,78E-09 | Inosine-5'-monophosphate dehydrogenase 2 |
| D3YTB1 | RPL32 | 0,2786528 74 | 3 | 0,001265476 | 0,0010884 86 | 60S ribosomal protein L32 (Fragment) |
| Q14764 | MVP | 0,2789745 01 | 18 | 3,71E-14 | 1,23E-13 | Major vault protein |
| P68871 | HBB | 0,2893143 58 | 17 | 1,21E-23 | 7,55E-23 | Hemoglobin subunit beta |
| J3KTA4 | DDX5 | 0,2926327 04 | 10 | 3,94E-23 | 2,39E-22 | Probable ATP-dependent RNA helicase DDX5 |
| A0A1W2PP H1 | ME2 | 0,2964871 84 | 3 | 1,90E-07 | 3,13E-07 | Malic enzyme |
| A0A2R8Y48 4 | CD47 | 0,2968481 34 | 2 | 1,24E-05 | 1,52E-05 | Leukocyte surface antigen CD47 (Fragment) |
| P27824 | CANX | 0,2968934 98 | 12 | 3,26E-15 | 1,15E-14 | Calnexin |
| C9JHR8 | CD163 | 0,2997071 8 | 4 | 0,000633893 | 0,0005829 45 | Scavenger receptor cysteine-rich type 1 protein M130 |
| P62834 | RAP1A | 0,2997088 41 | 3 | 0,000458568 | 0,0004297 43 | Ras-related protein Rap-1A |
| P07476 | IVL | 0,3001572 54 | 19 | 1,85E-18 | 8,50E-18 | Involucrin |
| Q01518 | CAP1 | 0,3016346 07 | 20 | 4,03E-26 | 2,72E-25 | Adenylyl cyclase-associated protein 1 |
| P63096 | GNAI1 | 0,3059826 48 | 7 | 0,00291671 | 0,0022634 41 | Guanine nucleotide-binding protein G(i) subunit alpha-1 |
| P61160 | ACTR2 | 0,3067344 76 | 13 | 2,51E-21 | 1,36E-20 | Actin-related protein 2 |
| P06899 | HIST1H2BJ | 0,3075735 51 | 2 | 5,25E-08 | 9,25E-08 | Histone H2B type 1-J |
| P52907 | CAPZA1 | 0,3101683 57 | 8 | 0,000129655 | 0,0001345 79 | F-actin-capping protein subunit alpha-1 |
| P08519 | LPA | 0,3146830 92 | 3 | 0,004379739 | 0,0032403 48 | Apolipoprotein(a) |
| Q14444 | CAPRIN1 | 0,3163454 24 | 4 | 1,14E-07 | 1,92E-07 | Caprin-1 |
| P62873 | GNB1 | 0,3171480 66 | 4 | 5,37E-07 | 8,21E-07 | Guanine nucleotide-binding protein G(I)/G(S)/G(T) subunit beta-1 |
| O95716 | RAB3D | 0,3177557 51 | 2 | 2,22E-06 | 3,15E-06 | Ras-related protein Rab-3D |
| Q92522 | H1FX | 0,3209239 93 | 2 | 0,001282032 | 0,0010988 84 | Histone H1x |
| A0A0A0MT S2 | GPI | 0,3281580 24 | 13 | 1,14E-16 | 4,67E-16 | Glucose-6-phosphate isomerase (Fragment) |
| O15144 | ARPC2 | 0,3311340 21 | 14 | 5,79E-28 | 4,25E-27 | Actin-related protein 2/3 complex subunit 2 |
| O75131 | CPNE3 | 0,3349286 38 | 12 | 2,42E-22 | 1,43E-21 | Copine-3 |
| P16401 | HIST1H1B | 0,3355597 01 | 5 | 0,000232453 | 0,0002305 78 | Histone H1.5 |
| P26038 | MSN | 0,3374762 05 | 24 | 5,23E-75 | 1,35E-73 | Moesin |
| P61158 | ACTR3 | 0,3376779 26 | 17 | 1,96E-28 | 1,51E-27 | Actin-related protein 3 |
| O95197-3 | RTN3 | 0,3376859 18 | 2 | 5,35E-07 | 8,20E-07 | Isoform 3 of Reticulon-3 |
| P50995 | ANXA11 | 0,3408306 65 | 11 | 9,21E-31 | 8,24E-30 | Annexin A11 |
| P31949 | S100A11 | 0,3430255 01 | 2 | 1,07E-05 | 1,34E-05 | Protein S100-A11 |
| A0A087X0Z 7 | DHRS7 | 0,3455753 88 | 3 | 4,44E-07 | 6,96E-07 | Dehydrogenase/reductase SDR family member 7 |
| P35579 | MYH9 | 0,3461971 63 | 96 | 2,24E-217 | 1,84E-215 | Myosin-9 |

| | | | | | | |
|---|---|---|---|---|---|---|
| A0A0G2JH46 | HLA-DRA | 0,360605673 | 3 | 5,43E-09 | 1,10E-08 | HLA class II histocompatibility antigen, DR alpha chain |
| P30520 | ADSS | 0,364053184 | 5 | 1,03E-12 | 3,12E-12 | Adenylosuccinate synthetase isozyme 2 |
| P61224 | RAP1B | 0,368958596 | 8 | 1,47E-06 | 2,13E-06 | Ras-related protein Rap-1b |
| Q16777 | HIST2H2AC | 0,370682255 | 5 | 3,79E-11 | 9,77E-11 | Histone H2A type 2-C |
| Q93084 | ATP2A3 | 0,374780608 | 5 | 0,000475369 | 0,000443798 | Sarcoplasmic/endoplasmic reticulum calcium ATPase 3 |
| Q16666 | IFI16 | 0,375260493 | 4 | 2,73E-05 | 3,12E-05 | Gamma-interferon-inducible protein 16 |
| F8WCF6 | ARPC4-TTLL3 | 0,377788576 | 8 | 3,56E-08 | 6,41E-08 | Actin-related protein 2/3 complex subunit 4 |

### Supplement Table 7. Dataset of mouse DIA-MS

**(bold font represents log2FC > |0.38|)**

| UNIPROT ID | Gene Name | AVG Log2 (INC/con) | N (number of unique total peptides) | P-value | BH (Q-value) | Name |
|---|---|---|---|---|---|---|
| **P04247** | **Mb** | **- 2,316215654** | **12** | **0,001416 36** | **0,002596 662** | **Myoglobin** |
| **P40936** | **Inmt** | **- 1,946711538** | **19** | **6,01E-05** | **0,000188 876** | **Indolethylamine N-methyltransferase** |
| **P14094** | **Atp1b1** | **- 1,807621235** | **36** | **0,007535 94** | **0,009845 803** | **Sodium/potassium-transporting ATPase subunit beta-1** |
| **P07310** | **Ckm** | **- 1,587404106** | **28** | **4,32E-06** | **1,91E-05** | **Creatine kinase M-type** |
| **P05977** | **Myl1** | **- 1,580821889** | **18** | **4,02E-05** | **0,000132 22** | **Myosin light chain 1/3, skeletal muscle isoform** |
| **A2A6J8** | **Tnni2** | **- 1,548019405** | **5** | **0,007022 87** | **0,009354 68** | **Troponin I, fast skeletal muscle (Fragment)** |
| **P16015** | **Ca3** | **- 1,521357466** | **22** | **1,70E-05** | **6,10E-05** | **Carbonic anhydrase 3** |
| **A2A6H6** | **Tnnt3** | **-1,45608353** | **12** | **0,002681 62** | **0,004323 557** | **Troponin T, fast skeletal muscle** |
| **E9PYJ9** | **Ldb3** | **- 1,446010395** | **17** | **0,029894 21** | **0,028715 282** | **LIM domain-binding protein 3** |
| **Q9R0Y5** | **Ak1** | **- 1,414441789** | **21** | **4,84E-06** | **2,12E-05** | **Adenylate kinase isoenzyme 1** |
| **Q5SX40** | **Myh1** | **- 1,409960911** | **157** | **3,68E-23** | **1,93E-21** | **Myosin-1** |
| **Q8R429** | **Atp2a1** | **- 1,402018974** | **54** | **0,027220 17** | **0,026749 504** | **Sarcoplasmic/endoplasmic reticulum calcium ATPase 1** |
| **Q62234** | **Myom1** | **- 1,390645477** | **46** | **0,003433 36** | **0,005215 035** | **Myomesin-1** |
| **P04117** | **Fabp4** | **-1,32630119** | **14** | **7,06E-05** | **0,000217 96** | **Fatty acid-binding protein, adipocyte** |
| **P58771** | **Tpm1** | **- 1,321883271** | **27** | **0,000642 72** | **0,001348 731** | **Tropomyosin alpha-1 chain** |
| **Q9QYG0** | **Ndrg2** | **- 1,313600727** | **28** | **0,010201 64** | **0,012536 07** | **Protein NDRG2** |
| **Q9WUB3** | **Pygm** | **- 1,259739917** | **81** | **0,000119 37** | **0,000329 605** | **Glycogen phosphorylase, muscle form** |
| **P97457** | **Mylpf** | **- 1,233586774** | **17** | **0,020535 91** | **0,021513 815** | **Myosin regulatory light chain 2, skeletal muscle isoform** |
| **Q9CQE8** | **NaN** | **- 1,198689885** | **24** | **0,035277 06** | **0,032646 006** | **UPF0568 protein C14orf166 homolog** |
| **P21812** | **Mcpt4** | **-1,19468189** | **13** | **1,37E-05** | **5,12E-05** | **Mast cell protease 4** |
| **Q9JI91** | **Actn2** | **- 1,181698061** | **69** | **6,81E-05** | **0,000211 166** | **Alpha-actinin-2** |
| **F6RT34** | **Mbp** | **-1,12024016** | **11** | **5,97E-10** | **6,47E-09** | **Myelin basic protein (Fragment)** |
| **Q62425** | **Ndufa4** | **- 1,092310562** | **12** | **0,004783 83** | **0,006956 45** | **Cytochrome c oxidase subunit NDUFA4** |
| **Q04447** | **Ckb** | **- 1,083037953** | **48** | **6,57E-10** | **6,89E-09** | **Creatine kinase B-type** |
| **G5E846** | **Prph** | **- 1,061919326** | **76** | **7,95E-05** | **0,000243 025** | **Peripherin** |
| **A2AEX6** | **Fhl1** | **- 1,055485892** | **26** | **0,039036 74** | **0,035497 412** | **Four and a half LIM domains protein 1** |
| **A0A087WP F8** | **Abhd14b** | **- 1,051646048** | **5** | **8,22E-06** | **3,28E-05** | **Alpha/beta hydrolase domain-containing protein 14B (Fragment)** |
| **P00329** | **Adh1** | **- 1,043509488** | **25** | **2,76E-08** | **2,07E-07** | **Alcohol dehydrogenase 1** |
| **P47857** | **Pfkm** | **-1,01386263** | **59** | **0,020596 72** | **0,021544 422** | **ATP-dependent 6-phosphofructokinase, muscle type** |
| **Q8CI43** | **Myl6b** | **- 1,009397251** | **21** | **0,009478** | **0,011992 569** | **Myosin light chain 6B** |
| **Q9R0P9** | **Uchl1** | **- 1,007432671** | **23** | **0,001018 49** | **0,001978 949** | **Ubiquitin carboxyl-terminal hydrolase isozyme L1** |
| **E9QNA7** | **Sorbs1** | **- 1,001357898** | **8** | **0,043250 92** | **0,038364 539** | **Sorbin and SH3 domain-containing protein 1** |
| **Q9CQ60** | **Pgls** | **- 0,996689133** | **22** | **0,000194 59** | **0,000484 206** | **6-phosphogluconolactonase** |
| **P48962** | **Slc25a4** | **- 0,977182377** | **32** | **0,001635 94** | **0,002890 445** | **ADP/ATP translocase 1** |

| | | | | | | |
|---|---|---|---|---|---|---|
| O70423 | Aoc3 | -0,954033705 | 29 | 0,03249818 | 0,030783 | Membrane primary amine oxidase |
| P15089 | Cpa3 | -0,949319167 | 17 | 3,90E-06 | 1,78E-05 | Mast cell carboxypeptidase A |
| Q9DB20 | Atp5o | -0,942825363 | 20 | 0,00012554 | 0,000343857 | ATP synthase subunit O, mitochondrial |
| Q9CPQ8 | Atp5l | -0,940728274 | 14 | 0,02095836 | 0,021822297 | ATP synthase subunit g, mitochondrial |
| Q09PK2 | Asprv1 | -0,933629653 | 8 | 0,00202499 | 0,003452611 | Retroviral-like aspartic protease 1 |
| P21550 | Eno3 | -0,919798483 | 29 | 0,00275658 | 0,004413108 | Beta-enolase |
| Q922B1 | Macrod1 | -0,917837407 | 18 | 0,06165215 | 0,049936774 | O-acetyl-ADP-ribose deacetylase MACROD1 |
| P16110 | Lgals3 | -0,917659024 | 15 | 2,97E-14 | 6,76E-13 | Galectin-3 |
| P19783 | Cox4i1 | -0,917251091 | 22 | 0,00255832 | 0,004144354 | Cytochrome c oxidase subunit 4 isoform 1, mitochondrial |
| P62631 | Eef1a2 | -0,916909802 | 27 | 0,06036911 | 0,049072386 | Elongation factor 1-alpha 2 |
| P05063 | Aldoc | -0,913493028 | 41 | 0,01241704 | 0,014651253 | Fructose-bisphosphate aldolase C |
| P08551 | Nefl | -0,898520823 | 95 | 1,46E-06 | 7,61E-06 | Neurofilament light polypeptide |
| Q9ERD7 | Tubb3 | -0,88099288 | 22 | 0,00103645 | 0,001996771 | Tubulin beta-3 chain |
| Q9ERS2 | Ndufa13 | -0,878551522 | 15 | 0,00674814 | 0,009023979 | NADH dehydrogenase [ubiquinone] 1 alpha subcomplex subunit 13 |
| A2AJ28 | Clic3 | -0,871081193 | 5 | 0,01326668 | 0,015413762 | Chloride intracellular channel protein |
| Q99M73 | Krt84 | -0,870718958 | 13 | 0,00071507 | 0,001468915 | Keratin, type II cuticular Hb4 |
| G3X9L6 | Gm10250 | -0,865388555 | 10 | 4,29E-07 | 2,55E-06 | MCG55033 |
| O35215 | Ddt | -0,864732774 | 17 | 3,18E-06 | 1,51E-05 | D-dopachrome decarboxylase |
| P08074 | Cbr2 | -0,864578188 | 19 | 1,81E-05 | 6,45E-05 | Carbonyl reductase [NADPH] 2 |
| Q64105 | Spr | -0,863111282 | 11 | 4,82E-11 | 6,57E-10 | Sepiapterin reductase |
| Q9DB60 | Fam213b | -0,863000951 | 17 | 0,00179267 | 0,00312685 | Prostamide/prostaglandin F synthase |
| P08553 | Nefm | -0,85486283 | 135 | 6,13E-08 | 4,40E-07 | Neurofilament medium polypeptide |
| Q9CQQ7 | Atp5f1 | -0,843607605 | 27 | 4,94E-05 | 0,000160422 | ATP synthase F(0) complex subunit B1, mitochondrial |
| Q64437 | Adh7 | -0,841178319 | 4 | 0,03551049 | 0,032749344 | Alcohol dehydrogenase class 4 mu/sigma chain |
| P31001 | Des | -0,837173618 | 44 | 2,87E-05 | 9,80E-05 | Desmin |
| O54724 | Ptrf | -0,832768345 | 30 | 2,96E-12 | 4,48E-11 | Polymerase I and transcript release factor |
| P60824 | Cirbp | -0,819795043 | 14 | 0,0019448 | 0,003340933 | Cold-inducible RNA-binding protein |
| D3YXP6 | Pmvk | -0,81912892 | 18 | 0,0049633 | 0,007081534 | Phosphomevalonate kinase |
| Q9CQD1 | Rab5a | -0,815695003 | 13 | 0,01037155 | 0,01269192 | Ras-related protein Rab-5A |
| Q9D967 | Mdp1 | -0,812603522 | 11 | 0,00037512 | 0,000861384 | Magnesium-dependent phosphatase 1 |
| P34884 | Mif | -0,806608118 | 12 | 0,00010675 | 0,000303341 | Macrophage migration inhibitory factor |
| O55026 | Entpd2 | -0,805817668 | 19 | 2,02E-05 | 7,14E-05 | Ectonucleoside triphosphate diphosphohydrolase 2 |
| Q8K0Y2 | Krt33a | -0,799039132 | 10 | 0,00817411 | 0,010558228 | Keratin, type I cuticular Ha3-I |
| P17563 | Selenbp1 | -0,785895712 | 30 | 2,74E-11 | 3,82E-10 | Selenium-binding protein 1 |
| P63168 | Dynll1 | -0,783978719 | 11 | 0,00033921 | 0,000792262 | Dynein light chain 1, cytoplasmic |
| Q9JHQ0 | Anxa9 | -0,777622672 | 16 | 0,04704961 | 0,040942152 | Annexin A9 |
| O08638 | Myh11 | -0,776550873 | 143 | 3,98E-05 | 0,000131923 | Myosin-11 |
| Q9CZ04 | Cops7a | -0,769538275 | 21 | 0,01856559 | 0,019980599 | COP9 signalosome complex subunit 7a |
| P05064 | Aldoa | -0,769192768 | 51 | 0,02586227 | 0,025786649 | Fructose-bisphosphate aldolase A |

| | | | | | | |
|---|---|---|---|---|---|---|
| O55103 | Prx | -0,765297303 | 89 | 6,85E-11 | 8,50E-10 | Periaxin |
| P19157 | Gstp1 | -0,741349659 | 25 | 3,32E-07 | 2,02E-06 | Glutathione S-transferase P 1 |
| O08970 | Tuft1 | -0,734062618 | 17 | 0,00042063 | 0,000946763 | Tuftelin |
| Q9D023 | Mpc2 | -0,733940449 | 12 | 0,04196793 | 0,037512622 | Mitochondrial pyruvate carrier 2 |
| Q9WU63 | Hebp2 | -0,701535495 | 9 | 0,00122935 | 0,002301233 | Heme-binding protein 2 |
| Q61495 | Dsg1a | -0,701044059 | 2 | 0,02757621 | 0,027021516 | Desmoglein-1-alpha |
| Q9CXS4 | Cenpv | -0,694278562 | 23 | 0,03936066 | 0,035744301 | Centromere protein V |
| Q8CA72 | Gan | -0,689542653 | 27 | 0,01451117 | 0,016412305 | Gigaxonin |
| Q9CQ69 | Uqcrq | -0,685238731 | 15 | 0,02603981 | 0,025887979 | Cytochrome b-c1 complex subunit 8 |
| Q3U3G8 | Serhl | -0,683939203 | 7 | 0,04868267 | 0,04197419 | Serine hydrolase-like protein |
| P61971 | Nutf2 | -0,683350113 | 8 | 1,67E-05 | 6,01E-05 | Nuclear transport factor 2 |
| O08709 | Prdx6 | -0,677484371 | 17 | 2,32E-06 | 1,15E-05 | Peroxiredoxin-6 |
| D3YYN7 | Atp1a2 | -0,67502127 | 41 | 0,00013588 | 0,000369203 | Sodium/potassium-transporting ATPase subunit alpha-2 |
| O55126 | Gbas | -0,673720611 | 28 | 0,05218268 | 0,04404528 | Protein NipSnap homolog 2 |
| Q924Y0 | Bbox1 | -0,673279704 | 21 | 0,02228057 | 0,022919081 | Gamma-butyrobetaine dioxygenase |
| P42125 | Eci1 | -0,665840981 | 21 | 0,00017952 | 0,000455127 | Enoyl-CoA delta isomerase 1, mitochondrial |
| P15626 | Gstm2 | -0,660907564 | 29 | 4,96E-10 | 5,45E-09 | Glutathione S-transferase Mu 2 |
| Q3V156 | Osbpl1a | -0,659718364 | 7 | 0,05340681 | 0,044746244 | Oxysterol-binding protein |
| P20357 | Map2 | -0,653072908 | 5 | 0,05240975 | 0,044127716 | Microtubule-associated protein 2 |
| P48774 | Gstm5 | -0,645361003 | 28 | 0,00082863 | 0,001676133 | Glutathione S-transferase Mu 5 |
| A0A0A0MQ97 | Nccrp1 | -0,642962661 | 6 | 0,00154112 | 0,002787916 | F-box only protein 50 |
| Q9CRB6 | Tppp3 | -0,642067836 | 22 | 0,03532662 | 0,032646006 | Tubulin polymerization-promoting protein family member 3 |
| Q62426 | Cstb | -0,635940126 | 12 | 1,96E-09 | 1,88E-08 | Cystatin-B |
| E9QK82 | Mpz | -0,635460911 | 16 | 0,00454313 | 0,006677611 | Myelin protein P0 |
| E9QNP3 | Hrnr | -0,629673256 | 4 | 0,00639276 | 0,00859934 | Hornerin |
| Q9WTP6 | Ak2 | -0,626516245 | 25 | 0,0001172 | 0,000327584 | Adenylate kinase 2, mitochondrial |
| F6RPJ9 | Ide | -0,622790893 | 71 | 1,06E-10 | 1,27E-09 | Insulin-degrading enzyme (Fragment) |
| Q9D695 | Serpinb7 | -0,621434142 | 9 | 0,05532719 | 0,045959979 | Serpin B7 |
| Q8CI94 | Pygb | -0,620819465 | 91 | 0,01618711 | 0,017863442 | Glycogen phosphorylase, brain form |
| Q6P1B9 | Bin1 | -0,613540985 | 15 | 0,01588022 | 0,017610259 | Bin1 protein |
| Q61205 | Pafah1b3 | -0,610669976 | 22 | 0,02052746 | 0,021513815 | Platelet-activating factor acetylhydrolase IB subunit gamma |
| O70492 | Snx3 | -0,609527472 | 21 | 0,00457018 | 0,006691358 | Sorting nexin-3 |
| Q63918 | Sdpr | -0,60733358 | 31 | 0,0058974 | 0,008108925 | Serum deprivation-response protein |
| Q8R0W0 | Eppk1 | -0,607239567 | 138 | 4,46E-42 | 7,60E-40 | Epiplakin |
| Q5SRX1 | Tom1l2 | -0,60592995 | 36 | 0,01167788 | 0,014021674 | TOM1-like protein 2 |
| Q60675 | Lama2 | -0,603510587 | 174 | 0,00168206 | 0,002949085 | Laminin subunit alpha-2 |
| P08228 | Sod1 | -0,602467672 | 19 | 3,73E-06 | 1,72E-05 | Superoxide dismutase [Cu-Zn] |
| G5E8K2 | Ank3 | -0,601984944 | 20 | 0,00042488 | 0,000952393 | Ankyrin 3, epithelial, isoform CRA_l |

| | | | | | | |
|---|---|---|---|---|---|---|
| Q8C605 | Pfkp | -0,599518715 | 32 | 3,79E-05 | 0,000126029 | ATP-dependent 6-phosphofructokinase |
| Q8K4L4 | Pof1b | -0,598551375 | 11 | 5,74E-07 | 3,29E-06 | Protein POF1B |
| Q99LX0 | Park7 | -0,597170925 | 24 | 6,08E-08 | 4,40E-07 | Protein deglycase DJ-1 |
| P16125 | Ldhb | -0,595212507 | 31 | 0,00560399 | 0,007815787 | L-lactate dehydrogenase B chain |
| Q61425 | Hadh | -0,592962122 | 32 | 0,00329219 | 0,005056921 | Hydroxyacyl-coenzyme A dehydrogenase, mitochondrial |
| D3Z030 | Lrrc16a | -0,589885744 | 3 | 0,03269524 | 0,030883873 | Leucine-rich repeat-containing protein 16A |
| P14152 | Mdh1 | -0,586005175 | 27 | 1,84E-10 | 2,09E-09 | Malate dehydrogenase, cytoplasmic |
| Q99JY8 | Ppap2b | -0,585651464 | 19 | 0,00254331 | 0,004129847 | Lipid phosphate phosphohydrolase 3 |
| P08207 | S100a10 | -0,585350967 | 10 | 0,04204353 | 0,037531003 | Protein S100-A10 |
| Q9WTP7 | Ak3 | -0,581996663 | 24 | 0,00206311 | 0,003508839 | GTP:AMP phosphotransferase AK3, mitochondrial |
| Q9QXX4 | Slc25a13 | -0,581957444 | 56 | 0,01995993 | 0,021044202 | Calcium-binding mitochondrial carrier protein Aralar2 |
| A3KMP2 | Ttc38 | -0,578155706 | 31 | 0,03016623 | 0,028854657 | Tetratricopeptide repeat protein 38 |
| O09164 | Sod3 | -0,57486319 | 10 | 6,79E-05 | 0,000211166 | Extracellular superoxide dismutase [Cu-Zn] |
| P45376 | Akr1b1 | -0,574659976 | 29 | 1,01E-05 | 3,94E-05 | Aldose reductase |
| A0A0G2JEK2 | Crip1 | -0,571070033 | 8 | 0,00109957 | 0,00208307 | Cysteine-rich protein 1 |
| Q8R5F8 | Eps8l1 | -0,570662433 | 9 | 0,0012762 | 0,002378052 | Epidermal growth factor receptor kinase substrate 8-like protein 1 |
| Q8R3B1 | Plcd1 | -0,567110877 | 47 | 6,89E-08 | 4,75E-07 | 1-phosphatidylinositol 4,5-bisphosphate phosphodiesterase delta-1 |
| Q61823 | Pdcd4 | -0,564970066 | 25 | 0,00047629 | 0,001044475 | Programmed cell death protein 4 |
| Q3UEB4 | Mvk | -0,564958155 | 9 | 0,00500241 | 0,007092811 | Mevalonate kinase |
| Q60854 | Serpinb6 | -0,563192462 | 39 | 2,62E-09 | 2,45E-08 | Serpin B6 |
| Q6P8X1 | Snx6 | -0,562783692 | 39 | 0,00210396 | 0,003569411 | Sorting nexin-6 |
| O88986 | Gcat | -0,557448796 | 21 | 0,00157168 | 0,002828189 | 2-amino-3-ketobutyrate coenzyme A ligase, mitochondrial |
| Q62000 | Ogn | -0,557066634 | 19 | 0,00216163 | 0,003631118 | Mimecan |
| Q9Z2Y8 | Prosc | -0,555407447 | 19 | 0,00240336 | 0,003933147 | Proline synthase co-transcribed bacterial homolog protein |
| P32020 | Scp2 | -0,552753288 | 39 | 0,00986222 | 0,012287924 | Non-specific lipid-transfer protein |
| Q9WTR5 | Cdh13 | -0,552317423 | 27 | 0,00361117 | 0,005448708 | Cadherin-13 |
| Q9D0F9 | Pgm1 | -0,551445154 | 56 | 0,00423711 | 0,006295657 | Phosphoglucomutase-1 |
| P61089 | Ube2n | -0,550992373 | 13 | 0,0009391 | 0,00185642 | Ubiquitin-conjugating enzyme E2 N |
| O88587 | Comt | -0,550992197 | 22 | 0,00016651 | 0,000431737 | Catechol O-methyltransferase |
| P17751 | Tpi1 | -0,550217779 | 28 | 0,00774378 | 0,01005954 | Triosephosphate isomerase |
| Q3UW66 | Mpst | -0,550033422 | 20 | 0,00600608 | 0,008170638 | Sulfurtransferase |
| Q9JJU8 | Sh3bgrl | -0,545686914 | 14 | 0,00015968 | 0,00041884 | SH3 domain-binding glutamic acid-rich-like protein |
| E0CYV0 | Pcmt1 | -0,540709588 | 8 | 0,01773969 | 0,019295807 | Protein-L-isoaspartate O-methyltransferase |
| Q91ZU6 | Dst | -0,539599154 | 33 | 5,19E-05 | 0,000166872 | Dystonin |
| G3UWG1 | Gm10108 | -0,536034431 | 4 | 0,01278613 | 0,014983378 | MCG115977 |
| Q8CIB5 | Fermt2 | -0,534703068 | 49 | 0,01175499 | 0,014040113 | Fermitin family homolog 2 |
| P17879 | Hspa1b | -0,532421368 | 17 | 2,71E-12 | 4,21E-11 | Heat shock 70 kDa protein 1B |
| Q9ERE2 | Krt81 | -0,532086905 | 10 | 0,00569374 | 0,00787653 | Keratin, type II cuticular Hb1 |

| | | | | | | |
|---|---|---|---|---|---|---|
| Q921K8 | Tcaf2 | -0,531722824 | 20 | 2,09E-06 | 1,06E-05 | TRPM8 channel-associated factor 2 |
| E9Q174 | Myo6 | -0,530971762 | 16 | 0,01784657 | 0,019381146 | Unconventional myosin-VI |
| P33267 | Cyp2f2 | -0,530055737 | 37 | 0,00521 | 0,007356558 | Cytochrome P450 2F2 |
| P14602 | Hspb1 | -0,528652218 | 22 | 1,93E-08 | 1,48E-07 | Heat shock protein beta-1 |
| Q99LC3 | Ndufa10 | -0,527736127 | 37 | 0,03706069 | 0,033972301 | NADH dehydrogenase [ubiquinone] 1 alpha subcomplex subunit 10, mitochondrial |
| P70227 | Itpr3 | -0,527039942 | 106 | 0,00077513 | 0,001582754 | Inositol 1,4,5-trisphosphate receptor type 3 |
| Q8BH59 | Slc25a12 | -0,526833943 | 50 | 0,00917984 | 0,011702147 | Calcium-binding mitochondrial carrier protein Aralar1 |
| P05201 | Got1 | -0,526484213 | 45 | 0,0004607 | 0,001013544 | Aspartate aminotransferase, cytoplasmic |
| A0A0A0MQF6 | Gapdh | -0,525135081 | 22 | 0,00482222 | 0,006992827 | Glyceraldehyde-3-phosphate dehydrogenase |
| Q8VDN2 | Atp1a1 | -0,522178706 | 62 | 1,27E-06 | 6,78E-06 | Sodium/potassium-transporting ATPase subunit alpha-1 |
| Q9D0J4 | Arl2 | -0,517900955 | 13 | 0,02188351 | 0,02257875 | ADP-ribosylation factor-like protein 2 |
| P28654 | Dcn | -0,509221388 | 27 | 0,00011925 | 0,000329605 | Decorin |
| P97350 | Pkp1 | -0,50862807 | 15 | 0,00312516 | 0,004849168 | Plakophilin-1 |
| Q8R1G6 | Pdlim2 | -0,508626022 | 18 | 0,00022832 | 0,000556133 | PDZ and LIM domain protein 2 |
| P10649 | Gstm1 | -0,507274534 | 28 | 2,48E-07 | 1,55E-06 | Glutathione S-transferase Mu 1 |
| Q9JK53 | Prelp | -0,505264847 | 22 | 2,61E-07 | 1,60E-06 | Prolargin |
| P70271 | Pdlim4 | -0,504618189 | 27 | 0,00490795 | 0,007046782 | PDZ and LIM domain protein 4 |
| O88322 | Nid2 | -0,503412526 | 64 | 0,00107127 | 0,00204651 | Nidogen-2 |
| Q8BH95 | Echs1 | -0,501808228 | 24 | 0,00016712 | 0,000431737 | Enoyl-CoA hydratase, mitochondrial |
| Q99KI0 | Aco2 | -0,501178973 | 75 | 2,04E-05 | 7,16E-05 | Aconitate hydratase, mitochondrial |
| Q99K30 | Eps8l2 | -0,500833484 | 51 | 5,26E-06 | 2,27E-05 | Epidermal growth factor receptor kinase substrate 8-like protein 2 |
| D3Z6Y9 | Dohh | -0,500580574 | 2 | 0,00227044 | 0,003767489 | Deoxyhypusine hydroxylase |
| Q91V64 | Isoc1 | -0,49875091 | 18 | 0,00900088 | 0,011510423 | Isochorismatase domain-containing protein 1 |
| Q9JII6 | Akr1a1 | -0,49782973 | 34 | 7,71E-10 | 7,73E-09 | Alcohol dehydrogenase [NADP(+)] |
| G5E898 | Ppl | -0,497004244 | 80 | 5,43E-31 | 3,71E-29 | Periplakin |
| O09044 | Snap23 | -0,491975366 | 21 | 0,02253885 | 0,023080321 | Synaptosomal-associated protein 23 |
| P70296 | Pebp1 | -0,485727859 | 19 | 0,00020794 | 0,000510115 | Phosphatidylethanolamine-binding protein 1 |
| Q8BH64 | Ehd2 | -0,485283274 | 49 | 1,34E-13 | 2,69E-12 | EH domain-containing protein 2 |
| P97315 | Csrp1 | -0,484280995 | 16 | 0,00873311 | 0,011195451 | Cysteine and glycine-rich protein 1 |
| Q8VED9 | Lgalsl | -0,483454401 | 14 | 0,00293826 | 0,004627924 | Galectin-related protein |
| Q61239 | Fnta | -0,483055271 | 25 | 0,00051505 | 0,001122247 | Protein farnesyltransferase/geranylgeranyltransferase type-1 subunit alpha |
| Q9D172 | D10Jhu81e | -0,481371028 | 17 | 0,00039576 | 0,000899698 | ES1 protein homolog, mitochondrial |
| Q61753 | Phgdh | -0,479913613 | 35 | 1,87E-06 | 9,52E-06 | D-3-phosphoglycerate dehydrogenase |
| Q9EPB4 | Pycard | -0,479579773 | 16 | 6,14E-11 | 7,91E-10 | Apoptosis-associated speck-like protein containing a CARD |
| P01942 | Hba | -0,478372907 | 13 | 0,00485532 | 0,007015532 | Hemoglobin subunit alpha |
| Q9Z0S1 | Bpnt1 | -0,478063677 | 23 | 1,29E-06 | 6,79E-06 | 3'(2'),5'-bisphosphate nucleotidase 1 |
| Q64314 | Cd34 | -0,476885041 | 14 | 0,00020134 | 0,000495723 | Hematopoietic progenitor cell antigen CD34 |
| B9EHJ3 | Tjp1 | -0,476870204 | 18 | 0,01824832 | 0,019754526 | Tight junction protein ZO-1 |

| | | | | | | |
|---|---|---|---|---|---|---|
| Q9DCW4 | Etfb | -0,476135725 | 23 | 2,50E-07 | 1,55E-06 | Electron transfer flavoprotein subunit beta |
| E9Q557 | Dsp | -0,4744716 | 176 | 5,98E-47 | 2,04E-44 | Desmoplakin |
| Q9EQ20 | Aldh6a1 | -0,472484554 | 49 | 0,00092039 | 0,001834658 | Methylmalonate-semialdehyde dehydrogenase [acylating], mitochondrial |
| Q61830 | Mrc1 | -0,471363298 | 96 | 2,11E-07 | 1,35E-06 | Macrophage mannose receptor 1 |
| O08553 | Dpysl2 | -0,47040709 | 53 | 5,52E-07 | 3,22E-06 | Dihydropyrimidinase-related protein 2 |
| Q9CPU0 | Glo1 | -0,470401747 | 17 | 5,72E-09 | 4,87E-08 | Lactoylglutathione lyase |
| Q9JLY7 | Dusp14 | -0,46874511 | 7 | 0,00159283 | 0,002845462 | Dual specificity protein phosphatase 14 |
| Q9D2Q8 | S100a14 | -0,467569589 | 3 | 0,05004635 | 0,042717912 | Protein S100-A14 |
| Q02248 | Ctnnb1 | -0,466522421 | 54 | 7,16E-09 | 5,89E-08 | Catenin beta-1 |
| Q9EPL9 | Acox3 | -0,466056194 | 45 | 0,0127864 | 0,014983378 | Peroxisomal acyl-coenzyme A oxidase 3 |
| Q8BTY1 | Ccbl1 | -0,465055233 | 18 | 2,21E-06 | 1,11E-05 | Kynurenine--oxoglutarate transaminase 1 |
| Q8BVI4 | Qdpr | -0,459408299 | 19 | 4,67E-09 | 4,11E-08 | Dihydropteridine reductase |
| P70124 | Serpinb5 | -0,45598468 | 13 | 0,00142548 | 0,002606374 | Serpin B5 |
| G3XA48 | Idi1 | -0,453469688 | 8 | 0,0435734 | 0,038543525 | Isopentenyl-diphosphate Delta-isomerase 1 |
| Q9CQM5 | Txndc17 | -0,452670001 | 14 | 0,03122635 | 0,029702052 | Thioredoxin domain-containing protein 17 |
| O08800 | Serpinb8 | -0,452016377 | 19 | 0,00034288 | 0,000798111 | Serpin B8 |
| P29452 | Casp1 | -0,451973887 | 30 | 0,04164934 | 0,03727671 | Caspase-1 |
| P49817 | Cav1 | -0,450243007 | 14 | 0,00092271 | 0,001834658 | Caveolin-1 |
| B1AQF4 | Dusp3 | -0,449305399 | 5 | 0,00136068 | 0,002501303 | Dual-specificity protein phosphatase 3 |
| Q9JM76 | Arpc3 | -0,446752785 | 23 | 0,00997546 | 0,012379906 | Actin-related protein 2/3 complex subunit 3 |
| Q8K354 | Cbr3 | -0,446256113 | 28 | 1,94E-11 | 2,75E-10 | Carbonyl reductase [NADPH] 3 |
| P70202 | Lxn | -0,444734499 | 12 | 0,00922231 | 0,011734354 | Latexin |
| Q03265 | Atp5a1 | -0,442099222 | 70 | 2,41E-05 | 8,39E-05 | ATP synthase subunit alpha, mitochondrial |
| A0A0A6YX73 | Prkar2a | -0,440817048 | 10 | 0,00011233 | 0,00031526 | cAMP-dependent protein kinase type II-alpha regulatory subunit |
| Q8CF02 | Fam25c | -0,439505167 | 4 | 0,02766364 | 0,027029512 | Protein FAM25C |
| P28474 | Adh5 | -0,43821906 | 28 | 5,13E-05 | 0,000165863 | Alcohol dehydrogenase class-3 |
| Q9CR62 | Slc25a11 | -0,433883549 | 30 | 0,01442205 | 0,01636579 | Mitochondrial 2-oxoglutarate/malate carrier protein |
| Q9D051 | Pdhb | -0,432805953 | 24 | 1,82E-05 | 6,47E-05 | Pyruvate dehydrogenase E1 component subunit beta, mitochondrial |
| O08599 | Stxbp1 | -0,432621418 | 78 | 0,00743 | 0,009763503 | Syntaxin-binding protein 1 |
| Q923D2 | Blvrb | -0,431280088 | 19 | 0,00668869 | 0,00896205 | Flavin reductase (NADPH) |
| B1AZ46 | Baiap2 | -0,431050033 | 57 | 0,02973774 | 0,028605273 | Brain-specific angiogenesis inhibitor 1-associated protein 2 |
| Q9D6R2 | Idh3a | -0,428023684 | 34 | 0,00482936 | 0,006992827 | Isocitrate dehydrogenase [NAD] subunit alpha, mitochondrial |
| Q02257 | Jup | -0,427886772 | 49 | 1,57E-12 | 2,55E-11 | Junction plakoglobin |
| Q9D8U8 | Snx5 | -0,427612905 | 39 | 7,98E-06 | 3,20E-05 | Sorting nexin-5 |
| D3YU60 | Mgst1 | -0,427292747 | 3 | 0,01669 | 0,018299973 | Microsomal glutathione S-transferase 1 |
| Q9CWK8 | Snx2 | -0,426048278 | 37 | 5,64E-11 | 7,40E-10 | Sorting nexin-2 |
| P02088 | Hbb-b1 | -0,426019917 | 20 | 0,00748841 | 0,009821342 | Hemoglobin subunit beta-1 |
| P99029 | Prdx5 | -0,424893117 | 17 | 0,02384814 | 0,024131203 | Peroxiredoxin-5, mitochondrial |

| | | | | | | |
|---|---|---|---|---|---|---|
| Q91VJ2 | Prkcdbp | -0,424428066 | 18 | 0,00015184 | 0,000401377 | Protein kinase C delta-binding protein |
| G5E850 | Cyb5a | -0,418462175 | 6 | 0,00280563 | 0,004470655 | Cytochrome b-5, isoform CRA_a |
| Q3TCD4 | Eci2 | -0,41733201 | 27 | 0,03988623 | 0,036125379 | Enoyl-CoA delta isomerase 2, mitochondrial |
| Q9D952 | Evpl | -0,416447391 | 27 | 8,38E-09 | 6,73E-08 | Envoplakin |
| Q11136 | Pepd | -0,414681863 | 30 | 0,00287144 | 0,004543671 | Xaa-Pro dipeptidase |
| P10833 | Rras | -0,414654009 | 17 | 0,01428805 | 0,016267862 | Ras-related protein R-Ras |
| Q8K1J6 | Trnt1 | -0,41332469 | 38 | 0,01350867 | 0,015588685 | CCA tRNA nucleotidyltransferase 1, mitochondrial |
| E9Q0J5 | Kif21a | -0,410701017 | 11 | 0,03402515 | 0,031787878 | Kinesin-like protein |
| Q8JZN5 | Acad9 | -0,409911406 | 50 | 0,01720468 | 0,018834015 | Acyl-CoA dehydrogenase family member 9, mitochondrial |
| P35492 | Hal | -0,409411945 | 40 | 0,00024669 | 0,000594496 | Histidine ammonia-lyase |
| P45952 | Acadm | -0,40904523 | 32 | 5,69E-06 | 2,40E-05 | Medium-chain specific acyl-CoA dehydrogenase, mitochondrial |
| Q8K4X7 | Agpat4 | -0,406889501 | 22 | 0,05643691 | 0,046541683 | 1-acyl-sn-glycerol-3-phosphate acyltransferase delta |
| Q9D832 | Dnajb4 | -0,406504986 | 32 | 0,00517894 | 0,007327878 | DnaJ homolog subfamily B member 4 |
| P53996 | Cnbp | -0,403728368 | 14 | 0,02824097 | 0,027514774 | Cellular nucleic acid-binding protein |
| Q99L13 | Hibadh | -0,402353389 | 24 | 0,01395948 | 0,016027553 | 3-hydroxyisobutyrate dehydrogenase, mitochondrial |
| O54974 | Lgals7 | -0,401785104 | 4 | 0,00271235 | 0,004352522 | Galectin-7 |
| D3Z7U4 | Mecp2 | -0,401393459 | 20 | 0,03984222 | 0,036125379 | Methyl-CpG-binding protein 2 |
| P35486 | Pdha1 | -0,398642784 | 44 | 0,04906526 | 0,042197358 | Pyruvate dehydrogenase E1 component subunit alpha, somatic form, mitochondrial |
| Q9JJI8 | Rpl38 | -0,398061441 | 6 | 0,05699092 | 0,046847251 | 60S ribosomal protein L38 |
| Q9R069 | Bcam | -0,398036985 | 32 | 0,0005443 | 0,001167344 | Basal cell adhesion molecule |
| O09131 | Gsto1 | -0,397863435 | 29 | 1,37E-05 | 5,12E-05 | Glutathione S-transferase omega-1 |
| Q61206 | Pafah1b2 | -0,396980661 | 20 | 0,05839781 | 0,047869362 | Platelet-activating factor acetylhydrolase IB subunit beta |
| Q9D3I6 | Krtap7-1 | -0,394857602 | 2 | 0,04795705 | 0,041432742 | Keratin-associated protein 7-1 |
| Q8JZK9 | Hmgcs1 | -0,394635969 | 37 | 1,51E-05 | 5,49E-05 | Hydroxymethylglutaryl-CoA synthase, cytoplasmic |
| P17439 | Gba | -0,391543493 | 25 | 0,00499344 | 0,007092811 | Glucosylceramidase |
| P28271 | Aco1 | -0,390724353 | 72 | 3,08E-05 | 0,000103633 | Cytoplasmic aconitate hydratase |
| G3X8T3 | Ctsa | -0,389509858 | 26 | 0,00103254 | 0,001994874 | Carboxypeptidase |
| Q924C1 | Xpo5 | -0,388904935 | 63 | 0,05592599 | 0,046255802 | Exportin-5 |
| P54822 | Adsl | -0,387631515 | 44 | 0,05149907 | 0,043630267 | Adenylosuccinate lyase |
| P70404 | Idh3g | -0,387461428 | 26 | 0,02618952 | 0,025998914 | Isocitrate dehydrogenase [NAD] subunit gamma 1, mitochondrial |
| Q91VH6 | Memo1 | -0,386613868 | 21 | 0,00148882 | 0,002714901 | Protein MEMO1 |
| Q9CPY7 | Lap3 | -0,382542604 | 46 | 0,00017217 | 0,00043978 | Cytosol aminopeptidase |
| O88736 | Hsd17b7 | -0,382454626 | 18 | 0,06163572 | 0,049936774 | 3-keto-steroid reductase |
| Q9DC69 | Ndufa9 | -0,382395259 | 37 | 0,00565564 | 0,007871723 | NADH dehydrogenase [ubiquinone] 1 alpha subcomplex subunit 9, mitochondrial |
| G3UVV4 | Hk1 | -0,381643162 | 83 | 0,00311187 | 0,004845418 | Hexokinase |
| Q6P069 | Sri | 0,387125046 | 18 | 0,01066351 | 0,01291743 | Sorcin |
| Q93092 | Taldo1 | 0,394061784 | 36 | 0,00027301 | 0,000652412 | Transaldolase |

| | | | | | | |
|---|---|---|---|---|---|---|
| P83882 | Rpl36a | 0,396505117 | 13 | 0,014904 64 | 0,016773 873 | 60S ribosomal protein L36a |
| Q8CIN4 | Pak2 | 0,397224945 | 30 | 1,42E-05 | 5,24E-05 | Serine/threonine-protein kinase PAK 2 |
| Q8BH61 | F13a1 | 0,400530852 | 52 | 9,77E-06 | 3,83E-05 | Coagulation factor XIII A chain |
| A2AQ53 | Fbn1 | 0,403425482 | 56 | 2,92E-06 | 1,41E-05 | Fibrillin-1 |
| P70335 | Rock1 | 0,403767557 | 104 | 0,055954 6 | 0,046255 802 | Rho-associated protein kinase 1 |
| H3BL37 | Tcof1 | 0,404648863 | 70 | 0,012689 97 | 0,014921 65 | Treacle protein |
| Q8VCI0 | Plbd1 | 0,410446464 | 24 | 0,044790 62 | 0,039301 275 | Phospholipase B-like 1 |
| P17932 | Rpl32-ps | 0,411184143 | 5 | 0,038061 47 | 0,034703 105 | Putative 60S ribosomal protein L32' |
| Q9QZQ8 | H2afy | 0,413194928 | 32 | 0,024391 24 | 0,024535 141 | Core histone macro-H2A.1 |
| P97310 | Mcm2 | 0,413338255 | 64 | 8,34E-05 | 0,000249 375 | DNA replication licensing factor MCM2 |
| Q7TNV0 | Dek | 0,413718857 | 39 | 0,010001 95 | 0,012379 906 | Protein DEK |
| O88342 | Wdr1 | 0,414686246 | 58 | 3,03E-08 | 2,25E-07 | WD repeat-containing protein 1 |
| P11031 | Sub1 | 0,417912195 | 21 | 1,39E-05 | 5,15E-05 | Activated RNA polymerase II transcriptional coactivator p15 |
| O09172 | Gclm | 0,419671717 | 18 | 0,005678 15 | 0,007876 53 | Glutamate--cysteine ligase regulatory subunit |
| TRYP_PIG | NaN | 0,423958904 | 9 | 2,90E-05 | 9,83E-05 | Trypsin - Sus scrofa (Pig). |
| Q6WVG3 | Kctd12 | 0,424735652 | 30 | 0,000182 38 | 0,000460 676 | BTB/POZ domain-containing protein KCTD12 |
| B1ATI0 | Aldh3a2 | 0,429523824 | 34 | 0,006855 5 | 0,009149 613 | Aldehyde dehydrogenase |
| P24547 | Impdh2 | 0,436181294 | 41 | 0,000552 61 | 0,001177 748 | Inosine-5'-monophosphate dehydrogenase 2 |
| Q569Z5 | Ddx46 | 0,44316019 | 84 | 0,011623 45 | 0,013980 936 | Probable ATP-dependent RNA helicase DDX46 |
| P70248 | Myo1f | 0,45009878 | 16 | 0,044351 75 | 0,039079 969 | Unconventional myosin-If |
| A2AJI0 | Map7d1 | 0,451549335 | 9 | 0,006034 15 | 0,008170 638 | MAP7 domain-containing protein 1 |
| Q61792 | Lasp1 | 0,455667245 | 26 | 0,034100 69 | 0,031814 869 | LIM and SH3 domain protein 1 |
| E9Q414 | Apob | 0,458298292 | 140 | 0,007348 02 | 0,009693 137 | Apolipoprotein B-100 |
| Q8BVY0 | Rsl1d1 | 0,459386092 | 32 | 6,98E-06 | 2,89E-05 | Ribosomal L1 domain-containing protein 1 |
| A2ASS6 | Ttn | 0,460708868 | 678 | 3,33E-15 | 9,48E-14 | Titin |
| P20029 | Hspa5 | 0,461487798 | 72 | 4,52E-09 | 4,06E-08 | 78 kDa glucose-regulated protein |
| Q9JJ28 | Flii | 0,462564874 | 75 | 0,012252 1 | 0,014509 179 | Protein flightless-1 homolog |
| P35564 | Canx | 0,466315726 | 80 | 1,50E-05 | 5,47E-05 | Calnexin |
| Q3TML0 | Pdia6 | 0,476907954 | 14 | 0,012854 25 | 0,015026 243 | Protein disulfide-isomerase A6 |
| P08113 | Hsp90b1 | 0,48047188 | 85 | 7,59E-06 | 3,08E-05 | Endoplasmin |
| Q00612 | G6pdx | 0,480895483 | 64 | 0,001026 09 | 0,001988 058 | Glucose-6-phosphate 1-dehydrogenase X |
| Q922Q8 | Lrrc59 | 0,484913902 | 23 | 0,001956 44 | 0,003352 487 | Leucine-rich repeat-containing protein 59 |
| E9Q5J9 | Tpm3 | 0,497981673 | 16 | 3,62E-07 | 2,19E-06 | Tropomyosin alpha-3 chain |
| P08003 | Pdia4 | 0,498912231 | 60 | 4,13E-06 | 1,85E-05 | Protein disulfide-isomerase A4 |
| Q9D2V7 | Coro7 | 0,501033882 | 61 | 8,44E-05 | 0,000251 319 | Coronin-7 |
| P62748 | Hpcal1 | 0,508021292 | 12 | 0,009855 32 | 0,012287 924 | Hippocalcin-like protein 1 |
| Q99JY9 | Actr3 | 0,513587036 | 45 | 0,000109 65 | 0,000310 289 | Actin-related protein 3 |
| Q9D906 | Atg7 | 0,5136916 | 53 | 0,000438 55 | 0,000967 936 | Ubiquitin-like modifier-activating enzyme ATG7 |
| P46412 | Gpx3 | 0,527495196 | 9 | 4,15E-05 | 0,000135 48 | Glutathione peroxidase 3 |
| P17918 | Pcna | 0,528628063 | 27 | 0,005391 09 | 0,007549 746 | Proliferating cell nuclear antigen |

| | | | | | | |
|---|---|---|---|---|---|---|
| Q9D8V0 | Hm13 | 0,528937158 | 14 | 0,002239 87 | 0,003734 01 | Minor histocompatibility antigen H13 |
| B7FAU9 | Flna | 0,532418525 | 211 | 3,24E-42 | 7,36E-40 | Filamin, alpha |
| O88207 | Col5a1 | 0,536767777 | 19 | 0,047966 42 | 0,041432 742 | Collagen alpha-1(V) chain |
| P22752 | Hist1h2a b | 0,546840584 | 9 | 0,000335 12 | 0,000785 407 | Histone H2A type 1 |
| P70290 | Mpp1 | 0,563112582 | 37 | 0,018762 39 | 0,020119 415 | 55 kDa erythrocyte membrane protein |
| Q62266 | Sprr1a | 0,571330812 | 12 | 0,018715 37 | 0,020100 599 | Cornifin-A |
| Q6ZQI3 | Mlec | 0,572581165 | 27 | 0,004347 05 | 0,006430 998 | Malectin |
| P26231 | Ctnna1 | 0,577088818 | 69 | 5,73E-05 | 0,000183 583 | Catenin alpha-1 |
| P62806 | Hist1h4a | 0,585693706 | 25 | 8,01E-14 | 1,71E-12 | Histone H4 |
| P61161 | Actr2 | 0,587552985 | 34 | 0,000122 52 | 0,000336 933 | Actin-related protein 2 |
| P07901 | Hsp90aa 1 | 0,596019723 | 74 | 0,000192 05 | 0,000481 541 | Heat shock protein HSP 90-alpha |
| P24527 | Lta4h | 0,602210245 | 55 | 6,02E-13 | 1,11E-11 | Leukotriene A-4 hydrolase |
| P43276 | Hist1h1b | 0,611050523 | 19 | 5,89E-12 | 8,73E-11 | Histone H1.5 |
| F8WJE0 | Samhd1 | 0,624243315 | 27 | 0,031735 49 | 0,030102 371 | Deoxynucleoside triphosphate triphosphohydrolase SAMHD1 |
| Q3THE2 | Myl12b | 0,62535957 | 19 | 6,71E-09 | 5,65E-08 | Myosin regulatory light chain 12B |
| P68433 | Hist1h3a | 0,635429257 | 25 | 6,96E-09 | 5,79E-08 | Histone H3.1 |
| Q9WUU7 | Ctsz | 0,648918562 | 21 | 0,025829 08 | 0,025786 649 | Cathepsin Z |
| P47753 | Capza1 | 0,655581627 | 9 | 7,50E-10 | 7,63E-09 | F-actin-capping protein subunit alpha-1 |
| P26041 | Msn | 0,661093985 | 65 | 4,21E-13 | 8,21E-12 | Moesin |
| P62962 | Pfn1 | 0,688742285 | 7 | 9,12E-06 | 3,59E-05 | Profilin-1 |
| P40124 | Cap1 | 0,711077724 | 54 | 3,47E-05 | 0,000116 072 | Adenylyl cyclase-associated protein 1 |
| Q9CPW4 | Arpc5 | 0,713713773 | 19 | 0,000325 19 | 0,000767 414 | Actin-related protein 2/3 complex subunit 5 |
| Q91VE6 | Nifk | 0,7196942 | 22 | 0,011004 78 | 0,013283 648 | MKI67 FHA domain-interacting nucleolar phosphoprotein |
| P08752 | Gnai2 | 0,725601166 | 35 | 1,66E-11 | 2,41E-10 | Guanine nucleotide-binding protein G(i) subunit alpha-2 |
| P63001 | Rac1 | 0,732588081 | 7 | 0,050877 55 | 0,043318 964 | Ras-related C3 botulinum toxin substrate 1 |
| P10853 | Hist1h2bf | 0,735926009 | 11 | 0,054714 29 | 0,045617 541 | Histone H2B type 1-F/J/L |
| Q8BJS4 | Sun2 | 0,753073902 | 49 | 2,13E-16 | 6,60E-15 | SUN domain-containing protein 2 |
| P43277 | Hist1h1d | 0,753794948 | 35 | 7,22E-10 | 7,46E-09 | Histone H1.3 |
| Q62422 | Ostf1 | 0,754142515 | 19 | 0,001616 54 | 0,002871 044 | Osteoclast-stimulating factor 1 |
| P05480 | Src | 0,764998257 | 5 | 0,022042 22 | 0,022708 149 | Neuronal proto-oncogene tyrosine-protein kinase Src |
| A1BN54 | Actn1 | 0,769439067 | 33 | 3,25E-09 | 2,96E-08 | Alpha actinin 1a |
| Q5SS83 | Flot2 | 0,770579893 | 10 | 0,010290 36 | 0,012622 347 | Flotillin 2, isoform CRA_a |
| P43274 | Hist1h1e | 0,772198801 | 9 | 0,000158 3 | 0,000416 834 | Histone H1.4 |
| E9PUF7 | Arhgef1 | 0,773007821 | 15 | 0,000117 73 | 0,000327 735 | Rho guanine nucleotide exchange factor 1 |
| Q3UH59 | Myh10 | 0,785300134 | 30 | 5,93E-05 | 0,000187 28 | Myosin-10 |
| Q9R111 | Gda | 0,786958197 | 43 | 1,06E-06 | 5,82E-06 | Guanine deaminase |
| P97384 | Anxa11 | 0,789008453 | 28 | 1,55E-14 | 3,64E-13 | Annexin A11 |
| P43275 | Hist1h1a | 0,800292939 | 10 | 6,20E-08 | 4,40E-07 | Histone H1.1 |
| Q3U962 | Col5a2 | 0,805348203 | 25 | 0,020911 57 | 0,021806 864 | Collagen alpha-2(V) chain |
| E9Q3Z5 | Svil | 0,81636229 | 13 | 0,018457 8 | 0,019949 635 | Supervillin |
| P52293 | Kpna2 | 0,826563018 | 31 | 2,64E-05 | 9,05E-05 | Importin subunit alpha-1 |

| | | | | | | |
|---|---|---|---|---|---|---|
| P26039 | Tln1 | 0,827388858 | 218 | 1,18E-13 | 2,45E-12 | Talin-1 |
| Q8VCT3 | Rnpep | 0,829402266 | 56 | 0,001511 47 | 0,002748 857 | Aminopeptidase B |
| Q04750 | Top1 | 0,836209033 | 66 | 0,000145 67 | 0,000388 08 | DNA topoisomerase 1 |
| P10107 | Anxa1 | 0,83757079 | 42 | 9,34E-08 | 6,31E-07 | Annexin A1 |
| Q8VDD5 | Myh9 | 0,863282347 | 242 | 2,56E-76 | 1,75E-73 | Myosin-9 |
| A0A0A0M QM2 | Plch1 | 0,892209938 | 4 | 0,014845 66 | 0,016735 111 | Phosphoinositide phospholipase C |
| Q91Z25 | Arpc1b | 0,905090919 | 36 | 1,52E-07 | 9,89E-07 | Actin-related protein 2/3 complex subunit 1B |
| P01887 | B2m | 0,947350666 | 9 | 0,014567 | 0,016448 171 | Beta-2-microglobulin |
| Q3UZ39 | Lrrfip1 | 0,961153858 | 52 | 9,95E-05 | 0,000287 019 | Leucine-rich repeat flightless-interacting protein 1 |
| Q8BT60 | Cpne3 | 0,962426944 | 32 | 0,004632 58 | 0,006765 348 | Copine-3 |
| P15864 | Hist1h1c | 0,974257357 | 12 | 1,45E-06 | 7,61E-06 | Histone H1.2 |
| Q9DCD0 | Pgd | 1,022439575 | 44 | 7,33E-09 | 5,95E-08 | 6-phosphogluconate dehydrogenase, decarboxylating |
| P47791 | Gsr | 1,033420629 | 33 | 0,003460 89 | 0,005233 545 | Glutathione reductase, mitochondrial |
| P30681 | Hmgb2 | 1,043325996 | 22 | 4,89E-08 | 3,59E-07 | High mobility group protein B2 |
| A2APM1 | Cd44 | 1,079687642 | 7 | 6,05E-06 | 2,53E-05 | CD44 antigen |
| P97290 | Serping1 | 1,117261213 | 17 | 3,29E-06 | 1,54E-05 | Plasma protease C1 inhibitor |
| Q8BND5 | Qsox1 | 1,132695551 | 19 | 1,24E-07 | 8,27E-07 | Sulfhydryl oxidase 1 |
| P54116 | Stom | 1,21097851 | 20 | 1,16E-05 | 4,44E-05 | Erythrocyte band 7 integral membrane protein |
| P35175 | Stfa1 | 1,255388754 | 10 | 7,08E-06 | 2,91E-05 | Stefin-1 |
| P07309 | Ttr | 1,257315414 | 11 | 2,25E-06 | 1,12E-05 | Transthyretin |
| Q9CQI6 | Cotl1 | 1,262269134 | 21 | 0,005292 77 | 0,007427 309 | Coactosin-like protein |
| Q9ET01 | Pygl | 1,282596729 | 58 | 8,71E-07 | 4,87E-06 | Glycogen phosphorylase, liver form |
| Q9D154 | Serpinb1 a | 1,292121699 | 30 | 9,33E-15 | 2,27E-13 | Leukocyte elastase inhibitor A |
| P84096 | Rhog | 1,313535081 | 16 | 0,000151 78 | 0,000401 377 | Rho-related GTP-binding protein RhoG |
| P32261 | Serpinc1 | 1,336364034 | 31 | 5,17E-11 | 6,91E-10 | Antithrombin-III |
| P29351 | Ptpn6 | 1,339119327 | 56 | 5,32E-06 | 2,27E-05 | Tyrosine-protein phosphatase non-receptor type 6 |
| Q3U9G9 | Lbr | 1,356729247 | 32 | 1,28E-07 | 8,49E-07 | Lamin-B receptor |
| P04919 | Slc4a1 | 1,360241567 | 38 | 0,000168 75 | 0,000432 665 | Band 3 anion transport protein |
| A0A075B5 P4 | Ighg1 | 1,376113851 | 7 | 0,006038 13 | 0,008170 638 | Ig gamma-1 chain C region secreted form (Fragment) |
| Q80YX1 | Tnc | 1,378076053 | 109 | 0,004999 88 | 0,007092 811 | Tenascin |
| Q8CIH5 | Plcg2 | 1,397616675 | 97 | 0,013365 27 | 0,015475 57 | 1-phosphatidylinositol 4,5-bisphosphate phosphodiesterase gamma-2 |
| P49182 | Serpind1 | 1,405978804 | 10 | 6,32E-08 | 4,45E-07 | Heparin cofactor 2 |
| Q8BL97 | Srsf7 | 1,442801557 | 18 | 0,033892 16 | 0,031707 071 | Serine/arginine-rich splicing factor 7 |
| Q9DBD0 | Ica | 1,448194605 | 36 | 0,008732 81 | 0,011195 451 | Inhibitor of carbonic anhydrase |
| P23953 | Ces1c | 1,462579187 | 28 | 1,72E-10 | 1,98E-09 | Carboxylesterase 1C |
| P07759 | Serpina3 k | 1,463524428 | 16 | 1,22E-20 | 5,19E-19 | Serine protease inhibitor A3K |
| P01644 | NaN | 1,471195549 | 4 | 0,019964 22 | 0,021044 202 | Ig kappa chain V-V region HP R16.7 |
| Q03734 | Serpina3 m | 1,481613548 | 10 | 0,007666 57 | 0,009997 33 | Serine protease inhibitor A3M |
| P21614 | Gc | 1,492069087 | 32 | 6,56E-17 | 2,13E-15 | Vitamin D-binding protein |
| A0A0A0M QA3 | Serpina1 a | 1,498529756 | 22 | 5,46E-17 | 1,86E-15 | Alpha-1-antitrypsin 1-1 |
| Q921M7 | Fam49b | 1,508381208 | 32 | 7,66E-06 | 3,09E-05 | Protein FAM49B |

| | | | | | | |
|---|---|---|---|---|---|---|
| P07724 | Alb | 1,525785341 | 69 | 2,34E-31 | 1,78E-29 | Serum albumin |
| D3YTY9 | Kng1 | 1,528180313 | 19 | 7,31E-11 | 8,90E-10 | Kininogen-1 |
| Q00898 | Serpina1e | 1,539445973 | 11 | 2,64E-05 | 9,05E-05 | Alpha-1-antitrypsin 1-5 |
| E9PZD8 | Cp | 1,544024451 | 63 | 3,27E-10 | 3,66E-09 | Ceruloplasmin |
| P11087 | Col1a1 | 1,545627384 | 68 | 2,06E-12 | 3,27E-11 | Collagen alpha-1(I) chain |
| P06728 | Apoa4 | 1,545955667 | 32 | 0,00039537 | 0,000899698 | Apolipoprotein A-IV |
| Q921I1 | Tf | 1,548364759 | 66 | 1,14E-41 | 1,30E-39 | Serotransferrin |
| Q6YJU1 | Fetub | 1,554167126 | 10 | 1,37E-09 | 1,33E-08 | Fetuin-B |
| P22599 | Serpina1b | 1,575703723 | 7 | 1,22E-06 | 6,63E-06 | Alpha-1-antitrypsin 1-2 |
| P01899 | H2-D1 | 1,580491942 | 25 | 0,00022187 | 0,000542343 | H-2 class I histocompatibility antigen, D-B alpha chain |
| P08226 | Apoe | 1,597444256 | 33 | 5,14E-15 | 1,35E-13 | Apolipoprotein E |
| Q01149 | Col1a2 | 1,607281411 | 59 | 1,47E-08 | 1,16E-07 | Collagen alpha-2(I) chain |
| Q91X72 | Hpx | 1,611279346 | 33 | 8,38E-13 | 1,50E-11 | Hemopexin |
| P29699 | Ahsg | 1,613411511 | 12 | 0,00012729 | 0,000347245 | Alpha-2-HS-glycoprotein |
| Q00897 | Serpina1d | 1,616581486 | 10 | 3,25E-14 | 7,16E-13 | Alpha-1-antitrypsin 1-4 |
| O35639 | Anxa3 | 1,622938307 | 45 | 1,66E-10 | 1,96E-09 | Annexin A3 |
| Q61704 | Itih3 | 1,626384259 | 23 | 0,00131824 | 0,002443043 | Inter-alpha-trypsin inhibitor heavy chain H3 |
| B8JJN0 | Gm20547 | 1,639447972 | 35 | 3,92E-06 | 1,78E-05 | Protein Gm20547 |
| G3X8T9 | Serpina3n | 1,726279905 | 14 | 1,24E-12 | 2,08E-11 | Serine (Or cysteine) peptidase inhibitor, clade A, member 3N, isoform CRA_a |
| Q61233 | Lcp1 | 1,73126668 | 67 | 2,10E-18 | 7,87E-17 | Plastin-2 |
| Q64726 | Azgp1 | 1,735033275 | 15 | 0,0057282 | 0,007908168 | Zinc-alpha-2-glycoprotein |
| Q00623 | Apoa1 | 1,777856263 | 29 | 8,04E-20 | 3,22E-18 | Apolipoprotein A-I |
| Q61599 | Arhgdib | 1,860978444 | 23 | 3,53E-06 | 1,64E-05 | Rho GDP-dissociation inhibitor 2 |
| A0A075B5P2 | Igkc | 1,870412166 | 9 | 0,00610963 | 0,008251019 | Protein Igkc (Fragment) |
| Q8BH35 | C8b | 1,889799149 | 21 | 0,00016849 | 0,000432665 | Complement component C8 beta chain |
| P28665 | Mug1 | 1,951882835 | 79 | 4,49E-22 | 2,04E-20 | Murinoglobulin-1 |
| P06683 | C9 | 1,95413609 | 22 | 0,01412194 | 0,016145735 | Complement component C9 |
| Q9ESB3 | Hrg | 2,015091862 | 17 | 9,37E-05 | 0,000271967 | Histidine-rich glycoprotein |
| D3YW52 | Pzp | 2,032159077 | 61 | 5,86E-32 | 5,00E-30 | Alpha-2-macroglobulin |
| Q8K1B8 | Fermt3 | 2,083782343 | 57 | 2,91E-05 | 9,84E-05 | Fermitin family homolog 3 |
| F8WJ05 | Itih1 | 2,11404846 | 17 | 3,16E-09 | 2,91E-08 | Inter-alpha-trypsin inhibitor heavy chain H1 |
| A2A997 | C8a | 2,147219775 | 12 | 2,56E-05 | 8,86E-05 | Complement component C8 alpha chain |
| A6X935 | Itih4 | 2,149746337 | 23 | 0,0016821 | 0,002949085 | Inter alpha-trypsin inhibitor, heavy chain 4 |
| E9Q8I0 | Cfh | 2,179006921 | 51 | 7,75E-07 | 4,40E-06 | Complement factor H |
| G3X977 | Itih2 | 2,209491753 | 30 | 4,03E-05 | 0,00013222 | Inter-alpha trypsin inhibitor, heavy chain 2 |
| P01029 | C4b | 2,235198055 | 65 | 0,0009701 | 0,001890309 | Complement C4-B |
| Q06890 | Clu | 2,244390732 | 25 | 5,69E-15 | 1,44E-13 | Clusterin |
| A0A075B5P6 | Ighm | 2,2736472 | 15 | 4,54E-06 | 2,00E-05 | Ig mu chain C region (Fragment) |
| A2A6J4 | Lsp1 | 2,351851085 | 11 | 2,93E-06 | 1,41E-05 | Lymphocyte-specific protein 1 |
| P01027 | C3 | 2,417610928 | 119 | 8,66E-42 | 1,18E-39 | Complement C3 |
| Q61247 | Serpinf2 | 2,490031176 | 18 | 1,42E-08 | 1,13E-07 | Alpha-2-antiplasmin |
| Q01339 | Apoh | 2,505299806 | 20 | 4,70E-09 | 4,11E-08 | Beta-2-glycoprotein 1 |

| | | | | | | |
|---|---|---|---|---|---|---|
| **A0A087WR50** | **Fn1** | **2,54572677** | **89** | **1,13E-34** | **1,11E-32** | **Fibronectin** |
| **P19221** | **F2** | **2,551020726** | **25** | **0,00010391** | **0,00029652** | **Prothrombin** |
| **P20918** | **Plg** | **2,55422493** | **52** | **4,79E-15** | **1,31E-13** | **Plasminogen** |
| **O89053** | **Coro1a** | **2,628558871** | **47** | **1,47E-05** | **5,40E-05** | **Coronin-1A** |
| **P29788** | **Vtn** | **3,517039668** | **20** | **1,64E-07** | **1,05E-06** | **Vitronectin** |
| **E9PV24** | **Fga** | **3,914043664** | **37** | **2,19E-18** | **7,87E-17** | **Fibrinogen alpha chain** |
| **P31725** | **S100a9** | **3,919831457** | **12** | **8,87E-07** | **4,92E-06** | **Protein S100-A9** |
| **Q3UER8** | **Fgg** | **4,16295624** | **35** | **8,39E-23** | **4,09E-21** | **Fibrinogen gamma chain** |
| **Q8K0E8** | **Fgb** | **4,184160523** | **27** | **3,34E-23** | **1,90E-21** | **Fibrinogen beta chain** |
| | | | | | | **q-value Threshold** |
| P49443 | Ppm1a | -0,378793396 | 31 | 0,0159513 | 0,017660369 | Protein phosphatase 1A |
| Q61292 | Lamb2 | -0,37688801 | 122 | 6,52E-08 | 4,54E-07 | Laminin subunit beta-2 |
| A2AUD5 | Tpd52l2 | -0,376389952 | 8 | 0,04281293 | 0,038068344 | Tumor protein D54 |
| Q99MQ4 | Aspn | -0,375626495 | 16 | 0,00067126 | 0,001388266 | Asporin |
| Q8BGC4 | Zadh2 | -0,373227921 | 20 | 0,00750688 | 0,009826666 | Zinc-binding alcohol dehydrogenase domain-containing protein 2 |
| Q8C2Q8 | Atp5c1 | -0,371346433 | 21 | 0,00294517 | 0,004628128 | ATP synthase subunit gamma |
| P06801 | Me1 | -0,370596651 | 46 | 1,75E-06 | 8,97E-06 | NADP-dependent malic enzyme |
| Q8BFS6 | Cpped1 | -0,370506661 | 16 | 0,00135969 | 0,002501303 | Serine/threonine-protein phosphatase CPPED1 |
| Q9Z2I9 | Sucla2 | -0,365049701 | 43 | 0,00413186 | 0,006197599 | Succinyl-CoA ligase [ADP-forming] subunit beta, mitochondrial |
| Q9DCT2 | Ndufs3 | -0,364150906 | 31 | 0,02320997 | 0,023625668 | NADH dehydrogenase [ubiquinone] iron-sulfur protein 3, mitochondrial |
| Q8VE70 | Pdcd10 | -0,363780946 | 19 | 0,01856936 | 0,019980599 | Programmed cell death protein 10 |
| Q8BWT1 | Acaa2 | -0,361750683 | 40 | 6,35E-10 | 6,77E-09 | 3-ketoacyl-CoA thiolase, mitochondrial |
| Q9JHW2 | Nit2 | -0,359161155 | 22 | 0,00368851 | 0,005553117 | Omega-amidase NIT2 |
| Q9D6P8 | Calml3 | -0,358250285 | 3 | 0,01110265 | 0,013378108 | Calmodulin-like protein 3 |
| E9PX84 | Dab2 | -0,356875045 | 6 | 0,01330052 | 0,015426794 | Disabled homolog 2 |
| Q3U1J4 | Ddb1 | -0,356764416 | 79 | 0,01286705 | 0,015026243 | DNA damage-binding protein 1 |
| Q9D0K2 | Oxct1 | -0,349635282 | 41 | 0,00027359 | 0,000652412 | Succinyl-CoA:3-ketoacid coenzyme A transferase 1, mitochondrial |
| Q8R1F1 | Fam129b | -0,349391734 | 43 | 8,37E-06 | 3,32E-05 | Niban-like protein 1 |
| F2Z471 | Vdac1 | -0,348483416 | 25 | 0,0096064 | 0,012065494 | Voltage-dependent anion-selective channel protein 1 |
| Q9JJZ2 | Tuba8 | -0,348404954 | 13 | 0,00052146 | 0,001125427 | Tubulin alpha-8 chain |
| E9Q8T1 | Tacc2 | -0,347653804 | 12 | 0,01557726 | 0,017401296 | Transforming acidic coiled-coil-containing protein 2 |
| Q9Z2T6 | Krt85 | -0,347319605 | 16 | 0,00066122 | 0,001374857 | Keratin, type II cuticular Hb5 |
| Q9CQ62 | Decr1 | -0,34693535 | 27 | 0,05993723 | 0,048896159 | 2,4-dienoyl-CoA reductase, mitochondrial |
| P48758 | Cbr1 | -0,346469492 | 31 | 2,24E-07 | 1,41E-06 | Carbonyl reductase [NADPH] 1 |
| A2AS98 | Nckap1 | -0,345447543 | 99 | 0,00525568 | 0,007390466 | Nck-associated protein 1 |
| P00493 | Hprt1 | -0,343647962 | 17 | 0,00440764 | 0,00649246 | Hypoxanthine-guanine phosphoribosyltransferase |
| Q08189 | Tgm3 | -0,343519732 | 7 | 0,00155435 | 0,002804416 | Protein-glutamine gamma-glutamyltransferase E |
| E9Q9C0 | Ablim1 | -0,343135701 | 15 | 0,01968457 | 0,020868433 | Actin-binding LIM protein 1 |
| Q01768 | Nme2 | -0,34252063 | 12 | 0,00941793 | 0,011938714 | Nucleoside diphosphate kinase B |
| P67778 | Phb | -0,341981654 | 32 | 0,00265142 | 0,004284994 | Prohibitin |

| | | | | | | |
|---|---|---|---|---|---|---|
| P30275 | Ckmt1 | -0,341179661 | 43 | 0,00084395 | 0,001695861 | Creatine kinase U-type, mitochondrial |
| Q9CZY3 | Ube2v1 | -0,341071369 | 13 | 0,0031285 | 0,004849168 | Ubiquitin-conjugating enzyme E2 variant 1 |
| P25085 | Il1rn | -0,34036554 | 5 | 0,03265978 | 0,030883873 | Interleukin-1 receptor antagonist protein |
| Q6IMF0 | Krt83 | -0,339780406 | 14 | 0,00035467 | 0,000819951 | Keratin, type II cuticular Hb3 |
| P51885 | Lum | -0,33977593 | 9 | 0,00014503 | 0,000387895 | Lumican |
| D3YYS6 | Mgll | -0,338535153 | 24 | 0,0268154 | 0,026466142 | Monoglyceride lipase |
| P10493 | Nid1 | -0,338475727 | 72 | 0,0006568 | 0,001369845 | Nidogen-1 |
| B1AQY9 | Septin8 | -0,337238002 | 42 | 0,00017881 | 0,000455029 | Septin-8 |
| A2AWT5 | Ubtf | -0,337063652 | 15 | 0,06157732 | 0,049936774 | Nucleolar transcription factor 1 |
| Q9QZ06 | Tollip | -0,333864489 | 16 | 0,00013682 | 0,000370272 | Toll-interacting protein |
| O89051 | Itm2b | -0,332922314 | 21 | 0,01566627 | 0,017401296 | Integral membrane protein 2B |
| O08749 | Dld | -0,330071814 | 39 | 0,00162557 | 0,002879584 | Dihydrolipoyl dehydrogenase, mitochondrial |
| Q9CZU6 | Cs | -0,330058153 | 40 | 0,00901256 | 0,011510423 | Citrate synthase, mitochondrial |
| Q9JHU2 | Palmd | -0,330000041 | 31 | 0,00959704 | 0,012065494 | Palmdelphin |
| Q9R0X4 | Acot9 | -0,3266902 | 46 | 0,02639407 | 0,026163892 | Acyl-coenzyme A thioesterase 9, mitochondrial |
| P68372 | Tubb4b | -0,326358737 | 48 | 1,49E-07 | 9,75E-07 | Tubulin beta-4B chain |
| D3YUM1 | Ndufv1 | -0,325921711 | 39 | 0,00808028 | 0,010476708 | NADH dehydrogenase [ubiquinone] flavoprotein 1, mitochondrial |
| P52196 | Tst | -0,324995413 | 28 | 1,14E-05 | 4,40E-05 | Thiosulfate sulfurtransferase |
| P51859 | Hdgf | -0,324905565 | 32 | 0,00019387 | 0,000484206 | Hepatoma-derived growth factor |
| Q99JY0 | Hadhb | -0,323623144 | 47 | 0,00321078 | 0,004955173 | Trifunctional enzyme subunit beta, mitochondrial |
| P53395 | Dbt | -0,322815863 | 42 | 0,00924579 | 0,011742322 | Lipoamide acyltransferase component of branched-chain alpha-keto acid dehydrogenase complex, mitochondrial |
| Q9JKB3 | Ybx3 | -0,322766094 | 25 | 0,00035273 | 0,000818245 | Y-box-binding protein 3 |
| Q9QZZ6 | Dpt | -0,321748262 | 9 | 0,01346707 | 0,015567017 | Dermatopontin |
| Q8BMF4 | Dlat | -0,320198262 | 38 | 0,02880642 | 0,027906222 | Dihydrolipoyllysine-residue acetyltransferase component of pyruvate dehydrogenase complex, mitochondrial |
| Q9R1P3 | Psmb2 | -0,319151551 | 28 | 0,0012316 | 0,002301233 | Proteasome subunit beta type-2 |
| O35405 | Pld3 | -0,318430594 | 25 | 0,05134006 | 0,043549649 | Phospholipase D3 |
| Q91WD5 | Ndufs2 | -0,316471899 | 44 | 0,04721464 | 0,041019602 | NADH dehydrogenase [ubiquinone] iron-sulfur protein 2, mitochondrial |
| P36993 | Ppm1b | -0,314102508 | 5 | 0,04003247 | 0,036161781 | Protein phosphatase 1B |
| A2A9Q2 | Nrd1 | -0,312818146 | 80 | 1,16E-05 | 4,44E-05 | Nardilysin |
| Q8C1A5 | Thop1 | -0,312797394 | 61 | 6,74E-11 | 8,50E-10 | Thimet oligopeptidase |
| G5E8N5 | Ldha | -0,31265542 | 23 | 0,00054124 | 0,00116443 | L-lactate dehydrogenase |
| P12382 | Pfkl | -0,311127037 | 47 | 0,03643885 | 0,033537513 | ATP-dependent 6-phosphofructokinase, liver type |
| Q99KQ4 | Nampt | -0,311088243 | 45 | 0,05067919 | 0,04320401 | Nicotinamide phosphoribosyltransferase |
| Q920E5 | Fdps | -0,31100204 | 15 | 0,00339518 | 0,005180125 | Farnesyl pyrophosphate synthase |
| Q99K85 | Psat1 | -0,310909609 | 31 | 6,89E-06 | 2,86E-05 | Phosphoserine aminotransferase |
| P04925 | Prnp | -0,310499672 | 19 | 0,0408159 | 0,036722688 | Major prion protein |
| P51660 | Hsd17b4 | -0,307841905 | 66 | 0,00602821 | 0,008170638 | Peroxisomal multifunctional enzyme type 2 |

| | | | | | | |
|---|---|---|---|---|---|---|
| Q9CPV4 | Glod4 | -0,30740156 | 33 | 2,35E-09 | 2,23E-08 | Glyoxalase domain-containing protein 4 |
| A2CEL1 | Mup1 | -0,306844872 | 19 | 0,00339244 | 0,005180125 | Major urinary protein 1 |
| O89086 | Rbm3 | -0,306273454 | 17 | 0,00602015 | 0,008170638 | RNA-binding protein 3 |
| P62311 | Lsm3 | -0,305294305 | 9 | 0,02446581 | 0,024573908 | U6 snRNA-associated Sm-like protein LSm3 |
| Q9QUR6 | Prep | -0,304574401 | 59 | 0,00214584 | 0,00361349 | Prolyl endopeptidase |
| Q61171 | Prdx2 | -0,304467473 | 23 | 5,58E-07 | 3,22E-06 | Peroxiredoxin-2 |
| Q80XN0 | Bdh1 | -0,304124356 | 28 | 0,00599671 | 0,008170638 | D-beta-hydroxybutyrate dehydrogenase, mitochondrial |
| Q62261 | Sptbn1 | -0,303923974 | 314 | 1,25E-12 | 2,08E-11 | Spectrin beta chain, non-erythrocytic 1 |
| E9PYA3 | Hagh | -0,302482437 | 7 | 0,00052138 | 0,001125427 | Hydroxyacylglutathione hydrolase, mitochondrial (Fragment) |
| Q3UF75 | Parva | -0,302082481 | 8 | 0,01609281 | 0,017788162 | Alpha-parvin |
| Q68FH4 | Galk2 | -0,301701781 | 5 | 0,01181614 | 0,014063889 | N-acetylgalactosamine kinase |
| O35640 | Anxa8 | -0,301037296 | 14 | 0,05204863 | 0,043986579 | Annexin A8 |
| Q99KK7 | Dpp3 | -0,300945667 | 61 | 4,60E-07 | 2,71E-06 | Dipeptidyl peptidase 3 |
| P54071 | Idh2 | -0,29595152 | 38 | 0,00112745 | 0,002129981 | Isocitrate dehydrogenase [NADP], mitochondrial |
| Q922Q1 | Mtarc2 | -0,295629618 | 23 | 0,0016563 | 0,002918858 | Mitochondrial amidoxime reducing component 2 |
| P50544 | Acadvl | -0,295320176 | 50 | 9,99E-05 | 0,000287019 | Very long-chain specific acyl-CoA dehydrogenase, mitochondrial |
| Q8BMS1 | Hadha | -0,29353802 | 71 | 7,47E-06 | 3,05E-05 | Trifunctional enzyme subunit alpha, mitochondrial |
| Q8CG72 | Adprhl2 | -0,293535137 | 21 | 0,01014261 | 0,012508613 | Poly(ADP-ribose) glycohydrolase ARH3 |
| P17742 | Ppia | -0,293332428 | 20 | 0,00427793 | 0,006342503 | Peptidyl-prolyl cis-trans isomerase A |
| Q62446 | Fkbp3 | -0,292222984 | 29 | 0,02933135 | 0,028294175 | Peptidyl-prolyl cis-trans isomerase FKBP3 |
| P62965 | Crabp1 | -0,289101419 | 16 | 0,01734292 | 0,018954919 | Cellular retinoic acid-binding protein 1 |
| Q9Z2M7 | Pmm2 | -0,288531102 | 20 | 0,00028691 | 0,000681775 | Phosphomannomutase 2 |
| E9Q8Z6 | Ctnnd1 | -0,287716046 | 22 | 0,00152313 | 0,002762691 | Catenin delta-1 |
| J3QMM7 | Carkd | -0,287672393 | 11 | 0,00277827 | 0,004437425 | ATP-dependent (S)-NAD(P)H-hydrate dehydratase |
| Q9D892 | Itpa | -0,287297014 | 16 | 0,02666442 | 0,026355268 | Inosine triphosphate pyrophosphatase |
| P51881 | Slc25a5 | -0,286843758 | 55 | 0,00078622 | 0,001600599 | ADP/ATP translocase 2 |
| Q9JLF6 | Tgm1 | -0,285643589 | 34 | 0,00087496 | 0,001749911 | Protein-glutamine gamma-glutamyltransferase K |
| P16546 | Sptan1 | -0,285334869 | 363 | 3,50E-30 | 2,17E-28 | Spectrin alpha chain, non-erythrocytic 1 |
| Q8QZT1 | Acat1 | -0,283357267 | 41 | 3,95E-07 | 2,36E-06 | Acetyl-CoA acetyltransferase, mitochondrial |
| Q60597 | Ogdh | -0,281925408 | 94 | 0,00041608 | 0,000939625 | 2-oxoglutarate dehydrogenase, mitochondrial |
| F6TCF9 | Bag1 | -0,281103091 | 17 | 0,0548573 | 0,04565744 | BAG family molecular chaperone regulator 1 |
| Q3UPH1 | Prrc1 | -0,280667486 | 18 | 0,02184323 | 0,022571342 | Protein PRRC1 |
| Q9CZ13 | Uqcrc1 | -0,278467303 | 37 | 0,01402329 | 0,01607376 | Cytochrome b-c1 complex subunit 1, mitochondrial |
| A2AGT5 | Ckap5 | -0,277667355 | 161 | 0,01310345 | 0,01525009 | Cytoskeleton-associated protein 5 |
| P12815 | Pdcd6 | -0,277525644 | 17 | 0,00038835 | 0,000888781 | Programmed cell death protein 6 |
| D3Z041 | Acsl1 | -0,277316709 | 24 | 0,0086969 | 0,011191102 | Long-chain-fatty-acid--CoA ligase 1 |
| Q91VD9 | Ndufs1 | -0,276599775 | 70 | 0,00182638 | 0,003177528 | NADH-ubiquinone oxidoreductase 75 kDa subunit, mitochondrial |
| E9Q7Q3 | Tpm3 | -0,276006227 | 22 | 0,00159379 | 0,002845462 | Tropomyosin alpha-3 chain |

| | | | | | | |
|---|---|---|---|---|---|---|
| D3Z4B2 | Napg | -0,274119227 | 4 | 0,00307171 | 0,004804837 | Gamma-soluble NSF attachment protein (Fragment) |
| Q8BFW7 | Lpp | -0,273588336 | 40 | 0,00060089 | 0,001268757 | Lipoma-preferred partner homolog |
| Q07076 | Anxa7 | -0,27331896 | 32 | 3,13E-06 | 1,49E-05 | Annexin A7 |
| Q9WTI7 | Myo1c | -0,272022945 | 96 | 2,09E-05 | 7,30E-05 | Unconventional myosin-Ic |
| A0A0A6YX18 | Atp6v1h | -0,270738791 | 17 | 0,00067378 | 0,001388266 | V-type proton ATPase subunit H |
| Q7TSV4 | Pgm2 | -0,269347087 | 45 | 0,00953448 | 0,012041693 | Phosphoglucomutase-2 |
| Q6ZQK5 | Acap2 | -0,268725514 | 57 | 0,00093068 | 0,001845134 | Arf-GAP with coiled-coil, ANK repeat and PH domain-containing protein 2 |
| P62843 | Rps15 | -0,266676237 | 16 | 0,01938833 | 0,020628457 | 40S ribosomal protein S15 |
| Q920A5 | Scpep1 | -0,265968853 | 20 | 0,03072851 | 0,029310273 | Retinoid-inducible serine carboxypeptidase |
| B1AT82 | Prpsap1 | -0,265115302 | 23 | 0,05633207 | 0,046511464 | MCG6846, isoform CRA_c |
| G3UWS4 | Ppp2r1b | -0,264599458 | 14 | 0,02938136 | 0,028302387 | Serine/threonine-protein phosphatase 2A 65 kDa regulatory subunit A beta isoform |
| Q9QY23 | Pkp3 | -0,262541357 | 19 | 0,01000161 | 0,012379906 | Plakophilin-3 |
| Q7TQI3 | Otub1 | -0,262161524 | 21 | 0,0253877 | 0,025424984 | Ubiquitin thioesterase OTUB1 |
| P35700 | Prdx1 | -0,261302617 | 34 | 7,22E-05 | 0,000221694 | Peroxiredoxin-1 |
| P14824 | Anxa6 | -0,260623365 | 84 | 0,0408688 | 0,036722688 | Annexin A6 |
| O09005 | Degs1 | -0,260549586 | 8 | 0,00191867 | 0,003304385 | Sphingolipid delta(4)-desaturase DES1 |
| Q91V61 | Sfxn3 | -0,260024926 | 25 | 8,17E-05 | 0,000247776 | Sideroflexin-3 |
| Q9CY58 | Serbp1 | -0,259168068 | 49 | 0,02918319 | 0,028231115 | Plasminogen activator inhibitor 1 RNA-binding protein |
| B1AX58 | Pls3 | -0,257868204 | 55 | 0,00108927 | 0,002075095 | Plastin-3 |
| P63323 | Rps12 | -0,257550681 | 11 | 0,05381439 | 0,044977219 | 40S ribosomal protein S12 |
| Q9DB77 | Uqcrc2 | -0,256612878 | 38 | 0,04130108 | 0,037062286 | Cytochrome b-c1 complex subunit 2, mitochondrial |
| O35643 | Ap1b1 | -0,252632182 | 48 | 0,00118027 | 0,002223612 | AP-1 complex subunit beta-1 |
| D3YXG2 | Nagk | -0,251408531 | 4 | 0,03381498 | 0,031678316 | N-acetyl-D-glucosamine kinase |
| O08547 | Sec22b | -0,246193907 | 23 | 0,0273449 | 0,026833413 | Vesicle-trafficking protein SEC22b |
| Q9CQX2 | Cyb5b | -0,246177459 | 15 | 0,02925486 | 0,028260356 | Cytochrome b5 type B |
| P55264 | Adk | -0,244586307 | 36 | 8,25E-05 | 0,000248138 | Adenosine kinase |
| Q64133 | Maoa | -0,2427807 | 27 | 0,03783384 | 0,034588042 | Amine oxidase [flavin-containing] A |
| P54775 | Psmc4 | -0,242049345 | 51 | 0,00321142 | 0,004955173 | 26S protease regulatory subunit 6B |
| P56480 | Atp5b | -0,241764581 | 48 | 0,0005192 | 0,001125427 | ATP synthase subunit beta, mitochondrial |
| Q61768 | Kif5b | -0,241532123 | 117 | 1,58E-06 | 8,17E-06 | Kinesin-1 heavy chain |
| Q8BGQ7 | Aars | -0,239634556 | 84 | 0,00211364 | 0,003576926 | Alanine--tRNA ligase, cytoplasmic |
| Q9Z1F9 | Uba2 | -0,239513154 | 51 | 0,05907933 | 0,048311878 | SUMO-activating enzyme subunit 2 |
| E9QPD7 | Pcx | -0,239355166 | 90 | 0,01420201 | 0,016196939 | Pyruvate carboxylase |
| P05202 | Got2 | -0,238874548 | 59 | 0,01241458 | 0,014651253 | Aspartate aminotransferase, mitochondrial |
| Q60770 | Stxbp3 | -0,236178923 | 41 | 0,02852947 | 0,027677236 | Syntaxin-binding protein 3 |
| P62196 | Psmc5 | -0,233296132 | 46 | 0,0103843 | 0,01269192 | 26S protease regulatory subunit 8 |
| Q99J36 | Thumpd1 | -0,229733526 | 32 | 0,03791956 | 0,034619997 | THUMP domain-containing protein 1 |
| P48678 | Lmna | -0,229412015 | 93 | 2,20E-15 | 6,53E-14 | Prelamin-A/C |

| | | | | | | |
|---|---|---|---|---|---|---|
| P68369 | Tuba1a | -0,229192838 | 16 | 0,0012963 | 0,00240892 | Tubulin alpha-1A chain |
| Q9Z2Q6 | Septin5 | -0,229090195 | 41 | 0,00213882 | 0,003610589 | Septin-5 |
| P42669 | Pura | -0,226901397 | 21 | 0,00119347 | 0,002242276 | Transcriptional activator protein Pur-alpha |
| P46935 | Nedd4 | -0,226032288 | 63 | 0,00061593 | 0,001296496 | E3 ubiquitin-protein ligase NEDD4 |
| P56399 | Usp5 | -0,223651276 | 78 | 0,01882427 | 0,020154081 | Ubiquitin carboxyl-terminal hydrolase 5 |
| Q9CY27 | Tecr | -0,223228189 | 25 | 0,0135736 | 0,015637155 | Very-long-chain enoyl-CoA reductase |
| P97807 | Fh | -0,223153102 | 37 | 0,0045721 | 0,006691358 | Fumarate hydratase, mitochondrial |
| P00405 | Mtco2 | -0,222824129 | 10 | 0,01052283 | 0,012795451 | Cytochrome c oxidase subunit 2 |
| Q61598 | Gdi2 | -0,222813507 | 56 | 0,00014218 | 0,000381762 | Rab GDP dissociation inhibitor beta |
| P58389 | Ppp2r4 | -0,221970819 | 29 | 0,00421794 | 0,006280867 | Serine/threonine-protein phosphatase 2A activator |
| Q8BIJ6 | Iars2 | -0,212255271 | 60 | 0,00308962 | 0,004821789 | Isoleucine--tRNA ligase, mitochondrial |
| Q9CZD3 | Gars | -0,211362334 | 69 | 5,77E-05 | 0,000183902 | Glycine--tRNA ligase |
| Q99KC8 | Vwa5a | -0,210767346 | 63 | 9,24E-05 | 0,000269367 | von Willebrand factor A domain-containing protein 5A |
| Q8CAQ8 | Immt | -0,210290388 | 97 | 0,01814476 | 0,019673647 | MICOS complex subunit Mic60 |
| Q9JKY5 | Hip1r | -0,208343104 | 98 | 0,00971451 | 0,012178855 | Huntingtin-interacting protein 1-related protein |
| D3Z2Z1 | Clip1 | -0,207923587 | 41 | 0,04537208 | 0,03967149 | CAP-Gly domain-containing linker protein 1 |
| Q9WVA4 | Tagln2 | -0,20749366 | 22 | 0,02085106 | 0,021777068 | Transgelin-2 |
| Q61035 | Hars | -0,207335463 | 53 | 0,00065187 | 0,001363721 | Histidine--tRNA ligase, cytoplasmic |
| P26443 | Glud1 | -0,206271401 | 62 | 5,15E-06 | 2,24E-05 | Glutamate dehydrogenase 1, mitochondrial |
| Q3V117 | Acly | -0,205848978 | 49 | 4,11E-06 | 1,85E-05 | ATP-citrate synthase |
| E9Q1V9 | Camk2d | -0,204353958 | 11 | 0,05942531 | 0,048536598 | Calcium/calmodulin-dependent protein kinase type II subunit delta |
| Q3V188 | Endou | -0,200423328 | 17 | 0,00019524 | 0,000484206 | Poly(U)-specific endoribonuclease |
| P47809 | Map2k4 | -0,199328525 | 35 | 0,04966155 | 0,042495833 | Dual specificity mitogen-activated protein kinase kinase 4 |
| Q9CQT1 | Mri1 | -0,199308749 | 23 | 0,0366297 | 0,033667733 | Methylthioribose-1-phosphate isomerase |
| P97372 | Psme2 | -0,199112645 | 29 | 0,02342207 | 0,023806036 | Proteasome activator complex subunit 2 |
| Q9JHU4 | Dync1h1 | -0,198509967 | 466 | 9,68E-13 | 1,69E-11 | Cytoplasmic dynein 1 heavy chain 1 |
| O88544 | Cops4 | -0,197916014 | 35 | 0,01175235 | 0,014040113 | COP9 signalosome complex subunit 4 |
| Q9CZ44 | Nsfl1c | -0,197859574 | 48 | 0,00224479 | 0,00373401 | NSFL1 cofactor p47 |
| Q9R1T2 | Sae1 | -0,197238726 | 36 | 0,02000203 | 0,021051516 | SUMO-activating enzyme subunit 1 |
| Q99KJ8 | Dctn2 | -0,196532207 | 31 | 8,48E-05 | 0,000251426 | Dynactin subunit 2 |
| E9PXY8 | Usp7 | -0,195751789 | 22 | 0,0408495 | 0,036722688 | Ubiquitin carboxyl-terminal hydrolase |
| Q62465 | Vat1 | -0,193681051 | 28 | 0,00246607 | 0,004023592 | Synaptic vesicle membrane protein VAT-1 homolog |
| Q8R422 | Cd109 | -0,190538561 | 49 | 0,00077342 | 0,001582754 | CD109 antigen |
| Q9DBL7 | Coasy | -0,188341905 | 29 | 0,02242288 | 0,023030727 | Bifunctional coenzyme A synthase |
| Q8R016 | Blmh | -0,18770019 | 45 | 0,04742768 | 0,041152263 | Bleomycin hydrolase |
| Q06138 | Cab39 | -0,185107969 | 31 | 0,01413344 | 0,016145735 | Calcium-binding protein 39 |
| Q8VHX6 | Flnc | -0,184147254 | 171 | 0,00418195 | 0,00625458 | Filamin-C |
| Q9CZ30 | Ola1 | -0,183344283 | 47 | 0,01053709 | 0,012795451 | Obg-like ATPase 1 |

| | | | | | | |
|---|---|---|---|---|---|---|
| P19096 | Fasn | -0,183217682 | 187 | 5,30E-09 | 4,57E-08 | Fatty acid synthase |
| Q9Z2I0 | Letm1 | -0,182054426 | 58 | 0,01191976 | 0,014162502 | LETM1 and EF-hand domain-containing protein 1, mitochondrial |
| P18242 | Ctsd | -0,181836877 | 33 | 0,01437793 | 0,016342911 | Cathepsin D |
| P17426 | Ap2a1 | -0,180297243 | 83 | 0,01565699 | 0,017401296 | AP-2 complex subunit alpha-1 |
| P97429 | Anxa4 | -0,179767495 | 33 | 0,00340579 | 0,005184705 | Annexin A4 |
| E9Q3M3 | Dctn1 | -0,178675745 | 32 | 0,01970568 | 0,020868433 | Dynactin subunit 1 |
| A0A0A0MQA5 | Tuba4a | -0,178603479 | 26 | 0,00014217 | 0,000381762 | Tubulin alpha-4A chain (Fragment) |
| F8VQJ3 | Lamc1 | -0,1784886 | 33 | 0,00240487 | 0,003933147 | Laminin subunit gamma-1 |
| B1B0C7 | Hspg2 | -0,178459628 | 63 | 2,37E-06 | 1,16E-05 | Basement membrane-specific heparan sulfate proteoglycan core protein |
| Q9DBP5 | Cmpk1 | -0,178209223 | 20 | 0,00420789 | 0,006279608 | UMP-CMP kinase |
| Q3U125 | Fam213a | -0,177336113 | 25 | 0,0192202 | 0,020513573 | Redox-regulatory protein FAM213A |
| Q9D1A2 | Cndp2 | -0,176363809 | 50 | 0,00096826 | 0,001890309 | Cytosolic non-specific dipeptidase |
| O35129 | Phb2 | -0,175611115 | 36 | 0,01170999 | 0,014035524 | Prohibitin-2 |
| Q63844 | Mapk3 | -0,173645095 | 33 | 0,02720128 | 0,026749504 | Mitogen-activated protein kinase 3 |
| Q99KD5 | Unc45a | -0,169423068 | 54 | 0,00489182 | 0,007038445 | Protein unc-45 homolog A |
| P51432 | Plcb3 | -0,168589603 | 74 | 0,02690177 | 0,026513013 | 1-phosphatidylinositol 4,5-bisphosphate phosphodiesterase beta-3 |
| P28352 | Apex1 | -0,167712462 | 25 | 0,03012536 | 0,028854657 | DNA-(apurinic or apyrimidinic site) lyase |
| P07214 | Sparc | -0,166365963 | 19 | 0,02650053 | 0,026231292 | SPARC |
| Q9D379 | Ephx1 | -0,165120355 | 47 | 0,02828415 | 0,027517529 | Epoxide hydrolase 1 |
| Q9D8W5 | Psmd12 | -0,164759517 | 39 | 0,01369077 | 0,015745545 | 26S proteasome non-ATPase regulatory subunit 12 |
| Q7TMM9 | Tubb2a | -0,164515379 | 18 | 0,00987358 | 0,012287924 | Tubulin beta-2A chain |
| P24270 | Cat | -0,164476847 | 53 | 0,01745603 | 0,019017592 | Catalase |
| O55234 | Psmb5 | -0,163905059 | 26 | 0,0084354 | 0,010875124 | Proteasome subunit beta type-5 |
| P50247 | Ahcy | -0,162911079 | 49 | 0,03108102 | 0,02960511 | Adenosylhomocysteinase |
| E9PUD2 | Dnm1l | -0,162760577 | 25 | 0,00055957 | 0,001188878 | Dynamin-1-like protein |
| P17427 | Ap2a2 | -0,16240689 | 55 | 0,03368921 | 0,031647443 | AP-2 complex subunit alpha-2 |
| P57780 | Actn4 | -0,161670882 | 110 | 0,00322479 | 0,004964574 | Alpha-actinin-4 |
| Q3UE92 | Xpnpep1 | -0,16070689 | 52 | 0,02428261 | 0,024461945 | X-prolyl aminopeptidase (Aminopeptidase P) 1, soluble, isoform CRA_b |
| O88844 | Idh1 | -0,16013606 | 44 | 0,00023954 | 0,000580838 | Isocitrate dehydrogenase [NADP] cytoplasmic |
| P49586 | Pcyt1a | -0,159219291 | 28 | 0,03553448 | 0,032749344 | Choline-phosphate cytidylyltransferase A |
| Q9D819 | Ppa1 | -0,158880171 | 34 | 0,06027868 | 0,049057353 | Inorganic pyrophosphatase |
| Q9CX86 | Hnrnpa0 | -0,153820621 | 24 | 0,05543283 | 0,045991718 | Heterogeneous nuclear ribonucleoprotein A0 |
| A0JNY3 | Gphn | -0,153541551 | 8 | 0,041444 | 0,037141665 | Gephyrin |
| B2RUJ2 | Erbb2ip | -0,151605295 | 7 | 0,05706127 | 0,046847251 | Erbb2ip protein |
| P26638 | Sars | -0,149725576 | 29 | 0,00331663 | 0,005083015 | Serine--tRNA ligase, cytoplasmic |
| Q9DCN2 | Cyb5r3 | -0,149184317 | 31 | 0,0126128 | 0,014856534 | NADH-cytochrome b5 reductase 3 |
| Q60598 | Cttn | -0,148154345 | 49 | 0,01556188 | 0,017401296 | Src substrate cortactin |

| | | | | | | |
|---|---|---|---|---|---|---|
| Q9WV91 | Ptgfrn | -0,146020184 | 38 | 0,01977871 | 0,020913301 | Prostaglandin F2 receptor negative regulator |
| A2A863 | Itgb4 | -0,143677239 | 95 | 0,00042907 | 0,000953183 | Integrin beta-4 |
| Q8K010 | Oplah | -0,141791781 | 76 | 0,00157692 | 0,00283015 | 5-oxoprolinase |
| P50516 | Atp6v1a | -0,14040299 | 66 | 0,00050853 | 0,001111596 | V-type proton ATPase catalytic subunit A |
| E9PYT3 | Atl3 | -0,139150163 | 17 | 0,00059807 | 0,001266718 | Atlastin-3 |
| P15532 | Nme1 | -0,138743003 | 20 | 0,00706243 | 0,009389041 | Nucleoside diphosphate kinase A |
| P62334 | Psmc6 | -0,138241826 | 35 | 0,0237163 | 0,024033455 | 26S protease regulatory subunit 10B |
| O70400 | Pdlim1 | -0,136218695 | 26 | 8,26E-05 | 0,000248138 | PDZ and LIM domain protein 1 |
| P50518 | Atp6v1e1 | -0,135586325 | 30 | 0,00711432 | 0,009437415 | V-type proton ATPase subunit E 1 |
| Q8CGF7 | Tcerg1 | -0,135263267 | 77 | 0,04388849 | 0,038721795 | Transcription elongation regulator 1 |
| Q9WUM4 | Coro1c | -0,134784102 | 44 | 0,00222463 | 0,003721874 | Coronin-1C |
| Q9WU78 | Pdcd6ip | -0,134753125 | 84 | 0,00712649 | 0,009437415 | Programmed cell death 6-interacting protein |
| Q9D0R2 | Tars | -0,131217719 | 64 | 0,02145795 | 0,022274459 | Threonine--tRNA ligase, cytoplasmic |
| G5E829 | Atp2b1 | -0,128654781 | 81 | 0,02250063 | 0,023075837 | Plasma membrane calcium-transporting ATPase 1 |
| P10922 | H1f0 | -0,125058365 | 15 | 1,04E-05 | 4,02E-05 | Histone H1.0 |
| P63085 | Mapk1 | -0,12338435 | 30 | 0,04463086 | 0,039225456 | Mitogen-activated protein kinase 1 |
| B2RXS4 | Plxnb2 | -0,12275158 | 100 | 0,03995543 | 0,036140058 | Plexin-B2 |
| G3UYD0 | Gtf2i | -0,122685697 | 7 | 0,04799394 | 0,041432742 | General transcription factor II-I |
| Q7TMB8 | Cyfip1 | -0,11741212 | 93 | 0,02297258 | 0,023448994 | Cytoplasmic FMR1-interacting protein 1 |
| P63321 | Rala | -0,117310321 | 14 | 0,01041973 | 0,012712438 | Ras-related protein Ral-A |
| Q99PT1 | Arhgdia | -0,113506703 | 28 | 0,00042732 | 0,000952393 | Rho GDP-dissociation inhibitor 1 |
| Q91YR1 | Twf1 | -0,112390116 | 30 | 0,02008993 | 0,021111452 | Twinfilin-1 |
| Q91VE0 | Slc27a4 | -0,110824699 | 53 | 0,04325855 | 0,038364539 | Long-chain fatty acid transport protein 4 |
| Q6ZQ38 | Cand1 | -0,109549275 | 113 | 0,0446319 | 0,039225456 | Cullin-associated NEDD8-dissociated protein 1 |
| P40336 | Vps26a | -0,109106822 | 35 | 0,00305145 | 0,004784117 | Vacuolar protein sorting-associated protein 26A |
| P63087 | Ppp1cc | -0,107750185 | 29 | 0,0558873 | 0,046255802 | Serine/threonine-protein phosphatase PP1-gamma catalytic subunit |
| Q9DBG3 | Ap2b1 | -0,106901583 | 89 | 0,00096018 | 0,001890309 | AP-2 complex subunit beta |
| Q9JKF1 | Iqgap1 | -0,105252896 | 136 | 0,04706547 | 0,040942152 | Ras GTPase-activating-like protein IQGAP1 |
| Q9DBS1 | Tmem43 | -0,102730528 | 29 | 0,05685207 | 0,04682743 | Transmembrane protein 43 |
| E9Q1S3 | Sec23a | -0,100820415 | 49 | 9,14E-05 | 0,000268678 | Protein transport protein Sec23A |
| Q9CQ65 | Mtap | -0,099752134 | 24 | 0,0056926 | 0,00787653 | S-methyl-5'-thioadenosine phosphorylase |
| Q9ES97 | Rtn3 | -0,099490756 | 75 | 0,03684721 | 0,033822065 | Reticulon-3 |
| P26516 | Psmd7 | -0,097222749 | 26 | 0,02348491 | 0,023834384 | 26S proteasome non-ATPase regulatory subunit 7 |
| Q6P5F9 | Xpo1 | -0,097103277 | 72 | 0,00495876 | 0,007081534 | Exportin-1 |
| Q3THK7 | Gmps | -0,095662876 | 61 | 0,04761309 | 0,041260641 | GMP synthase [glutamine-hydrolyzing] |
| A0A0A0MQA6 | Macf1 | -0,093253601 | 97 | 0,0385029 | 0,035058717 | Microtubule-actin cross-linking factor 1 |
| E9Q390 | Myof | -0,093005878 | 69 | 5,33E-06 | 2,27E-05 | Myoferlin |
| P14069 | S100a6 | -0,092540471 | 10 | 0,04046741 | 0,036506311 | Protein S100-A6 |

| | | | | | | |
|---|---|---|---|---|---|---|
| Q8BH97 | Rcn3 | -0,090852166 | 20 | 0,00173597 | 0,003035715 | Reticulocalbin-3 |
| O08529 | Capn2 | -0,090575163 | 55 | 0,04483342 | 0,039301275 | Calpain-2 catalytic subunit |
| Q3UPL0 | Sec31a | -0,085591626 | 76 | 9,00E-05 | 0,000265695 | Protein transport protein Sec31A |
| Q9EQP2 | Ehd4 | -0,081873936 | 47 | 0,01450407 | 0,016412305 | EH domain-containing protein 4 |
| Q91VI7 | Rnh1 | -0,07662339 | 35 | 0,0438225 | 0,038713659 | Ribonuclease inhibitor |
| Q9QWR8 | Naga | -0,073922525 | 33 | 0,01564123 | 0,017401296 | Alpha-N-acetylgalactosaminidase |
| Q62167 | Ddx3x | -0,073131258 | 49 | 0,04957586 | 0,042489981 | ATP-dependent RNA helicase DDX3X |
| Q9DCL9 | Paics | -0,067571268 | 44 | 0,05226431 | 0,04405965 | Multifunctional protein ADE2 |
| P50396 | Gdi1 | -0,067464185 | 58 | 8,02E-05 | 0,000244111 | Rab GDP dissociation inhibitor alpha |
| Q9D0I9 | Rars | -0,065203117 | 68 | 0,02846392 | 0,02765298 | Arginine--tRNA ligase, cytoplasmic |
| Q922D8 | Mthfd1 | -0,061699362 | 80 | 0,03418963 | 0,031854272 | C-1-tetrahydrofolate synthase, cytoplasmic |
| O89079 | Cope | -0,060260054 | 28 | 0,01100202 | 0,013283648 | Coatomer subunit epsilon |
| P58252 | Eef2 | -0,060258385 | 99 | 0,0001632 | 0,000424823 | Elongation factor 2 |
| P61205 | Arf3 | -0,056788157 | 10 | 0,03293447 | 0,031066817 | ADP-ribosylation factor 3 |
| Q5SXR6 | Cltc | -0,056457262 | 176 | 8,61E-08 | 5,87E-07 | Clathrin heavy chain |
| P14206 | Rpsa | -0,05047768 | 24 | 0,05363023 | 0,0448783 | 40S ribosomal protein SA |
| P97449 | Anpep | -0,049226866 | 47 | 0,00736777 | 0,009700422 | Aminopeptidase N |
| Q5FWK3 | Arhgap1 | -0,041362017 | 43 | 0,01566255 | 0,017401296 | Rho GTPase-activating protein 1 |
| P48036 | Anxa5 | -0,031694063 | 43 | 2,39E-06 | 1,16E-05 | Annexin A5 |
| P16627 | Hspa1l | -0,029211407 | 14 | 0,00109947 | 0,00208307 | Heat shock 70 kDa protein 1-like |
| P28653 | Bgn | -0,028923178 | 17 | 0,05319589 | 0,044624352 | Biglycan |
| Q9QYB5 | Add3 | -0,026619339 | 50 | 0,03314561 | 0,031222802 | Gamma-adducin |
| Q01853 | Vcp | -0,025426332 | 107 | 0,00024017 | 0,000580838 | Transitional endoplasmic reticulum ATPase |
| E9PWE8 | Dpysl3 | -0,014977541 | 26 | 1,14E-09 | 1,13E-08 | Dihydropyrimidinase-related protein 3 |
| Q8BGD9 | Eif4b | -0,014592984 | 71 | 0,05299578 | 0,044511236 | Eukaryotic translation initiation factor 4B |
| O35887 | Calu | -0,012943213 | 37 | 0,05708221 | 0,046847251 | Calumenin |
| P84091 | Ap2m1 | 0,005029122 | 17 | 0,02996436 | 0,028742181 | AP-2 complex subunit mu |
| P26040 | Ezr | 0,005806113 | 54 | 0,02797825 | 0,027297811 | Ezrin |
| P49722 | Psma2 | 0,010469727 | 20 | 0,05423969 | 0,045277194 | Proteasome subunit alpha type-2 |
| P29391 | Ftl1 | 0,012055206 | 22 | 4,18E-06 | 1,86E-05 | Ferritin light chain 1 |
| Q9WVK4 | Ehd1 | 0,012277918 | 37 | 0,04874503 | 0,041974885 | EH domain-containing protein 1 |
| Q60930 | Vdac2 | 0,020783858 | 24 | 0,00161032 | 0,002867471 | Voltage-dependent anion-selective channel protein 2 |
| P62264 | Rps14 | 0,02672855 | 14 | 0,05489604 | 0,04565744 | 40S ribosomal protein S14 |
| P60710 | Actb | 0,030982994 | 61 | 0,0001104 | 0,000311139 | Actin, cytoplasmic 1 |
| Q640N1 | Aebp1 | 0,032054327 | 19 | 0,04947207 | 0,042489981 | Adipocyte enhancer-binding protein 1 |
| P17225 | Ptbp1 | 0,043508345 | 24 | 0,03476358 | 0,032300765 | Polypyrimidine tract-binding protein 1 |
| Q9D2G2 | Dlst | 0,047148335 | 31 | 0,00222658 | 0,003721874 | Dihydrolipoyllysine-residue succinyltransferase component of 2-oxoglutarate dehydrogenase complex, mitochondrial |
| Q9CPS5 | Psmd8 | 0,051388124 | 23 | 0,00466667 | 0,006800582 | 26S proteasome non-ATPase regulatory subunit 8 |

| | | | | | | |
|---|---|---|---|---|---|---|
| P20152 | Vim | 0,054389806 | 93 | 0,002362<br>43 | 0,003882<br>349 | Vimentin |
| Q9D0E1 | Hnrnpm | 0,066551718 | 78 | 0,000830<br>69 | 0,001676<br>133 | Heterogeneous nuclear ribonucleoprotein M |
| Q9QZD9 | Eif3i | 0,072348571 | 29 | 0,042396<br>61 | 0,037796<br>72 | Eukaryotic translation initiation factor 3 subunit I |
| Q3TWW8 | Srsf6 | 0,073430988 | 23 | 0,052988<br>32 | 0,044511<br>236 | Serine/arginine-rich splicing factor 6 |
| P62960 | Ybx1 | 0,074228112 | 33 | 0,011783<br>12 | 0,014049<br>106 | Nuclease-sensitive element-binding protein 1 |
| Q60864 | Stip1 | 0,083744033 | 83 | 0,051069<br>57 | 0,043428<br>235 | Stress-induced-phosphoprotein 1 |
| P62900 | Rpl31 | 0,087504209 | 17 | 0,060258<br>67 | 0,049057<br>353 | 60S ribosomal protein L31 |
| Q61029 | Tmpo | 0,09430702 | 22 | 0,051688<br>9 | 0,043736<br>758 | Lamina-associated polypeptide 2, isoforms beta/delta/epsilon/gamma |
| P42932 | Cct8 | 0,095251162 | 58 | 0,002352<br>36 | 0,003875<br>138 | T-complex protein 1 subunit theta |
| Q6ZQ58 | Larp1 | 0,096735294 | 80 | 0,047026<br>67 | 0,040942<br>152 | La-related protein 1 |
| Q9EST5 | Anp32b | 0,100570087 | 13 | 0,012254<br>09 | 0,014509<br>179 | Acidic leucine-rich nuclear phosphoprotein 32 family member B |
| P62281 | Rps11 | 0,108300524 | 26 | 5,70E-06 | 2,40E-05 | 40S ribosomal protein S11 |
| Q6IRU2 | Tpm4 | 0,108316124 | 38 | 0,021364<br>46 | 0,022211<br>22 | Tropomyosin alpha-4 chain |
| P19324 | Serpinh1 | 0,108922941 | 34 | 0,005409<br>5 | 0,007560<br>003 | Serpin H1 |
| P34022 | Ranbp1 | 0,112639036 | 21 | 0,031372<br>49 | 0,029799<br>495 | Ran-specific GTPase-activating protein |
| Q9JIK9 | Mrps34 | 0,115574915 | 19 | 0,049961<br>94 | 0,042699<br>305 | 28S ribosomal protein S34, mitochondrial |
| Q8R081 | Hnrnpl | 0,11675756 | 49 | 0,000360<br>11 | 0,000829<br>703 | Heterogeneous nuclear ribonucleoprotein L |
| P61358 | Rpl27 | 0,119067517 | 17 | 0,010544<br>05 | 0,012795<br>451 | 60S ribosomal protein L27 |
| P80315 | Cct4 | 0,126951848 | 53 | 0,002309<br>28 | 0,003822<br>648 | T-complex protein 1 subunit delta |
| O55142 | Rpl35a | 0,128183324 | 20 | 0,001837<br>6 | 0,003180<br>824 | 60S ribosomal protein L35a |
| O35381 | Anp32a | 0,134556526 | 22 | 0,002904<br>92 | 0,004586<br>015 | Acidic leucine-rich nuclear phosphoprotein 32 family member A |
| P60843 | Eif4a1 | 0,13719144 | 26 | 0,049592<br>41 | 0,042489<br>981 | Eukaryotic initiation factor 4A-I |
| Q80WS3 | Fbll1 | 0,142731822 | 24 | 0,021541<br>12 | 0,022292<br>932 | rRNA/tRNA 2'-O-methyltransferase fibrillarin-like protein 1 |
| Q9QXS1 | Plec | 0,144870095 | 569 | 0,001050<br>52 | 0,002016<br>403 | Plectin |
| Q99MR6 | Srrt | 0,145388296 | 66 | 0,022913<br>79 | 0,023429<br>096 | Serrate RNA effector molecule homolog |
| Q9D8E6 | Rpl4 | 0,145769307 | 42 | 1,81E-08 | 1,41E-07 | 60S ribosomal protein L4 |
| Q9D883 | U2af1 | 0,148447347 | 16 | 0,058987<br>87 | 0,048294<br>993 | Splicing factor U2AF 35 kDa subunit |
| Q3THS6 | Mat2a | 0,148564711 | 32 | 0,019514<br>78 | 0,020730<br>655 | S-adenosylmethionine synthase isoform type-2 |
| P70372 | Elavl1 | 0,150337951 | 26 | 0,000100<br>16 | 0,000287<br>019 | ELAV-like protein 1 |
| Q6ZWV7 | Rpl35 | 0,151072021 | 9 | 0,001837<br>54 | 0,003180<br>824 | 60S ribosomal protein L35 |
| P47962 | Rpl5 | 0,153150536 | 29 | 1,20E-05 | 4,54E-05 | 60S ribosomal protein L5 |
| O09167 | Rpl21 | 0,155758067 | 17 | 0,014992<br>27 | 0,016844<br>696 | 60S ribosomal protein L21 |
| Q6A068 | Cdc5l | 0,160835575 | 68 | 0,037375<br>62 | 0,034214<br>995 | Cell division cycle 5-like protein |
| P57776 | Eef1d | 0,161145491 | 23 | 0,025563<br>72 | 0,025563<br>721 | Elongation factor 1-delta |
| P63017 | Hspa8 | 0,16403007 | 87 | 0,030610<br>09 | 0,029238<br>214 | Heat shock cognate 71 kDa protein |
| Q60972 | Rbbp4 | 0,16434101 | 22 | 0,042734<br>02 | 0,038047<br>783 | Histone-binding protein RBBP4 |
| P80318 | Cct3 | 0,165174595 | 63 | 0,000329<br>49 | 0,000774<br>864 | T-complex protein 1 subunit gamma |
| Q5SQB0 | Npm1 | 0,167140177 | 13 | 0,051209<br>7 | 0,043493<br>17 | Nucleophosmin |
| Q62159 | Rhoc | 0,167530634 | 20 | 0,045947<br>56 | 0,040123<br>221 | Rho-related GTP-binding protein RhoC |

| | | | | | | |
|---|---|---|---|---|---|---|
| P62918 | Rpl8 | 0,16868394 | 37 | 6,65E-05 | 0,000207903 | 60S ribosomal protein L8 |
| Q60902 | Eps15l1 | 0,172479433 | 76 | 0,01634649 | 0,018010192 | Epidermal growth factor receptor substrate 15-like 1 |
| P47911 | Rpl6 | 0,177071136 | 40 | 0,01745499 | 0,019017592 | 60S ribosomal protein L6 |
| Q9DBG5 | Plin3 | 0,17773509 | 31 | 0,00282991 | 0,004498832 | Perilipin-3 |
| Q80X90 | Flnb | 0,179705727 | 193 | 0,00344559 | 0,005221989 | Filamin-B |
| P41105 | Rpl28 | 0,180374934 | 20 | 0,00132426 | 0,002447547 | 60S ribosomal protein L28 |
| Q8VIJ6 | Sfpq | 0,183367514 | 64 | 0,00067183 | 0,001388266 | Splicing factor, proline- and glutamine-rich |
| E9Q8L9 | Rab11fip1 | 0,188893012 | 17 | 0,02600438 | 0,025887979 | Rab11 family-interacting protein 1 |
| Q9Z2X1 | Hnrnpf | 0,191044399 | 32 | 0,00524844 | 0,007390466 | Heterogeneous nuclear ribonucleoprotein F |
| F6RJ39 | Acin1 | 0,196556106 | 15 | 0,00191309 | 0,003303103 | Apoptotic chromatin condensation inducer in the nucleus (Fragment) |
| Q62186 | Ssr4 | 0,206087109 | 9 | 0,01929748 | 0,020563879 | Translocon-associated protein subunit delta |
| Q62093 | Srsf2 | 0,206386097 | 21 | 0,00973656 | 0,012184104 | Serine/arginine-rich splicing factor 2 |
| P62242 | Rps8 | 0,206660531 | 23 | 0,00625903 | 0,008436081 | 40S ribosomal protein S8 |
| Q9JKX6 | Nudt5 | 0,210622556 | 17 | 0,01007135 | 0,012443224 | ADP-sugar pyrophosphatase |
| H7BX95 | Srsf1 | 0,213878324 | 37 | 0,02300202 | 0,023448994 | Serine/arginine-rich-splicing factor 1 |
| Q9CQN1 | Trap1 | 0,215423362 | 63 | 0,03529047 | 0,032646006 | Heat shock protein 75 kDa, mitochondrial |
| P35276 | Rab3d | 0,215596255 | 12 | 0,00769342 | 0,010013188 | Ras-related protein Rab-3D |
| Q9DB34 | Chmp2a | 0,216346096 | 20 | 0,02421099 | 0,024461945 | Charged multivesicular body protein 2a |
| D3YXK2 | Safb | 0,216812567 | 12 | 0,00026211 | 0,000629428 | Scaffold attachment factor B1 |
| P58021 | Tm9sf2 | 0,218093209 | 23 | 0,00959408 | 0,012065494 | Transmembrane 9 superfamily member 2 |
| Q61033 | Tmpo | 0,222757483 | 56 | 0,00494655 | 0,007081534 | Lamina-associated polypeptide 2, isoforms alpha/zeta |
| Q05D44 | Eif5b | 0,226858662 | 101 | 0,00815547 | 0,01055414 | Eukaryotic translation initiation factor 5B |
| Q9D1R9 | Rpl34 | 0,231124644 | 11 | 0,00018927 | 0,000476321 | 60S ribosomal protein L34 |
| P35282 | Rab21 | 0,231696578 | 20 | 0,03502769 | 0,032501882 | Ras-related protein Rab-21 |
| O70251 | Eef1b | 0,23229008 | 18 | 0,0048794 | 0,00703541 | Elongation factor 1-beta |
| P14733 | Lmnb1 | 0,23441827 | 82 | 4,63E-13 | 8,77E-12 | Lamin-B1 |
| P0C0S6 | H2afz | 0,237685386 | 7 | 0,02152755 | 0,022292932 | Histone H2A.Z |
| P27659 | Rpl3 | 0,242255233 | 45 | 0,00577181 | 0,007952271 | 60S ribosomal protein L3 |
| Q91ZW3 | Smarca5 | 0,243462974 | 80 | 0,03462489 | 0,032215787 | SWI/SNF-related matrix-associated actin-dependent regulator of chromatin subfamily A member 5 |
| Q62376 | Snrnp70 | 0,250050857 | 34 | 0,00096955 | 0,001890309 | U1 small nuclear ribonucleoprotein 70 kDa |
| Q8BXZ1 | Tmx3 | 0,252219194 | 28 | 0,00055227 | 0,001177748 | Protein disulfide-isomerase TMX3 |
| J3QK23 | Gm9825 | 0,256921431 | 6 | 0,0449132 | 0,039320671 | Protein Gm9825 |
| P47963 | Rpl13 | 0,26166171 | 29 | 0,0043859 | 0,006474426 | 60S ribosomal protein L13 |
| P12970 | Rpl7a | 0,262384439 | 34 | 2,23E-08 | 1,69E-07 | 60S ribosomal protein L7a |
| Q6ZWN5 | Rps9 | 0,263214084 | 27 | 1,28E-05 | 4,81E-05 | 40S ribosomal protein S9 |
| F6QL70 | Gm17669 | 0,264535126 | 14 | 0,0028409 | 0,004505799 | Protein Gm17669 |
| Q3UID0 | Smarcc2 | 0,268809029 | 75 | 0,00597533 | 0,008170638 | SWI/SNF complex subunit SMARCC2 |
| O08795 | Prkcsh | 0,271061839 | 32 | 0,00653747 | 0,008776687 | Glucosidase 2 subunit beta |

| | | | | | | |
|---|---|---|---|---|---|---|
| A2A8V8 | Srrm1 | 0,274264807 | 3 | 0,019024 96 | 0,020337 027 | Serine/arginine repetitive matrix protein 1 |
| Q6URW6 | Myh14 | 0,27557906 | 165 | 0,000296 93 | 0,000703 149 | Myosin-14 |
| Q8VI75 | Ipo4 | 0,276233147 | 55 | 0,016480 23 | 0,018128 255 | Importin-4 |
| P14115 | Rpl27a | 0,277613444 | 14 | 0,000427 02 | 0,000952 393 | 60S ribosomal protein L27a |
| P40142 | Tkt | 0,277931127 | 63 | 0,000795 85 | 0,001615 377 | Transketolase |
| Q9Z1Q5 | Clic1 | 0,293430903 | 28 | 0,000161 41 | 0,000421 761 | Chloride intracellular channel protein 1 |
| P99027 | Rplp2 | 0,295218703 | 13 | 0,043374 24 | 0,038417 182 | 60S acidic ribosomal protein P2 |
| Q921Y0 | Mob1a | 0,296067374 | 12 | 0,013067 65 | 0,015234 425 | MOB kinase activator 1A |
| O54988 | Slk | 0,297040879 | 90 | 0,015249 57 | 0,017105 608 | STE20-like serine/threonine-protein kinase |
| Q5SUR0 | Pfas | 0,298188533 | 84 | 0,000438 1 | 0,000967 936 | Phosphoribosylformylglycinamidine synthase |
| Q9ERG0 | Lima1 | 0,302609406 | 74 | 0,033208 75 | 0,031239 131 | LIM domain and actin-binding protein 1 |
| Q9CX34 | Sugt1 | 0,305620879 | 36 | 0,018574 34 | 0,019980 599 | Protein SGT1 homolog |
| Q9DCA5 | Brix1 | 0,310085416 | 27 | 0,002484 97 | 0,004044 756 | Ribosome biogenesis protein BRX1 homolog |
| Q9D8B3 | Chmp4b | 0,314477164 | 23 | 5,87E-05 | 0,000186 09 | Charged multivesicular body protein 4b |
| A2A547 | Rpl19 | 0,322716793 | 24 | 1,09E-06 | 5,96E-06 | Ribosomal protein L19 |
| Q6P5E4 | Uggt1 | 0,322773147 | 107 | 0,024674 11 | 0,024746 677 | UDP-glucose:glycoprotein glucosyltransferase 1 |
| P47757 | Capzb | 0,323085491 | 34 | 9,21E-05 | 0,000269 367 | F-actin-capping protein subunit beta |
| A2AMW0 | Capzb | 0,325398742 | 6 | 0,005952 41 | 0,008168 099 | Capping protein (Actin filament) muscle Z-line, beta, isoform CRA_a |
| P39749 | Fen1 | 0,329945277 | 19 | 0,007323 26 | 0,009679 189 | Flap endonuclease 1 |
| Q3U7R1 | Esyt1 | 0,331924412 | 71 | 0,024277 12 | 0,024461 945 | Extended synaptotagmin-1 |
| P21981 | Tgm2 | 0,333624382 | 58 | 0,027641 | 0,027029 512 | Protein-glutamine gamma-glutamyltransferase 2 |
| P82198 | Tgfbi | 0,337362532 | 32 | 0,000966 59 | 0,001890 309 | Transforming growth factor-beta-induced protein ig-h3 |
| Q3U741 | Ddx17 | 0,340240608 | 13 | 0,001052 55 | 0,002016 403 | DEAD (Asp-Glu-Ala-Asp) box polypeptide 17, isoform CRA_a |
| P27773 | Pdia3 | 0,343982547 | 61 | 0,010169 3 | 0,012518 884 | Protein disulfide-isomerase A3 |
| P23198 | Cbx3 | 0,344237116 | 12 | 0,001964 2 | 0,003357 36 | Chromobox protein homolog 3 |
| P68510 | Ywhah | 0,352306971 | 28 | 7,83E-07 | 4,42E-06 | 14-3-3 protein eta |
| P62911 | Rpl32 | 0,352564851 | 11 | 0,002702 98 | 0,004347 723 | 60S ribosomal protein L32 |
| P35980 | Rpl18 | 0,36286777 | 19 | 3,26E-06 | 1,53E-05 | 60S ribosomal protein L18 |
| Q60865 | Caprin1 | 0,363300893 | 36 | 0,016511 08 | 0,018132 938 | Caprin-1 |
| P12388 | Serpinb2 | 0,364204862 | 26 | 0,000408 93 | 0,000926 556 | Plasminogen activator inhibitor 2, macrophage |
| P13020 | Gsn | 0,365264462 | 32 | 0,002329 75 | 0,003847 195 | Gelsolin |
| Q9JKR6 | Hyou1 | 0,370023351 | 85 | 0,000845 44 | 0,001695 861 | Hypoxia up-regulated protein 1 |
| Q5EG47 | Prkaa1 | 0,373306047 | 29 | 0,033786 45 | 0,031678 316 | 5'-AMP-activated protein kinase catalytic subunit alpha-1 |
| Q9JIK5 | Ddx21 | 0,3752579 | 80 | 0,000200 38 | 0,000495 153 | Nucleolar RNA helicase 2 |
| P14211 | Calr | 0,378516152 | 60 | 1,24E-06 | 6,67E-06 | Calreticulin |
| P20060 | Hexb | 0,379597556 | 42 | 0,004134 76 | 0,006197 599 | Beta-hexosaminidase subunit beta |

### Supplement Table 8. Antibodies

| Antibody | Working dilution | Fluorophore | Supplier (Cat. No.) |
|---|---|---|---|
| **Rabbit MMP9 (primary)** | 1:100 | / | Sigma-Aldrich (AB19016) |
| **Rabbit ELANE (primary)** | 1:200 | / | Invitrogen (PA5-115648) |
| **Mouse CD66b (primary)** | 1:200 | Alexa Fluor 594 | BioLegend (392908) |
| **Goat Rabbit AF488 (secondary)** | 1:250 | Alexa Fluor Plus 488 | Invitrogen (A32731) |

Figure S1

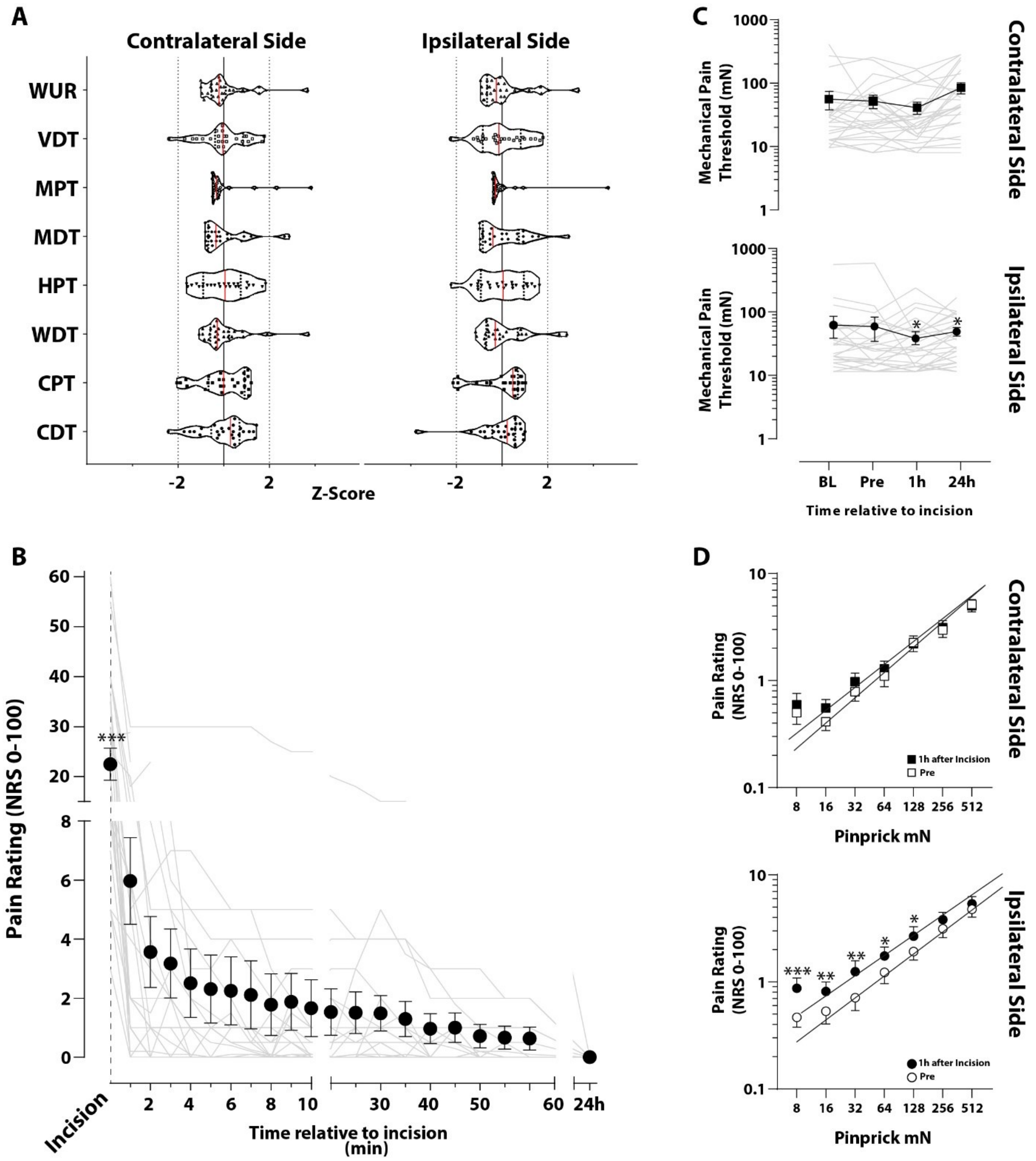


**Figure S1. Incision injury is induced ongoing pain and alterations in mechanical thresholds**

(A) Detection of somatosensory perception to natural stimuli (thermal and mechanical) was performed by a complete quantitative sensory testing (QST) battery on both forearms. Contra- and ipsilateral sides were determined randomly. Raw QST-data (table S3) were transformed to standard normal distribution (Z-transformation) to achieve the QST parameters independent of their physical dimension. Z-scores above '0' indicate a gain of function (more sensitive), below a loss of function (less sensitive). Values above or below twofold standard deviation (SD, dotted line) indicate pathological QST-scores. N=26, single values and median (red line) with interquartile range (dotted line) in violin plots. Aberrations: CDT, cold detection threshold; CPT, cold pain threshold; HPT, heat pain threshold; MDT, mechanical detection threshold; MPS, mechanical pain sensitivity; MPT, mechanical pain threshold; PPT, pressure pain threshold; TSL, thermal sensory limen; VDT, vibration detection threshold; WDT, warmth detection threshold; WUR, wind-up ratio. (B) Pain related to incision on the numeric rating scale (NRS, 0;100) was maximal during skin incision and rapidly decreased in the first 10 min and completely absent after 24h. Individual profiles of each volunteer were plotted in gray. N=26, Mean with SEM, Holm-Sidak multiple comparison test, ***P<0.001. (C) Time course of mechanical pain threshold (MPT) before and post-incision. MPT was determined in milli Newton (mN) on both sides. Upon incision, MPT was significantly decreased 1h and 24 h (POD1) post-incision. Individual profiles of each volunteer were plotted in gray. N=26, Mean with SEM, Holm-Sidak multiple comparison test, *P<0.05. (D) Stimulus-response (S/R-) function for suprathreshold stimuli by pinprick probes to detect mechanical pain sensitivity (MPS). Incision induced a significant left shift compared to baseline. Each data point represents average log-transformed pain ratings on a 0 to 100 numeric rating scale. ***P <0.001, **P<0.01, *P<0.05, LSD–post hoc test.

Figure S2

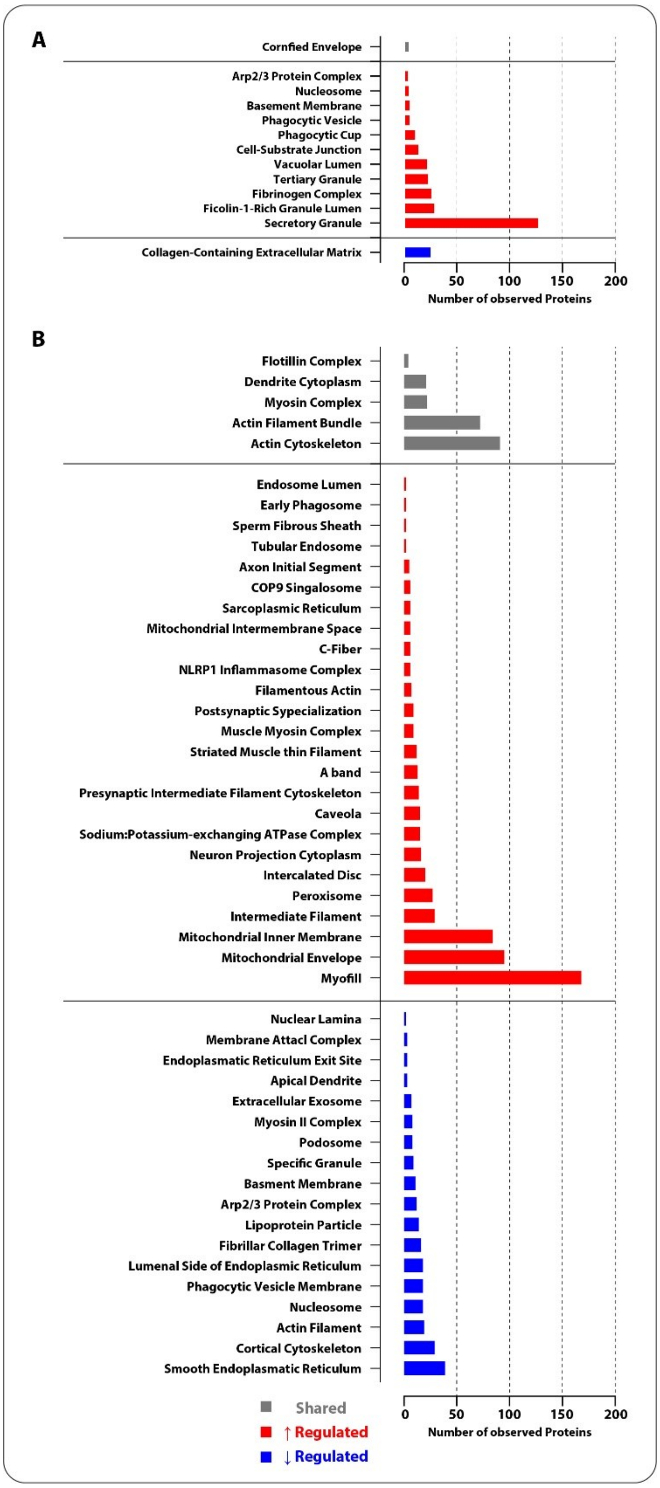


**Figure S2. Predicted localization of proteins in cell compartments**

(A) Using GO_CC analysis, enumeration of predicted cell compartments (CC) of protein in humans. Grey, shared; red, upregulated; blue, downregulated (B) Using GO_CC analysis, enumeration of predicted cell compartments (CC) of protein in mice. Grey, shared; red, upregulated; blue, downregulated

Figure S3

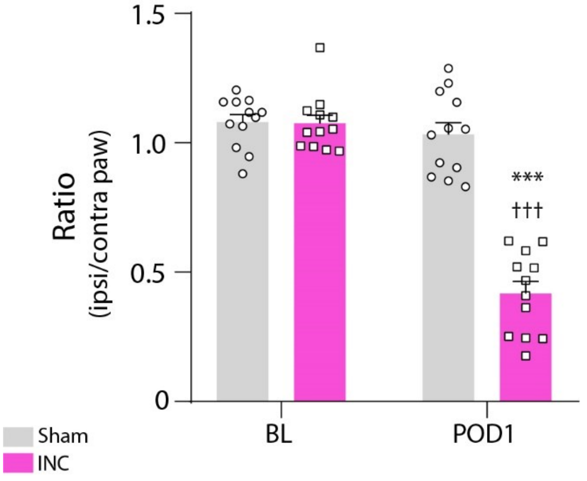


**Figure S3. Non-evoked pain-related behavior in the mouse after plantar incision**

Upon plantar incision, non-evoked pain-related behavior (NEP) was assessed 24h post-incision (INC) (post-operative day 1, POD1) compared to baseline (BL) and sham-treated mice. NEP was represented by the significant decrease in the ratio of the print area of incised vs. contralateral hindpaw at 24 h post-incision (*43*). Sham group displayed unchanged in print areas. NEP was significantly increased (decreased print area) on POD1 to BL (***P<0.001) and sham (†††P<0.001). Data are expressed as mean (bar) and single values (mean ± SEM). Statistics were performed by two-way ANOVA followed by Dunnett's post-hoc test. P-Values: ***P<.001 vs. baseline (BL), †††P<.001 vs. sham.

Figure S4

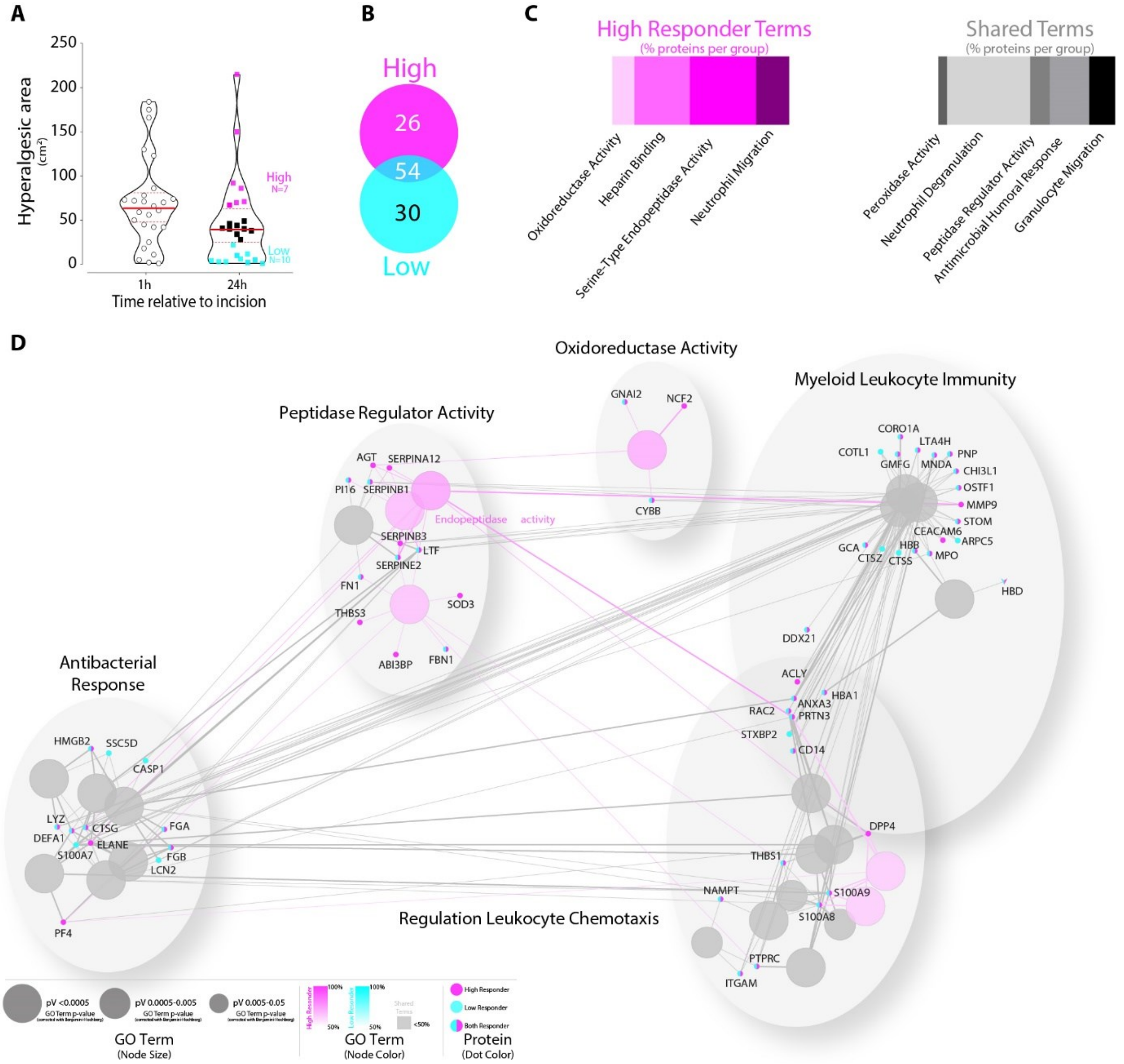


**Figure S4. Dominance of High-responder enriched term-term interaction networks post incision**

(A) A hyperalgesic area developed around 24h post-incision. The total cohort was phenotyped into High (n=7, magenta), Low Responders (n=10, cyan), and Undefined volunteers (n=9, black). Dimension of hyperalgesic area ($cm^2$) at 24h post-incision expresses as individual values in violin plots. HA24 categorization was determined using mean and SD (mean 45.04 $cm^2$ ± 49.62, figure S4A). (B) Twenty-six proteins that were uniquely regulated in High responders, 30 proteins for Low responders, and 54 overlapping proteins were identified. (C) GO-analysis revealed annotations of regulated proteins for High Responders (magenta shades). Shared terms belong to both phenotyping groups (grey shades). In contrast, regulated proteins of the Low responders could not be significantly annotated to distinct pathways. (D) Enriched term-term interaction (TTI) network analysis with associated proteins was functionally grouped by ClueGO (v2.5.8) as functional clusters (AutoAnnonate 1.3). Significantly (p-values ≤ 0.05) enriched GO terms visualized in a functionally grouped network that reflects the terms' relationships. Each node represents a molecular function or immune system process. Node colors associates with phenotyping (High Responder in magenta shades; Low Responder in cyan shades). Grey shades reflect shared terms, which belong to both phenotypes. Proteins display as dots in responder colors (magenta= High Responder, cyan= Low Responder, color shared= phenotype independent). The degree of connectivity between terms (kappa score= 0.3) was used to define functional groups. TTI was represented edges between nodes and dots. The size of nodes reflects the significant enrichment of the terms.

Figure S5

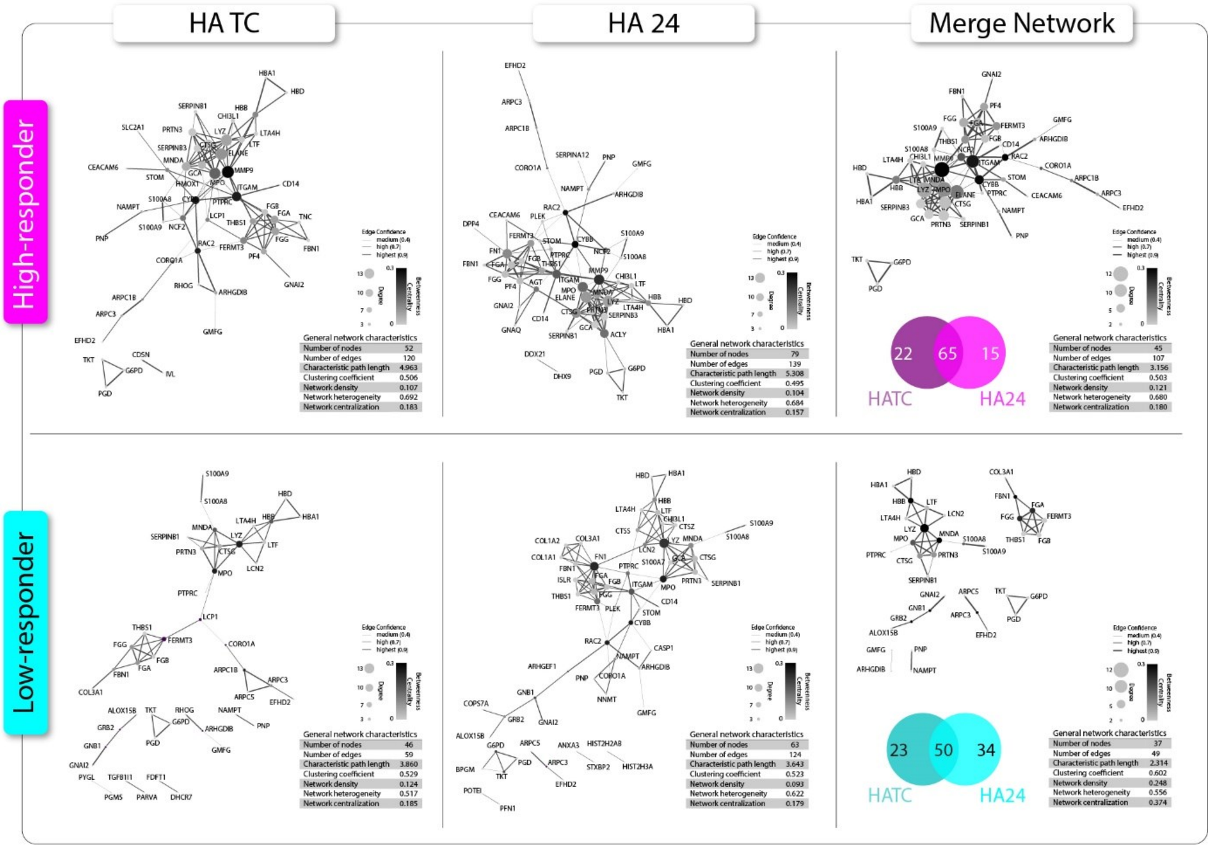


**Figure S5. Skin incision induces distinct phenotypes protein-protein interactions network topology**

Enriched protein-protein interaction (PPI) network topologies of incision-induced phenotypes were analyzed (FDR < 0.05; assessed via the web-based interface STRING) by stringAPP in Cytoscape (3.8.2). Each network displays the specific PPI for pain phenotypes, hyperalgesic area time course (HATC), and hyperalgesic area 24h post-incision (HA24) in both responders (magenta, High Responder; cyan, Low Responder). Merge networks were created within High and Low responders, respectively, representing the pain phenotype PPI networks. Nodes represent a single protein, including degree (size) and betweenness centrality (color). PPI reflects by edges (thickness represents the confidence).

Figure S6

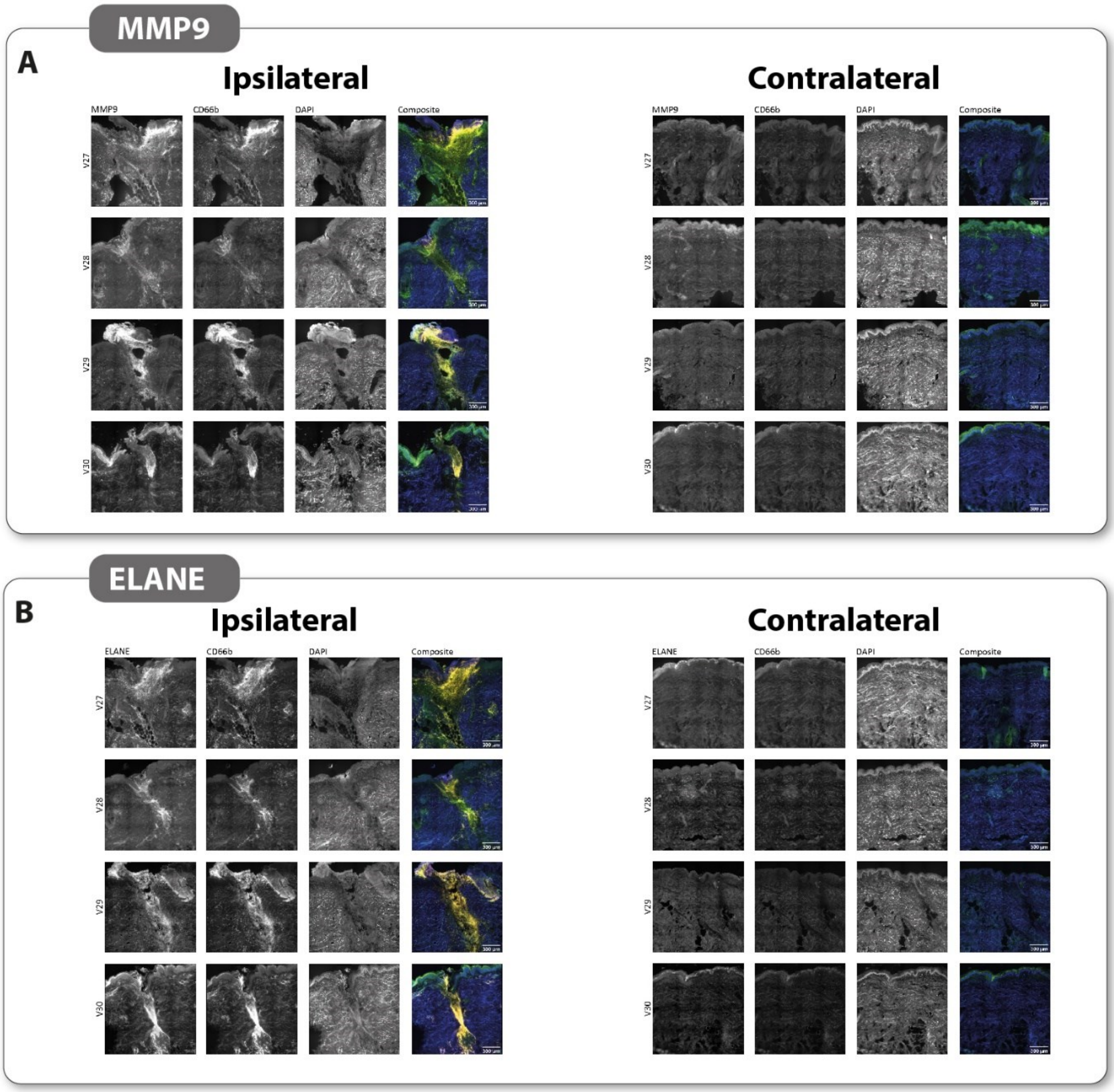


**Figure S6. Individual staining of MMP9 (A) and ELANE (B) in human incised skin of four volunteers**

Scale bar = 500 µm (Composite overview), 200 µm (Composite), 50 µm (Inset). MMP9, matrix metalloproteinase 9; ELANE, neutrophil elastase; DAPI, 4′,6-diamidino-2-phenylindole.
